# Proteome-wide reanalysis identifies coverage and error-rate limits on acquired lysines in cancer

**DOI:** 10.64898/2026.11.09.765507

**Authors:** Rajib Molla, Sona Tiwari, Ayan Mukherjee, Md Khaled Hossain, Mikail E. Abbasov

## Abstract

Missense mutation in cancer creates acquired lysines, residues present in tumor cells and absent from normal cells in the same patient. The class holds 295,411 sites across 20,365 genes and falls on driver hotspots more often than other substitutions (odds ratio 1.68). Yet researching 42 proteomic experiments with a mutation-aware database recovers the mutant peptide in 0.63% of 8,470 site-in-cell-line observations, because a new lysine is a new cleavage site and the peptide it creates runs 6 residues against 13 for the wild type. Reported error rates do not hold. Burden-matched entrapment puts the realized false-discovery proportion at 4.70% before score filtering, against 1% reported. Measured by competitive activity-based protein profiling across four cell lines, ligandability reproduces at the site level, no compound descriptor tested predicts it, and structure predicts it poorly (area under the curve 0.617). Acquired lysines are structurally ordinary, so the barrier is detection rather than chemistry.

**Significance:** A covalent drug is written around the residue it modifies, so the most valuable residue is one that exists in the diseased cell and nowhere else. Cancer supplies such residues by mutation, and the argument has already been made for cysteine, where a mutation-created thiol became a drug target and then a drugged one. Lysine has not been counted the same way, although mutation creates more of them. This work counts the class and then asks the question that decides whether it can be worked on: has anyone been able to see it? Re-searching 42 published proteomes with a mutation-aware database recovers the mutant peptide at well under 1% of the opportunities, and more than nine in ten are never observed in any form. The reason is specific rather than general. A new lysine is a new cleavage site, so the peptide a search must find is not the peptide it would have looked for, and the error control that would license the few identifications it does make has been calibrated over whole searches rather than within this atypical subset of one. Measured directly, with entrapment sequences built to be indistinguishable from their targets by mass and length, the realized error rate at a reported 1% filter is nearly fivefold higher, and a score margin restores it. Set against what three chemoproteomic screens show about which lysines compounds engage, nothing in the chemistry of these sites explains their absence from the record: they sit within the ordinary range of the lysines this chemistry already reaches. The barrier is detection, which is the tractable kind of problem. For anyone designing lysine-directed chemistry, the practical consequence is that this target class has to be measured before it can be prioritized, and the measurements that would do it are named here.

**Graphical abstract:** 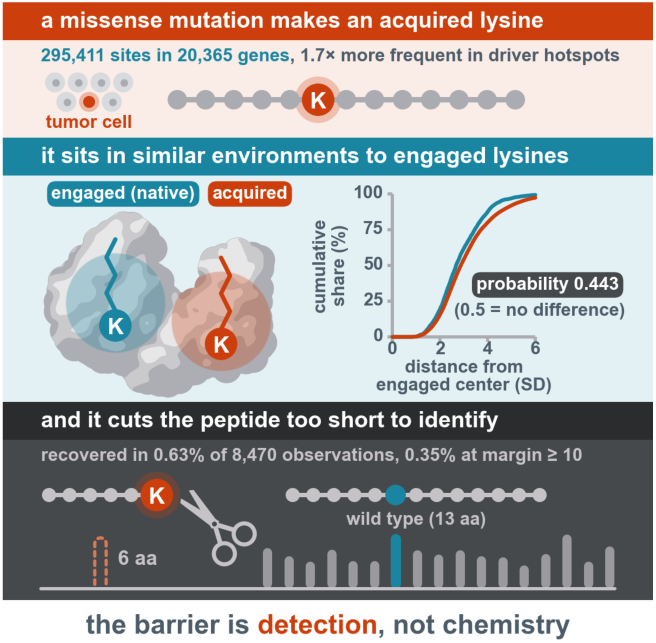

## Introduction

Covalent chemistry directed at lysine has reached the point where the limiting question is no longer whether a lysine can be engaged but which one to engage. Reactivity and ligandability vary enormously between sites^1^, and three subsequent screens mapped thousands of ligandable lysines with chemically distinct probes^2–4^. Newer aminophilic chemistry has enlarged that set again, quantifying over 7,000 modifiable lysines on its own^5^. None of that work supplies a principled way to choose one. Selectivity in a covalent campaign is usually argued from the chemistry of the molecule; the alternative is to choose a residue that exists only in the diseased cell.

Cancer supplies residues found only in diseased cells. A residue created by missense mutation is present in tumor cells and absent from every normal cell in the same patient, which is the logic that made KRAS G12C tractable^6^. That argument has been generalized for cysteine: acquired cysteines have been cataloged across cancer genomes and pursued chemoproteomically as a target class^7^, and the experimentally observed cysteines are kept as a curated community repository carrying functional, druggability and genetic-variant annotation^8^. Lysine has not been counted the same way, although mutation creates more of them, 295,411 acquired lysines against 211,386 acquired cysteines in the union of three variant catalogs. On net flux the order reverses, because cysteine is rarely lost and lysine is not; neither residue is exceptional, and the point is only that one of the two has been cataloged as a target class and the other has not.

It is already known that glutamate-to-lysine (E→K) substitutions are common in cancer. They are common because glutamate is abundant and the dominant mutational process in cancer is C>T. Being numerous does not make these lysines worth targeting, which is a separate question.

A second question is why nobody has counted them. The answer is a measurement problem with two parts, both specific to this class rather than general to proteomics. Lysine is a tryptic cleavage site, so creating one redraws the peptide boundaries: the mutant peptide is not the wild-type peptide with a letter changed but a different species with a different mass, length and charge, which a reference database cannot contain and a conventional search cannot find. And the error control that would license the identifications it does make has mostly been tested in the wrong place, since target-decoy rates are calibrated over a whole search^9^ and variant peptides are a small, atypical subset of one. Strategies that control error separately for the variant space exist^7^, and entrapment methods that measure the realized rate within a subset now exist^10^. What is rarely done is to turn that measurement on a variant search and report what comes back.

This work turns it on one. The data are public; the measurement is not. Re-searching 42 published tandem mass tag (TMT) plexes of the Cancer Cell Line Encyclopedia (CCLE) proteomes^11^ against a mutation-aware database measures how much of the acquired-lysine class a search can see, and burden-matched entrapment measures the realized error on what it returns. Three published chemoproteomic screens answer the chemical question on the same residues, across four cell lines and 34 chemotypes. Every script and panel-level table is deposited, together with the site catalog and the site-level tables, so the census extends to a new search engine, a larger variant catalog or the next screen rather than having to be rebuilt.

## Results

### Cancer mutation creates a large, driver-enriched class of new lysines

Cancer makes a question available that a proteome-wide survey cannot: how many lysines does the disease itself create? Whether that is a class worth building chemistry around depends first on how large it is.

The class is large, and it is not the product of one dataset. Pooling missense sites from DepMap^12^, the TCGA MC3 call set^13^ and COSMIC^14^ gives 295,411 sites that create a lysine, across 20,365 genes (Figure 1A). The three sets are nested rather than independent, so the union is a catalog of what has been reported and not three replicate surveys.

**Figure 1.**
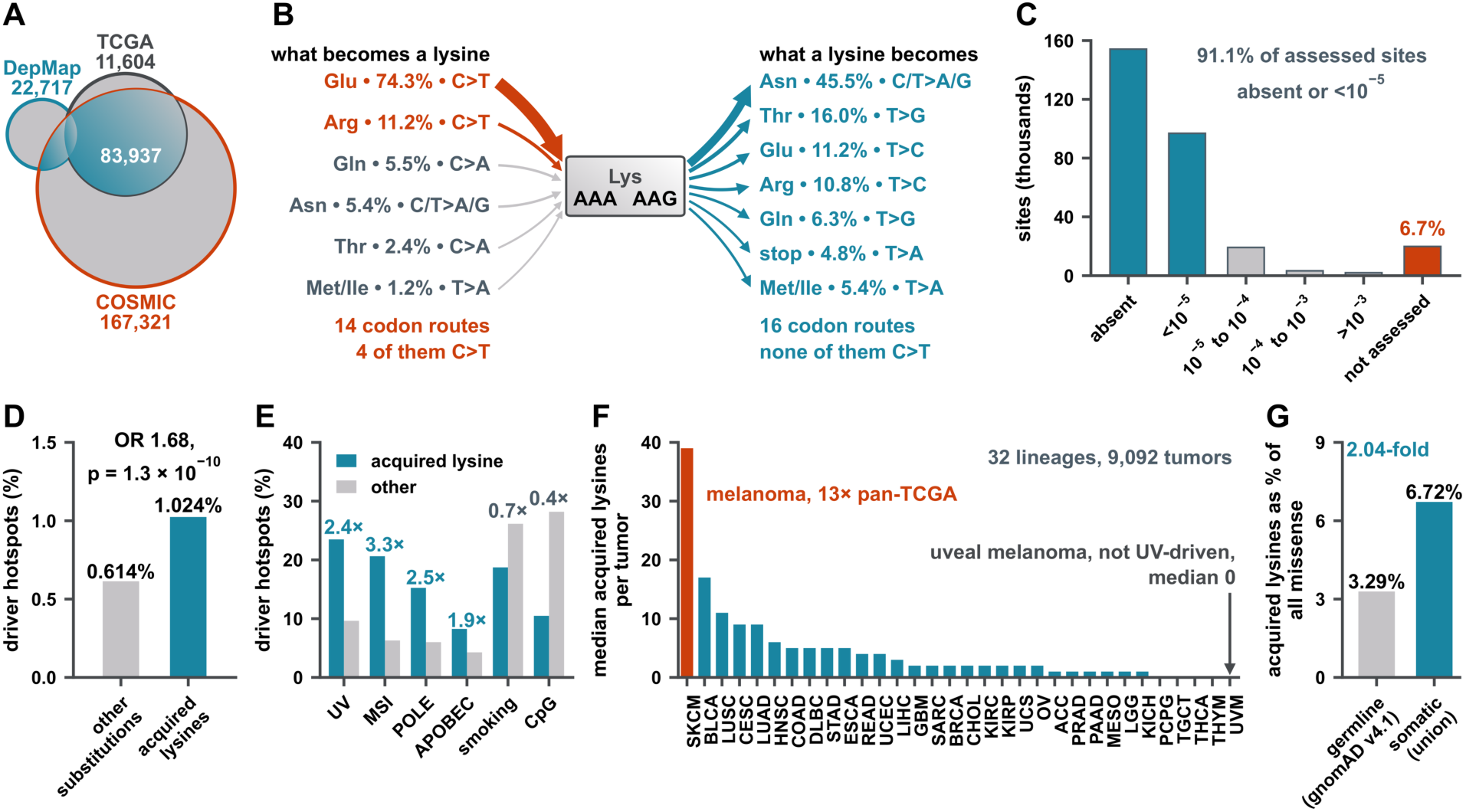
Cancer mutation creates a large, driver-enriched class of new lysines. Sites are missense positions in the union of DepMap, TCGA and COSMIC; a site is an acquired lysine if the substituted residue is lysine. The set and frequency panels are union-wide, the driver-hotspot panel is DepMap only because the flag is carried in that table, and the germline panel compares the union against two germline references. Counts are of sites, not of tumors or of individuals, except where the unit is stated. **(A)** The class, assembled from three variant sources. 295,411 acquired-lysine sites across 20,365 genes, from 4,395,403 missense sites; 93,769 (31.7%) supported by more than one source, which is the shaded territory. The three small overlap regions are unlabeled: 6,547 sites are shared by DepMap and COSMIC only, 147 by DepMap and TCGA only, and 3,138 by all three sources. **(B)** Codon routes into and out of lysine. Lysine is gained 85.5% by C>T and lost 0% by C>T. Rust marks the C>T routes, which are G:C to A:T changes, half of all cancer missense mutation. No C>T route leaves lysine because AAA carries no G or C, and the only G to A available to AAG gives AAA, which is still lysine. Arrow width increases with the share shown beside it. Values in Table S2. **(C)** gnomAD v4.1 allele-frequency bands over all 295,411 sites: absent 154,188; below 10^-5^ 96,892; 10^-5^ to 10^-4^ 19,194; 10^-4^ to 10^-3^ 3,297; above 10^-3^ 1,971; not assessed 19,869. 91.12% of assessed sites are absent or below 10^-5^, and the 6.73% never assessed is excluded from that denominator. **(D)** 1.024% of 19,239 acquired-lysine calls carry the Hess driver-hotspot flag against 0.6138% of 311,166 other missense calls, where a call is one variant in one cell line; odds ratio 1.68, Fisher exact p = 1.3 × 10^-10^, two-sided. **(E)** Share of driver hotspots by assigned signature, acquired lysines (n = 315) against other substitutions (n = 3,465): ultraviolet 23.5% against 9.6%, microsatellite instability 20.6% against 6.3%, POLE 15.2% against 6.0%, APOBEC 8.3% against 4.3%, smoking 18.7% against 26.1%, CpG deamination 10.5% against 28.2%. Here n counts sequencing entries, so a cell line sequenced more than once is counted each time. Counted once per cell line, the 315 become the 197 flagged calls of panel D. **(F)** Median acquired lysines per tumor across 32 TCGA lineages. Cutaneous melanoma 39, thirteen times the pan-TCGA median; uveal melanoma 0. All 32 are shown because a truncated ranking cannot carry that control. **(G)** Somatic union 6.72% (295,411 of 4,395,403) against germline 3.29%, 2.04-fold. The germline value is an expected diploid count per individual, 471.1 acquired lysines among 14,342 missense variants, and not a count of distinct variants. Values in Table S2. See also Figure S1.

Glutamate to lysine is the commonest substitution in cancer, and that is a statement about glutamate and about C>T mutation rather than about lysine. E→K ranks first of the 363 substitution pairs observed, but against a trinucleotide-context expectation its enrichment is only 1.12-fold, against more than three for each of the CpG hotspots R→H, R→C and R→Q (Figure 1B). Lysine is fourth of twenty residues by sites gained. What distinguishes it is the asymmetry of the routes: lysine is gained overwhelmingly by C>T and never lost by C>T, so a C>T-dominated process creates it and does not destroy it. The class is abundant for a mechanistic and unremarkable reason. The matrix in full, the gains and net flux residue by residue, and the signature control behind the asymmetry are in Figure S1.

The sites are somatic rather than common polymorphism. More than nine in ten assessed sites are absent from gnomAD v4.1^15^ or below an allele frequency of 10^-5^ (Figure 1C), and the sites never assessed are counted separately rather than folded into a denominator that would flatter the result. Frequency cannot exclude a private germline variant; only matched normals can, and these data have none.

Acquired lysines fall on driver hotspots more often than other substitutions do, at an odds ratio of 1.68 (Figure 1D). The flag is the one carried in the DepMap variant table rather than a gene list chosen here, and it is conservative by construction, since the method exists to strip recurrent passenger sites from hotspot lists^16^. Hotspot recurrence is a statement about where mutation is selected, not about whether the resulting protein is expressed or reachable.

The excess tracks mutational processes that make lysine, not mutation in general. Split by assigned signature, the four processes that generate the relevant base changes are all enriched among acquired-lysine driver hotspots, and the two that do not make lysine run the other way; both directions are labeled (Figure 1E).

Burden is lineage-specific, and the ranking carries its own control: cutaneous melanoma leads all 32 lineages, and uveal melanoma carries zero (Figure 1F). Both are melanocytic and only the first is ultraviolet-driven, so a property of the lineage rather than of the mutational process would make both high.

The excess survives a germline reference chosen to remove the obvious objection. Acquired lysines are twice as large a share of missense in the somatic union as in the germline (Figure 1G). Because gnomAD contains cancer-affected individuals, the same pass was run over its non-cancer exome subset, and the two agree to within 0.12 percentage points. They are not independent cohorts, both being gnomAD with heavy overlap, so what the second controls is specifically the presence of cancer-affected individuals and not reference bias in general. Repeating the pass for every residue on the identical file, filter and frequency field ranks lysine third of twenty and cysteine fourth, so the comparison the field draws with acquired cysteine can be made on one calculation instead of two, and on that calculation lysine is not the weaker case. The ratio is not a measure of selection: germline shares are depressed by purifying selection, which is the likeliest reading of phenylalanine and tyrosine, the only two residues ranking above lysine.

Together these exclude the cheapest alternative to the premise of this paper, that acquired lysines are simply what mutation does everywhere. They are created at twice the germline rate, they are enriched on driver hotspots, and the processes that make them are identifiable. None of that says the resulting proteins are expressed, detectable or reachable by a molecule, and the rest of the paper is about how far each of those holds.

### The class is mostly missed by a search built to find it

Whether anyone has been looking at these lysines is answerable on the 42 TMT plexes of the CCLE proteomes^11^, which cover 375 cell lines and are among the deepest shotgun datasets available. The unit is the site-in-cell-line observation rather than the distinct genomic site, because each line is an independent chance to detect. Four peptide species can carry an acquired-lysine site and only two of them contain it (Figure 2A).

**Figure 2.**
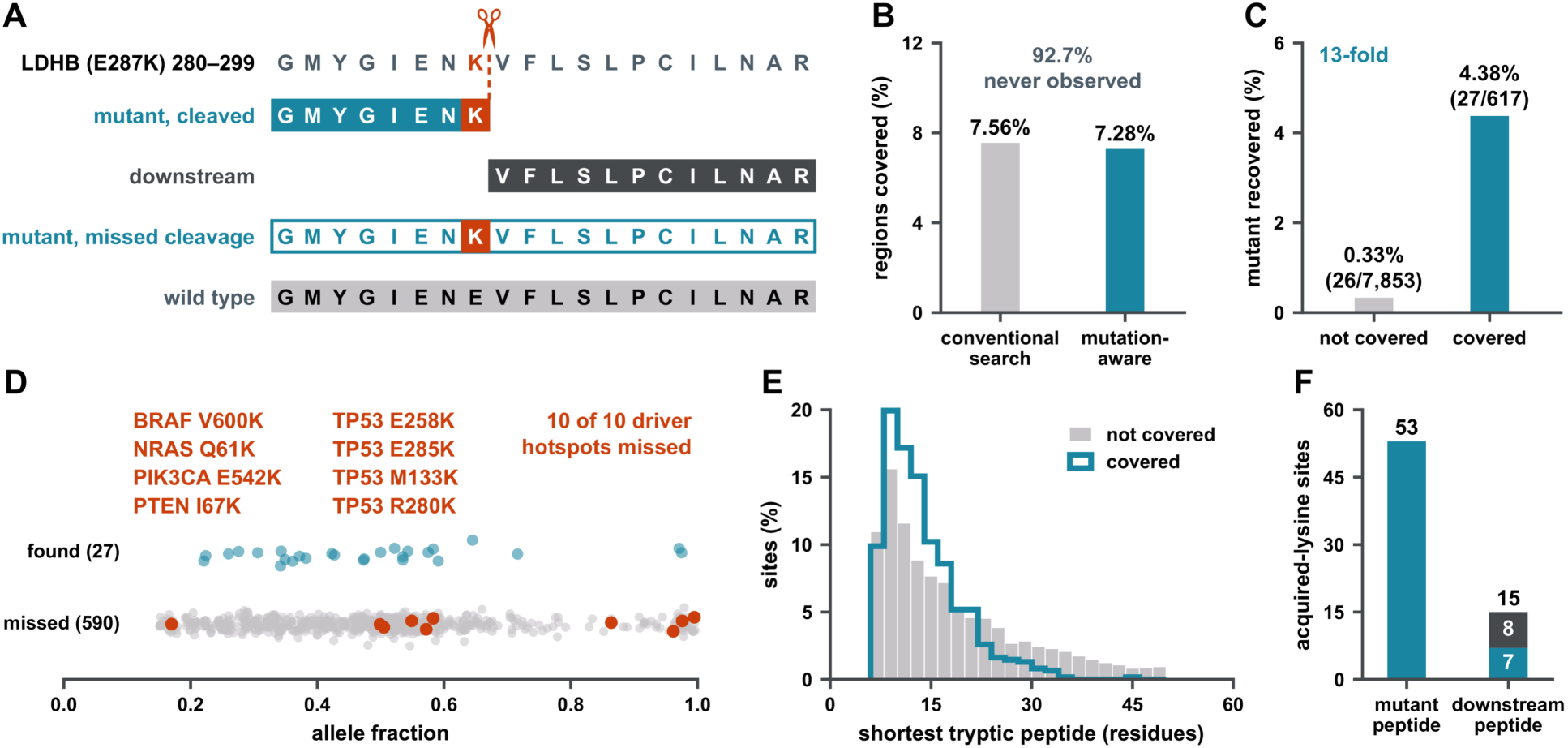
The class is mostly missed by a search built to find it. 42 TMT plexes across 375 cell lines. The unit is a site-in-cell-line observation, not a distinct genomic site: 8,470 acquired-lysine observations covering 8,055 distinct sites across 372 lines, with 1,902 non-lysine observations carried through the identical pipeline as the control (Figure S2). Every rate is per observation. Detection is counted at two gates that fail for different reasons and are never combined: gate 1 is whether the wild-type region was observed at all, gate 2 whether the mutant peptide was recovered where it was. **(A)** Four peptide species can carry an acquired-lysine site, and only two contain it. One real site, written out in full: LDHB E287K in NCIH1793, residues 280–299 of UniProt P07195, GMYGIENKVFLSLPCILNAR, with the acquired lysine at position 287. The scissors and dashed line mark the cleavage site that lysine creates; the residues flanking the peptide in the protein, lysine 279 and glycine 300, are not drawn. The acquired lysine, marked in rust on the two species that carry it, subdivides that peptide into an 8-residue mutant peptide and a 12-residue downstream peptide; the missed-cleavage form spans all 20, and the bars are drawn on one residue pitch so their lengths are those counts. Sequences and lengths are computed by digesting P07195 with and without the substitution, not transcribed. The mutant peptide and its missed-cleavage form carry the mutated residue; the downstream peptide and the wild-type peptide do not. An acquired lysine creates a tryptic cleavage site, so the mutant peptide has different boundaries from the wild-type peptide; the two are not one sequence with a letter changed. That same new boundary is what releases the downstream peptide, which therefore carries no mutated residue and still exists only in the mutant protein, and is why it can report a site it does not contain. Values in Table S2. **(B)** Both bars are the same 8,470 observations. Values in Table S2. **(C)** The mutant peptide was detected at 27 of the 617 covered sites (4.38%) and at 26 of the 7,853 uncovered ones (0.33%) by the mutant-only route, 53 sites in all, 0.63% of the class, on 52 distinct genomic sites, at the search’s reported 1% false-discovery rate, which is where the panel is drawn. Detection is by a cleaved species, a missed-cleavage species, or both. Values in Table S2. **(D)** The 617 covered sites arrayed by allele fraction and split into the 590 whose mutant peptide was missed and the 27 where it was found. All ten driver hotspots in that set fall on the missed row, eight gene and mutation labels across five genes, listed in Extended Results. NRAS Q61K was recovered in one line by the mutant-only route, where the wild-type region was never observed, so that observation is one of the 26 in Figure S3A and not in this population. **(E)** Where the region is never covered, the wild-type tryptic peptide is longer, and only in the tail. The shortest tryptic peptide covering the site, as a percentage of the sites in each group, so the two groups are comparable despite differing 12-fold in size. Mean 17.5 residues (n = 7,467) where the region was never observed against 13.3 (n = 617) where it was covered, Mann– Whitney p = 2.2 × 10^-15^. The difference is entirely above 20 residues. At or below 20 the two groups are one distribution, mean 11.70 residues against 11.71, p = 0.38; above 20 they separate and keep separating, 30.8% of never-covered sites against 12.0% of covered, and 13.3% against 1.1% above 30. Because each group is drawn as a percentage of itself, the taller covered peak at 8 to 10 residues is the absence of a tail and not a shorter body. 386 never-covered sites are excluded, every one of them a region for which no wild-type tryptic peptide was enumerated at all, where the pipeline writes a length of 0; no covered site is affected. A gate-1 result, kept separate from gate 2 because the two failures have different causes. **(F)** The downstream peptide gives 15 sites, 7 shared with the first route and 8 new, 61 in the union. It returned zero entrapment hits across 42 experiments (Figure S4D) against 0.58% for variant-containing peptides in the same searches, which is why the two are kept apart. Values in Table S2. This figure states what has been looked for and found, in these cell lines, with this workflow. It is not evidence that the peptides are absent from the sample. See also Figures S2 and S3.

Gate 1: the region is almost never covered. A conventional search and a mutation-aware search over 10.17 million peptide entries observe the wild-type region at the same rate, agreeing to within 0.3 percentage points, so coverage is a property of the sample rather than of the database (Figure 2B). More than nine in ten of these sites are never covered in any of the 42 plexes, and 26 of them are detected only through the mutant peptide itself (Figure 2C).

Gate 2: where the region is covered, the mutant peptide is still usually missed. Fifty-three detections across the two routes are 0.63% of the class, and 30 of them also clear the score margin at which the next section measures the realized error rate, 0.35% (Figure 2C). The two thresholds are not interchangeable. The larger is the census of what the searches returned, the smaller the subset over which the false-discovery proportion is 0.83% rather than 4.70% (next section). The acquired-lysine class contributes zero entrapment peptides against 38 variant peptides at that margin, so its own proportion is bounded at 7.89% by the rule of three rather than measured at zero. Under the conventional search the mutant count is zero, which is a passed control rather than a failure, since that database contains no variant sequences and can speak to coverage and to nothing else.

The misses include the driver hotspots. Every driver hotspot among the covered sites falls on the missed side, across five genes (Figure 2D). Driver status is the DepMap hotspot flag rather than a gene list chosen here^16^, and allele fraction is on the axis so that the misses can be seen not to be confined to subclonal sites. The panel counts hotspots among the covered sites, which is the population it plots: NRAS Q61K was recovered in one line by the mutant-only route, where the wild-type region was never observed, so that observation is not in this population.

Where the region is never covered, the shortest tryptic peptide spanning the site is the longer one, and the two distributions diverge in the tail rather than shifting as a whole (Figure 2E). The mutant peptide has the opposite geometry, and the search absorbs it rather than being stopped by it. A new lysine is a new cleavage site, so the fully cleaved mutant peptide runs about half the length of the wild-type peptide over the same site, and half the class falls below the seven-residue minimum that this search and the original one both enforced (Figure S3F). That is not a second hidden exclusion. Both searches allowed two missed cleavages, and an upstream missed cleavage returns a too-short peptide to the window, so 95% of observations still carry a searchable species ending at the acquired lysine. What the geometry costs is identification quality rather than existence, and whether it costs detections cannot be settled here.

The non-lysine control demonstrates the mechanism rather than controlling it away: variant sites that do not create a lysine are covered less and recovered more, the direction expected if peptide boundaries are what matters, and the median lengths differ by one residue so it is not a length artifact (Figure S2).

A second, independent route reaches the same class of site. The tryptic peptide released downstream of the new cleavage site carries no mutated residue but exists only if the cleavage happened (Figure 2F). The two routes are reported separately rather than pooled because their error properties differ, which is the subject of Figure 3.

**Figure 3.**
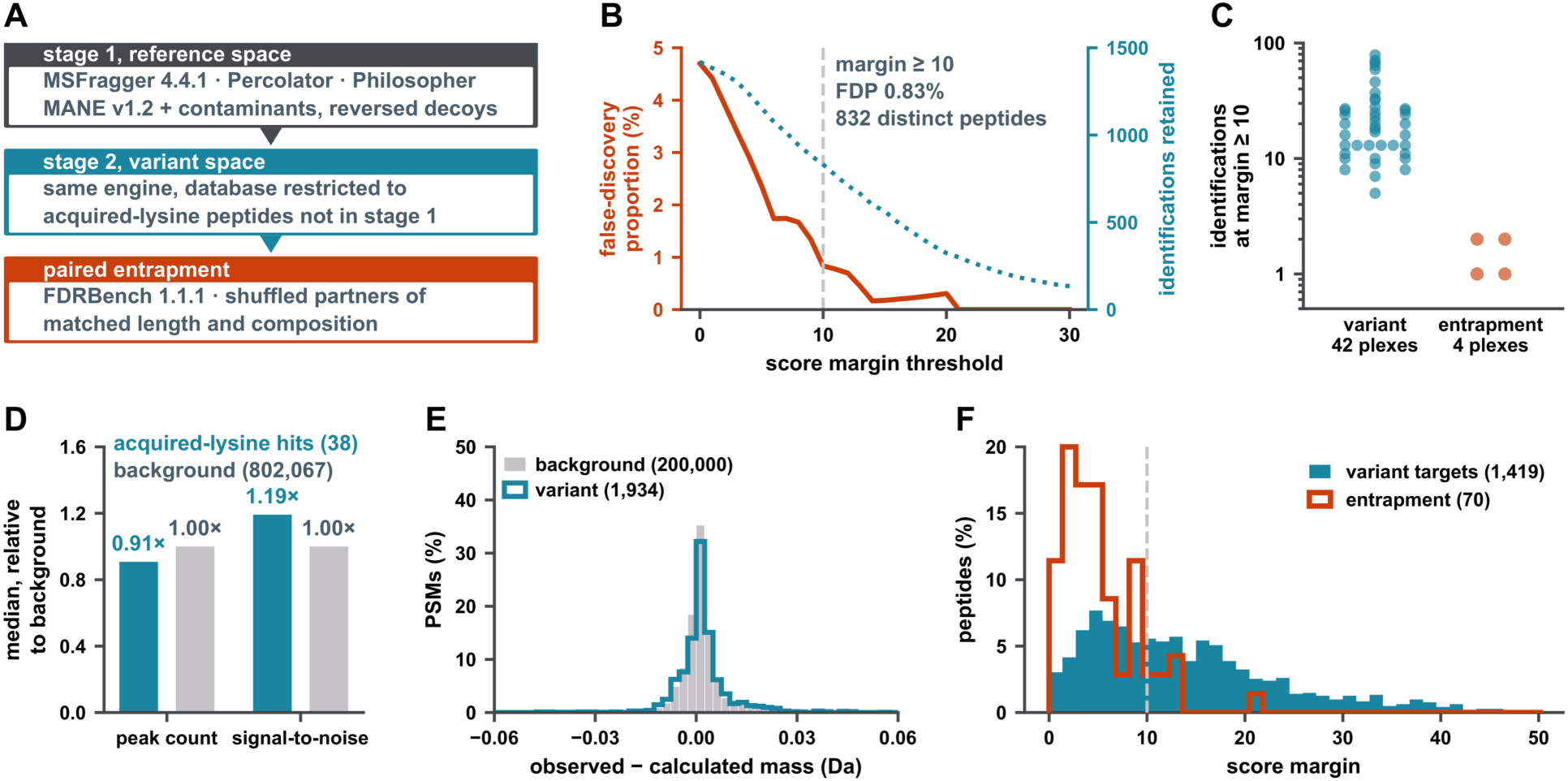
Reported error rates do not hold for variant peptides. Burden-matched entrapment at r = 1 (Methods). Every entrapment identification is by construction false. Two denominators appear and are not interchangeable: panel B counts distinct peptides, one best spectrum each; Extended Results counts identifications. **(A)** The design, matched to the variant set on the properties that drive identification, so the comparison is like for like (Methods): 589,588 variant-pair rows across eight plexes, entrapment sets of 147,397 each, all of them sharing residues with their target (Figure S4A to S4C). **(B)** Margin dose-response over 1,489 rows, counted as distinct peptides. Past a margin of 15 only one entrapment peptide remains, so the rate there reflects a single count rather than a cleaner set. A 1% reported false-discovery rate is not a 1% realized one for this class of peptide unless a margin is applied. Restricted to the acquired-lysine class the proportion is bounded at 7.9% by the rule of three, with the counts and the comparison against the rest of the variant class in Extended Results. Values in Table S2. **(C)** Per-plex consistency. Identifications at margin ≥ 10, one point per plex on a log axis, with the two populations side by side. Six among 1,039 is 0.58% pooled, and it is spread across four experiments rather than carried by one. This is the identification denominator, not panel B’s peptide one. Values in Table S2. **(D)** Median spectral-quality features for the 38 acquired-lysine identifications expressed relative to 802,067 background spectra from 20 of the 77 runs in the same searches, so 1.0 is the background. These features score a spectrum on its own properties, with no reference to the search engine, the database or the false-discovery rate. The plus one dalton tail accounts for 7.7% of variant and 7.3% of background spectra. SQSGM is omitted because the MGF export zeroed its intensity term, which would collapse a geometric mean. Values in Table S2. **(E)** Precursor mass error. Observed minus calculated precursor mass, variant against background, drawn as the percentage of each group falling inside the plotted window, in bins of 0.0025 Da. 91% of all 1,934 variant peptide-spectrum matches (PSMs) are within 0.05 Da, and inside the window the medians are 0.0013 Da for variant against 0.0009 Da for background. A tail sits at +1 Da and takes 7.7% of variant PSMs against 7.3% of background, the first isotope selected instead of the monoisotopic peak at the same rate in both, which makes it an instrument behavior rather than a property of variant peptides. **(F)** Score-margin distributions for variant and entrapment identifications, each drawn as a percentage of its own total because 1,419 cannot be compared with 70 on counts. Entrapment peptides are identified, and they score like a population that cannot be present should, which is what makes the margin in panel B do work rather than simply discard the weakest identifications of every kind. 7 of the 70 entrapment peptides clear the margin of 10 and 90% sit below it. The axis ends at 50; 14 of the 1,419 variant peptides score above it, up to 69.4, and are not drawn. Pooled false-discovery proportion across all plexes is 0.58% as identifications, six among 1,039 and appearing in four of 42 plexes (Extended Results). Panel B’s 0.83% is seven among 839 distinct peptides, in five plexes. The difference is one peptide at a margin of 10.53, which clears the threshold in the recomputed margins behind panel B and not in the per-plex summary behind panel C. See also Figure S4.

What the evidence behind these counts will bear is limited, and Figure 3 exists because of it: four in five of the mutant-peptide sites rest on a single spectrum, and none on more than one experiment (Figure S3E). Three alternative explanations for the gate-1 failure were tested and none accounts for it, and what predicts identification is incomplete digestion rather than charge (Figure S3A to S3D). Gate 1 is a statement about tryptic geometry and gate 2 about search sensitivity, and neither licenses a claim about what the proteome contains.

### Reported error rates do not hold for variant peptides

Figure 2 recovered a small number of mutant peptides, and a result carried by single-spectrum identifications has to survive an error-rate audit before anything is built on it. A global 1% filter does not settle it, for the reason given above: a rate that holds over a whole search need not hold within a tiny, atypical subset of one^9^.

Chemoproteomic work on mutation-created residues has answered this in two ways, and both control the error: one searches canonical and cell-line-specific mutant sequences in a single pass and filters globally^17^, the other in two stages, removing spectra confidently assigned to canonical sequences before the variant database is searched^7^. Neither reports a realized false-discovery proportion measured within the variant class, which is what follows, and what follows is a property of this search and this database, not of theirs.

The test is burden-matched entrapment, an application of a published method rather than a new one^10^. Each variant peptide is given a partner built from its own residues, permuted, with the C-terminal lysine or arginine kept in place (Figure 3A). The pair therefore shares an exact precursor mass and an identical length, and only fragment ions can separate them, which is what makes this an upper-bound test rather than a decoy in a different costume. An entrapment sequence cannot be in the sample, so every entrapment identification is a false one, counted directly rather than estimated.

At the reported filter the realized error rate is roughly fivefold higher than the nominal one, and a score margin recovers the intended rate (Figure 3B). The margin is the gap between a spectrum’s best interpretation and its next best, and requiring it to be large removes exactly the identifications that a mass-matched impostor could have produced. Every error rate quoted in this paper, and every identification quoted from the entrapment analysis, is taken at a margin of 10 or more; Figure 2’s detection census is reported at the search’s own 1% filter and is given at both thresholds where it appears.

Three properties of the identifications themselves agree with that reading, and none of them refers to the search engine or the database. The entrapment rate is spread across plexes rather than carried by one experiment (Figure 3C), the spectra score close to the background on peak count and signal-to-noise (Figure 3D), and the precursor mass errors sit in the same window as the background, including the first-isotope tail, which appears at the same rate in both (Figure 3E).

The margin does work rather than simply discarding the weakest identifications of every kind. Entrapment peptides are identified, and they score like a population that cannot be present should (Figure 3F), and it is that separation the margin exploits. Neither rate is a per-site confidence, and a single spectrum past the margin remains a single spectrum, which is why Figure 2’s detections are treated as a lower bound on coverage rather than a catalog of sites.

### Reactivity and ligandability separate only where the structure is trustworthy

Two questions are routinely run together in work on covalent lysine chemistry: what makes a lysine reactive enough to be labeled by a probe at all, and what makes a labeled lysine blockable by a competing molecule^1^. One contrast cannot answer both, so two were run against two different control groups (Figure 4A). Reactivity compares probe-labeled lysines against every other lysine on the same protein; ligandability compares engaged lysines against all labeled ones. Both run through the same ladder of structural-confidence gates, always applied to both groups at once, since gating one side would manufacture a difference rather than reveal one.

**Figure 4.**
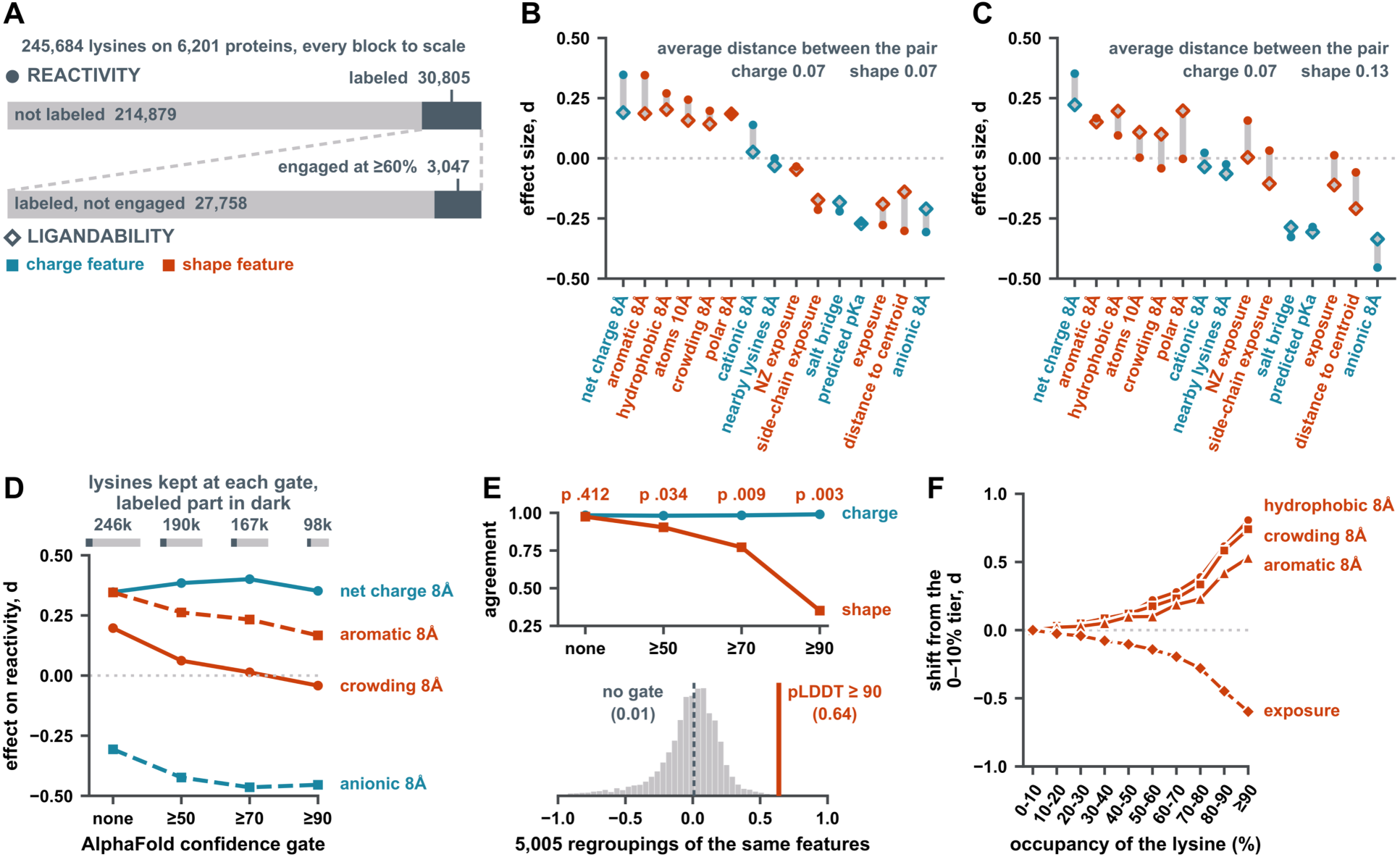
Reactivity and ligandability separate only where the structure is trustworthy. Reactivity and ligandability are measured on the same lysines and with the same estimator, one difference per protein tested against zero, so the two contrasts are directly comparable. The figure compares site sets; individual residue numbers within them are provisional (Methods). Effects are within-protein standardized differences, measured in cell lines. **(A)** Two comparisons against two control groups, run before and after a structural-confidence gate so the gate’s effect is visible rather than assumed. 245,684 lysines across 6,201 proteins, 30,805 of them labeled (12.5%). Fifteen features, six charge and nine shape, the identical set on both ladders^18,19^, with the AlphaFold terms excluded (Methods). **(B, C)** The paired display, ungated and at a confidence of 90 or above. Each feature’s two effects are drawn on one line, a filled circle for the effect on being labeled and an open diamond for the effect on being blocked, so the movement under the gate reads directly. Values in Table S2. **(D)** The gate, step by step. Four representative features’ reactivity effect at each rung. Charge features hold or strengthen across the ladder; shape features fall. The gate therefore costs 60% of all lysines by its tightest rung while raising the labeled share from 13% to 18%, so the two groups do not shrink at the same rate. Values in Table S2. **(E)** The permutation test, over the gate ladder and against the null. Two stacked axes. Agreement is the correlation, across the features of one kind, between a feature’s effect on reactivity and its effect on ligandability. Printed above each gate is the exact one-sided permutation p for the gap between the two kinds at that gate: 0.412, 0.034, 0.009 and 0.003. Both are stated because either alone misleads, and Figure S5G is the control against the gate keeping only the strongest sites. Values in Table S2. **(F)** Occupancy dose-response. Shift from the 0–10% occupancy tier as occupancy rises, so the result does not depend on the 60% engagement cut existing at all. Monotonic across all ten tiers, with no discontinuity at the 60% engagement cut. The fully threshold-free version is Figure S7C. See also Figure S5.

Ungated, the two contrasts are indistinguishable. The mean distance between a feature’s two effects is the same for charge and for shape features, and within each class the two contrasts track each other almost perfectly (Figure 4B). An exact permutation over all 5,005 ways of splitting fifteen features six against nine gives p = 0.412. Read on its own, this panel says that reactivity and ligandability are one property measured twice.

At a confidence of 90 or above the two contrasts come apart, and only across the shape features (Figure 4C). The mean pair distance for shape nearly doubles while charge is essentially unchanged, the shape class correlation falls from 0.97 to 0.35 while charge holds at 0.98 to 0.99, and the permutation value rises to p = 0.003.

The packing features are where it happens, and the spread shows it is not range restriction: in each case the feature’s own standard deviation holds or widens under the gate, so the reactivity effect is not shrinking because the spread shrank (Extended Results). Two shape features are exceptions and are reported as such. Exposure and distance to the centroid lose 12% and 29% of their spread under the gate, so for those two the collapse is partly range restriction rather than artifact removal.

Gating costs sites. 30,805 labeled lysines fall to roughly 17,000 at a confidence of 90 or above (Figure 4D), and some of the discarded sites are engaged, so this is a real cost rather than a free cleanup.

What Figure 4E measures is whether the two contrasts move together, not whether either is stable. Several charge effects change substantially under the gate, but they change in step on both sides. The shape features do not: their reactivity arms collapse toward zero or reverse sign while their ligandability arms hold direction. That asymmetry, not the size of any single effect, is the result.

Both arms of the test have to be reported together, because either alone misleads. Across the ladder the shape coupling falls monotonically while charge holds throughout (Figure 4E). The separation therefore does not exist before the gate is applied, and it is not manufactured by the gate keeping only the strongest sites, which is the control in Figure S5G. Nor does it depend on the 60% cut existing, since the shift from the lowest occupancy tier rises monotonically across all ten tiers with no discontinuity where the cut falls (Figure 4F).

The claim is conditional. Where predicted structure is trustworthy, shape distinguishes ligandability from reactivity and charge does not; ungated, neither does, because the apparent shape contribution to reactivity is carried by sites whose geometry cannot be measured. This does not say that shape is irrelevant to reactivity in general; it says that the evidence for it in these data comes from the sites where the evidence is weakest. It also inherits the gate’s own bias: AlphaFold confidence is not random with respect to biology, and a claim that holds only above a confidence of 90 is a claim about ordered protein.

### Ligandability reproduces at the site level and resists prediction from site structure or compound properties

If ligandability is a property of the site rather than of the molecule, the same lysines should be preferred by compounds that have little else in common. Figure 4 measured which site properties separate engaged lysines from merely labeled ones; what follows is how well they predict ligandability, and whether any property of the compound predicts which lysines it prefers. Across the three screens^2–4^, 0.49% of site × compound measurements reach the 60% occupancy that defines engagement here (Figure 5A), and the distribution is smooth through 60% (Table S2), so the threshold is a line drawn through a continuum rather than a boundary in the data. For each compound separately, the sites it engaged are compared with the sites it was quantified at and left below the cut, and each site feature is standardized within that compound, so what is compared is compounds’ preferences rather than their site lists, and a compound tested on a different set of lysines cannot manufacture a difference.

**Figure 5.**
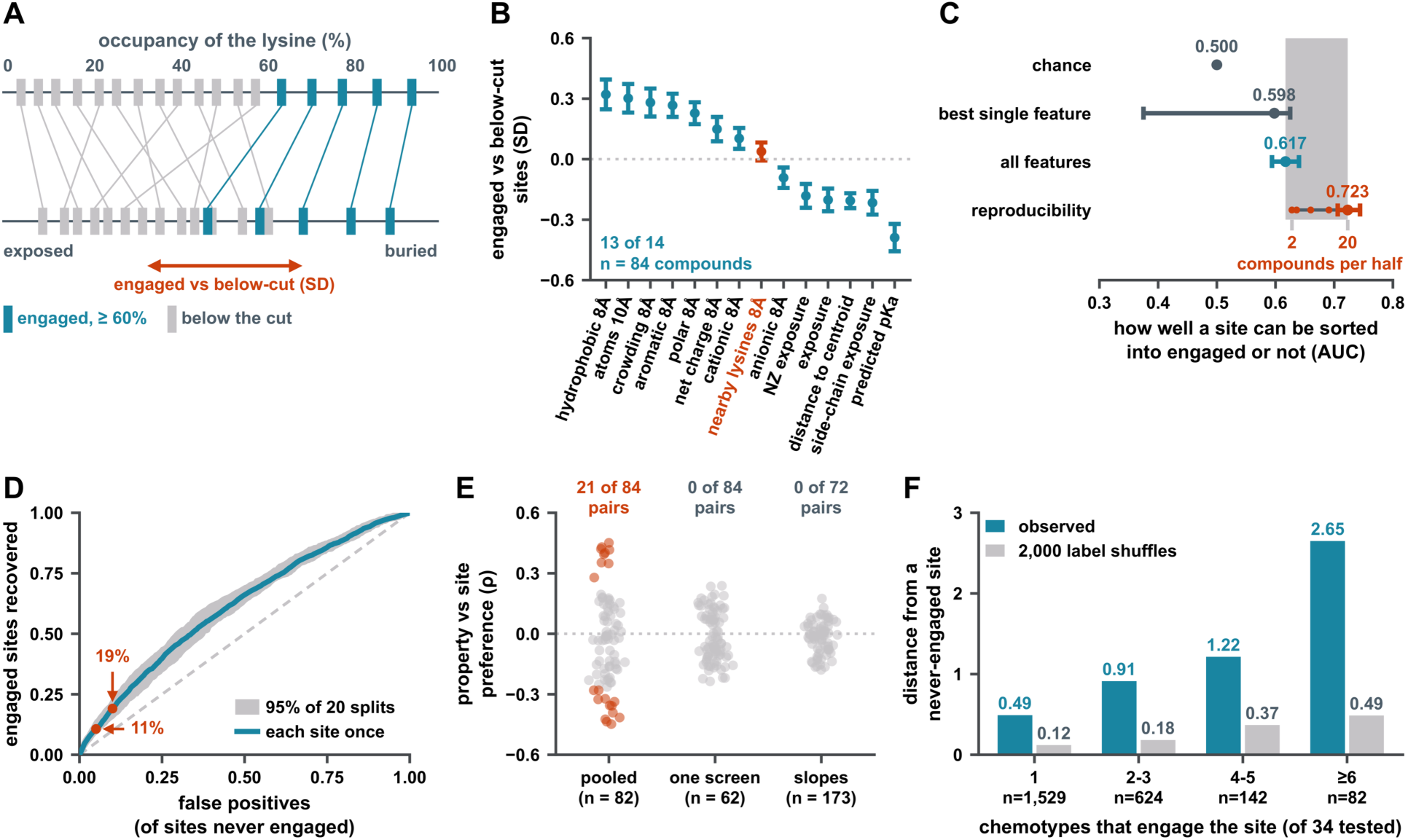
Ligandability reproduces at the site level and resists prediction from site structure or compound properties. Ligandability measured in three chemoproteomic screens (the lysine-targeting compound screen in Ramos and MDA-MB-231^2^; the meroterpenoid screen in MDA-MB-231 and MDA-MB-436^3^; the *N*-(cyanomethyl)-*N*-(phenylsulfonyl)amide (CyMPSA) screen in HeLa^4^), with engagement defined as ≥ 60% occupancy, 100 × (1 − compound / DMSO). The composite is a logistic model on fourteen site features, cross-validated by protein (Methods); the site features of panel B are a different set, and the two are never pooled. Intervals on model AUCs are the 2.5th to 97.5th percentile across 20 held-out splits, and on ceilings across half-splits (Methods). **(A)** What is measured, and what is done with it. A probe labels lysines; a compound competes with the probe, and occupancy is the share of probe labeling it competes away. The median compound quantifies 6,332 lysines and engages 14. The schematic is the design for one compound, its engaged sites against the sites it was quantified at and left below the cut. Values in Table S2. **(B)** Thirteen of fourteen site features point the same way in every compound. Mean standardized shift between engaged and below-cut sites with 95% intervals, across n = 84 compounds. Rust marks the single exception, nearby lysines within 8 Å (+0.038, q = 0.097). See also Figure S7B. **(C)** The measured model against the reproducibility ceiling. The ceiling drawn is the lysine-targeting compound screen’s at twenty compounds per half, the deepest split the data support. That gap is reproducible signal the structure does not reach. It is not overfitting, since optimism is +0.002 AUC. The ceiling is still rising at the deepest split, so 0.723 is a lower bound. See also Figures S6B and S6D. Values in Table S2. **(D)** What the model buys at any operating point. Receiver-operating curve for the composite, every site predicted once, with the band spanning 95% of 20 held-out splits. There is no operating point where this model is useful. Values in Table S2. **(E)** No compound property predicts which lysines a compound prefers. Every descriptor × site-feature-preference correlation, one point per pair, with rust marking pairs clearing q < 0.05. This is a statement about the descriptors that were tested, which Figure S7F and note N7 list. Values in Table S2. **(F)** Breadth across chemotypes. Sites grouped by how many of the 34 tested chemotypes engage them at ≥ 60% occupancy, against multivariate distance from a never-engaged site: one compound, n = 1,529, 0.491; two or three, n = 624, 0.913; four or five, n = 142, 1.215; six or more, n = 82, 2.650. Gray is the null over 2,000 label shuffles (0.120, 0.183, 0.369, 0.487); all four bins clear it at p < 0.0005, the resolution limit of 2,000 shuffles. What marks a site is not which chemistry engages it but how many can. See also Figure S6E. Engagement measured in cell lines. The figure compares site sets; individual residue numbers within them are provisional (Methods). See also Figures S6 and S7.

Thirteen of fourteen site features point the same way in every compound: engaged lysines are consistently more buried, more crowded, more hydrophobically packed and lower in predicted pK_a_ (Figure 5B). The single exception is the count of nearby lysines within 8 Å, which is a property of the neighborhood rather than of the lysine in question. The three charge features clear significance but are the three smallest of the thirteen that do, and two of them do not strengthen with occupancy (Figure S7B).

A logistic model was fitted on fourteen features, thirteen describing the lysine’s environment and one describing how confidently the residue was modeled, and cross-validated by protein rather than by site, since a site-level split leaks information between training and test (Methods). Held out, the composite reaches an area under the receiver operating characteristic curve (AUC) of 0.617, and its optimism against the training fit is +0.002, so the number is not depressed by overfitting (Figure 5C). Refitting on the thirteen environment features alone gives 0.611, which forecloses the objection that the model is partly predicting how well a site was modeled rather than what the site is.

Whether 0.617 is disappointing depends on what a perfect model could reach, and that limit is set by the assay. Engagement is the outcome of a measurement: if the same lysine is assayed twice and the two assays disagree, no predictor can agree with both. Splitting each site’s compound measurements into two independent halves, taking the label from one half and the score from the other, gives the AUC a perfect structural model would reach if structure captured everything the assay captures. At the deepest split the data support, that ceiling is 0.723 and the composite falls 0.106 short of it (Figure 5C). The shortfall is the smallest the data permit, on two counts. Each half carries fewer compounds than the full screen and is therefore noisier than the real label, so the ceiling is a lower bound and is still rising at the deepest split; and 0.723 is the lowest of the three screens, the other two sitting between 0.90 and 0.95 in every band where they are populated. The comparison is drawn against the weakest ceiling in the data, and is quoted as ≥ 0.72.

No operating point on the curve would change a decision. At a 5% false-positive rate the composite recovers 11% of engaged sites, and at 10% it recovers 19% (Figure 5D). Nothing on this curve would alter which lysines a screen went after, which is the practical content of the result: ligandability at these sites, in these cell lines, has to be measured.

No tested compound descriptor predicts which lysines a compound prefers. Correlating every compound descriptor against every site-feature preference gives 21 of 84 pairs clearing q < 0.05 when the three screens are pooled, against 4.2 expected, but 0 of 84 within one screen alone and 0 of 72 in the larger within-dataset slope analysis (Figure 5E). The pooled positives are provenance rather than chemistry: molecular weight all but names the screen a compound came from, each screen is a different cell line in a different sample state, and the descriptors that separate the three screens are the descriptors that produced the pooled positives (Extended Results).

What separates ligandable sites is breadth of engagement rather than a structural signature. Grouping sites by how many of the 34 tested chemotypes engage them at ≥ 60% occupancy, the distance from a never-engaged site rises monotonically across the four groups, from 0.49 to 2.65, against nulls of 0.12 to 0.49 from label shuffles (Figure 5F). The direction is secure across the gradient; the magnitude at the far end rests on 82 sites, and adjusting for how often each site was tested does not remove it (Figure S6E).

Both halves of the heading are bounded. Reproducible is a measurement rather than an assumption, and the same split-half analysis supplies the ceiling. Poorly predicted is bounded to these features, these compounds and these cell lines. The descriptors tested contain nothing that encodes charge, the largest correlation any of them reaches with a charge-related preference being 0.193 against a median of 0.552 for molecular weight, and every shape term among them is achiral by construction, since reflecting a conformer leaves the principal-moment ratios unchanged^20^ (Figure S7F). Ligandability is a property of the lysine over the descriptors tested, and the analysis is silent on exactly the two axes this paper goes on to test, charge in Figure 6 and chirality in the enantiomer pairs of Extended Results, both of which show molecular properties that do matter. The claim is about which site a compound prefers, not about whether a molecule’s properties matter at all.

**Figure 6.**
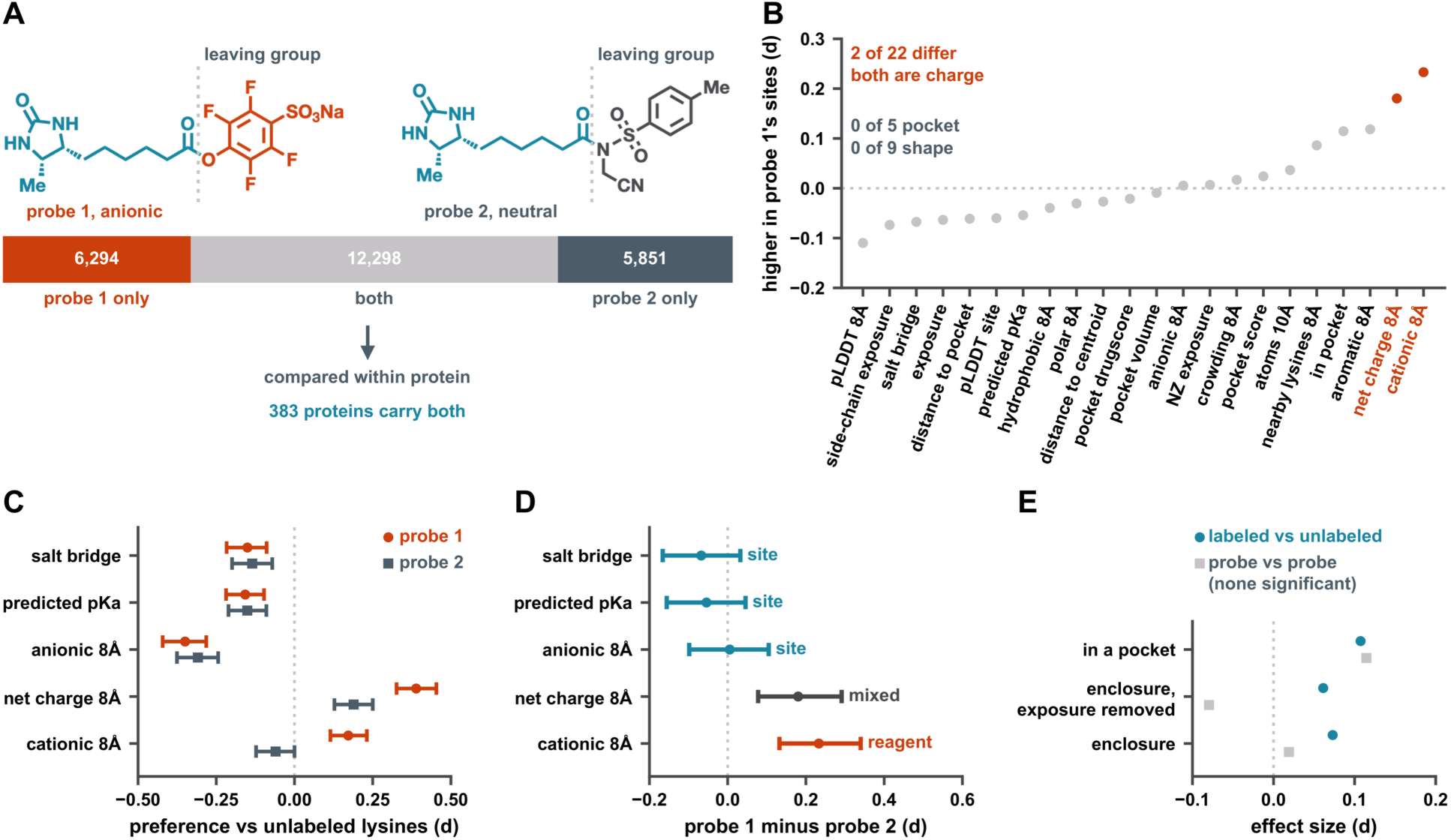
The probe is not a neutral reporter: reagent charge steers labeling. Two probes sharing a desthiobiotin-hexanoyl core and differing only in the acyl leaving group, applied to the same HeLa proteome. Twenty-two site features measured on 167,057 lysines at an AlphaFold per-residue confidence (pLDDT) ≥ 70^21,22^, listed with their sources in the Methods. Labeling is expressed as % occupancy. **(A)** The design. dTB-STP, a 4-sulfo-2,3,5,6-tetrafluorophenyl ester and anionic, against a neutral N-(cyanomethyl)-N-(4-methylbenzenesulfonyl)amide probe. 6,294 sites labeled only by the anionic probe, 5,851 only by the neutral one, 12,298 by both; the discriminating contrast is drawn within the 383 proteins carrying both. **(B)** Two features of twenty-two differ, and both are charge. Cationic residues within 8 Å d = +0.233 (q = 1.6 × 10^-4^); net charge within 8 Å d = +0.180 (q = 5.2 × 10^-3^). Zero of five pocket features and zero of nine shape features differ. Figure S8A shows each probe is individually informative, eight of twenty-two features significant for each against unlabeled sites, so this is a statement about the contrast rather than about power. **(C, D)** What each probe prefers, and which part of the charge effect belongs to which. Effects are standardized differences across proteins; intervals are 95% percentile bootstrap confidence intervals, resampling proteins (Methods). Zero of five pocket features and zero of nine shape features differ. The q values and the full twenty-two-feature test are in Figure 6B. Values in Table S2. **(E)** Concavity belongs to the site, not the reagent. See also Figures S8B and S8C. Values in Table S2. See also Figure S8.

### The probe is not a neutral reporter: reagent charge steers labeling

Charge was never among the descriptors tested for ligandability (Figure S7F), so the question is open. It is put to the reagent rather than to the compound, because in a competitive experiment the probe decides which lysines are visible to be competed for at all.

Two probes were compared that share a desthiobiotin-hexanoyl core and differ only in the acyl leaving group (Figure 6A). One carries a 4-sulfo-2,3,5,6-tetrafluorophenyl ester, anionic at physiological pH^1^; the other an N-(cyanomethyl)-N-(4-methylbenzenesulfonyl)amide, neutral overall^4^. Both were applied to the same HeLa proteome, and the discriminating contrast is drawn within the proteins carrying sites of both kinds, so a probe that happened to sample a different set of proteins cannot create a difference.

Two site features of twenty-two differ between the probes, and both are charge: lysines labeled only by the anionic probe sit among more cationic residues, and in more positive local environments overall. No pocket feature and no shape feature differs (Figure 6B). That pattern is what a steering effect should look like and what a general difference in proteome sampling should not. Neither probe is individually quiet: measured against unlabeled lysines, eight of the twenty-two features are significant for each (Figure S8A), so two of twenty-two is a statement about the contrast between the probes rather than about the power to detect one.

Bootstrap intervals over proteins separate the reagent’s contribution from the site’s (Figure 6C and Figure 6D). Cationic attraction is the reagent: the neutral probe’s interval brackets zero and excludes the anionic probe’s estimate. Acidic avoidance, predicted pK_a_ and salt bridges are the site, with the two probes indistinguishable on all three. Net local charge is mixed, and it is the one feature for which a partial attribution is the honest reading. Reagent attribution is not inferred from one probe reaching significance while the other does not, because a difference between significant and non-significant is not itself a significant difference, which is why the panels show intervals rather than asterisks.

This changes how an earlier reading of the same data has to be worded. Labeled lysines sit in cationic, acid-poor environments with fewer salt bridges and lower predicted pK_a_, and the natural reading of that is nucleophilicity. The decomposition says the reading is right for the pK_a_ and salt-bridge terms and wrong for cationic attraction, which tracks the reagent. Both effects are real; they are not the same effect.

Concavity, by contrast, belongs to the site. Labeled lysines sit in more enclosed environments than unlabeled ones, and the two probes do not differ on it, so this is a property of where reactive lysines are and not of what reaches them (Figure 6E). Two bounds belong with that result. Enclosure and relative solvent accessibility share about 74% of their variance across the 167,057 sites, so the concavity effect is largely, though not wholly, a solvent-accessibility effect (Figure S8B); and the effect is reported within protein, because pooling across sites reverses its sign (Figure S8C).

One assumption bounds the charge result rather than threatening it. The neutral probe carries hydrogens α to an electron-withdrawing acylsulfonamide and may enolize at physiological pH, acquiring partial anionic character; its pK_a_ is not known. If it does, the two probes are less different than assumed, and the measured reagent effect is an underestimate: it rises from the neutral-assumption value to +0.46 if the probe is substantially ionized, which is a sensitivity range and not a confidence interval (Figure S8D). The assumption cannot explain the result away.

A chemoproteomic probe is not a neutral window onto lysine reactivity: part of what a lysine-directed experiment measures is where its reagent is drawn. Coverage and reactivity are entangled at the point of capture, so a probe panel is a sampling variable to be varied deliberately rather than held fixed by convention.

### Acquired lysines are structurally ordinary

The acquired lysines of Figure 1 have been tested against nothing. Do they sit inside the same population as the lysines these rules were measured on, or does mutation create lysines in environments unlike the ones the atlas has sampled?

The comparison had to be anchored before it could be made. Our first version of it measured union positions at Cβ and measured lysines at Nζ, and returned a separation that would have been the strongest structural result in the paper. It is an artifact: Cβ and Nζ lie about 6.4 Å apart in an extended lysine side chain, so the two populations were being measured from different atoms. Figure 7A shows the artifact beside its correction, and three legitimate anchorings agree with each other (Figures S9A and S9B); the mismatched arm is retained there as a negative control, because it is the analysis we first ran. Where side chains had to be built, all-trans rotamers reproduce the amine positions of 30,805 measured lysines to a median of 4.26 Å, and all-trans is the weakest possible construction, so that is an upper bound on the error rather than an estimate of it (Figure S9E).

**Figure 7.**
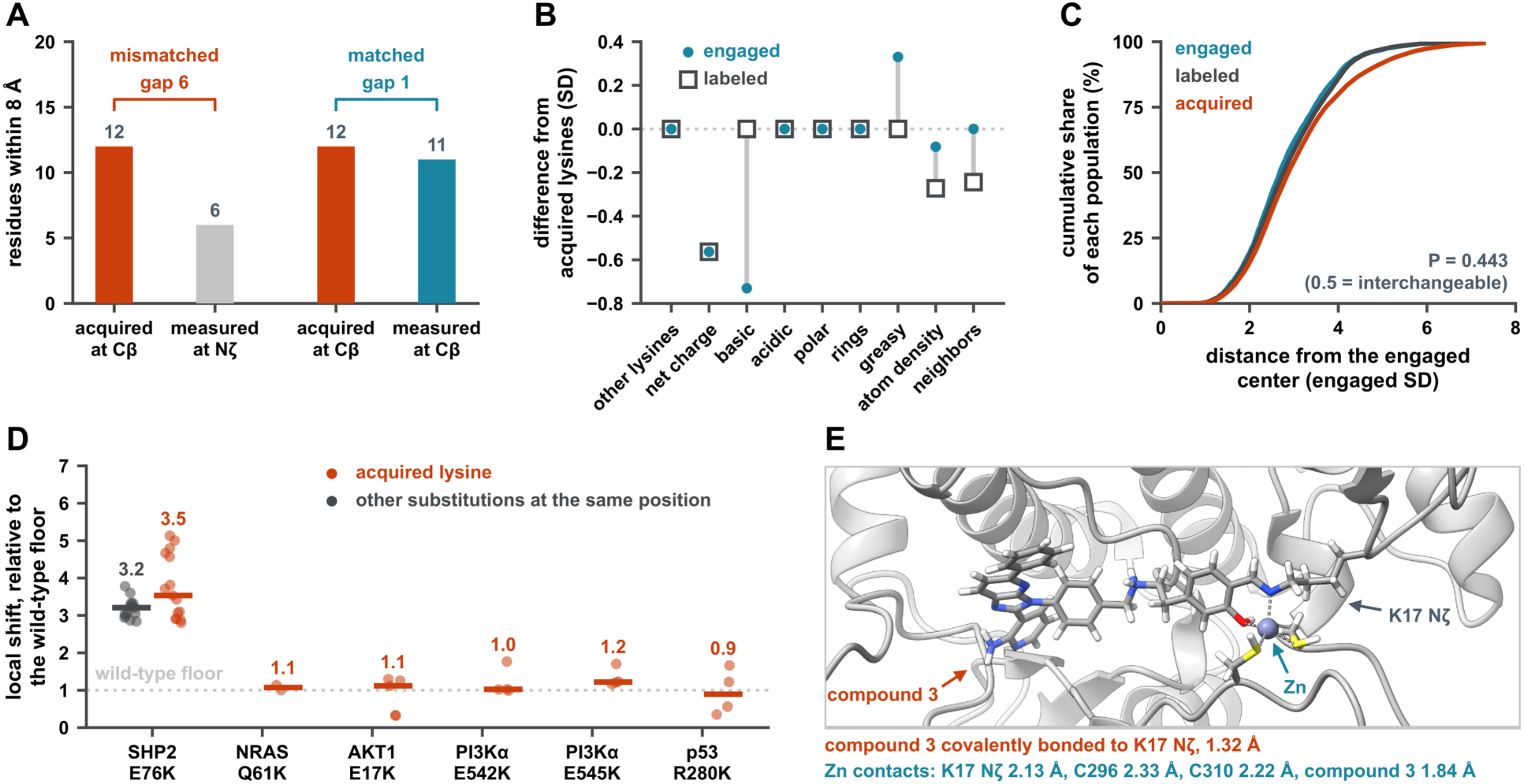
Acquired lysines are structurally ordinary. Nine environment features at 156,781 union positions with an AlphaFold confidence of at least 70^21,22^, against the engaged and unengaged lysines measured in the three screens^2–4^ (Methods). Engagement is expressed as % occupancy. **(A)** One reference atom, on both sides. The panel shows the artifact and the correction. Values in Table S2. **(B)** The three populations differ far less than the anchor artifact implied. Feature medians of engaged lysines (teal circle) and of lysines that were labeled but not engaged (open square), each drawn as its difference from the acquired median in standard deviations of the acquired distribution, so the dotted line at zero is the acquired set; a gray bar joins the two markers where they differ. Four of the nine features have identical medians across all three populations, and a fifth, neighbor count, is identical between the acquired and engaged sets; the largest difference is 0.73 standard deviations of the acquired distribution. **(C)** The distance between acquired and engaged lysines, against permutation nulls. Reported threshold-free: the probability that a randomly chosen acquired lysine sits closer to the engaged center than a randomly chosen engaged lysine does, against 0.5 for indistinguishable populations. The populations overlap heavily. **(D)** Crystallographic validation at six sites. Local Cα RMSD in a co-local shell, each mutant normalized to its own protein’s wild-type floor. Five of six are at that floor (0.89–1.22); SHP2 E76K is the exception at 3.53, and is the only site carrying a substitution series, where other substitutions at the same position give 3.21. Six sites were defined and all six measured. The NRAS wild-type set is restricted to a single nucleotide state so that every pair is state-matched, which leaves its floor resting on a single pair; the per-site floor counts are in the panel’s deposited table. The PI3Kα mutant structures 8GUA and 8GUD are cryo-EM. These are mutation sites from variant calling, not chemoproteomic assignments, so the provisional-assignment caveat does not apply to them. Structures used, all previously deposited and none determined here. PTPN11: 2SHP, 4DGP, 4DGX, 4H1O, 5EHR (wild type), 6CMS, 6CRF, 6CRG (E76K), 7EMN (E76A), 6CMR (E76D), 5IBS (E76Q); TP53: 2OCJ, 1TSR, 3KMD, 2AC0 (wild type), 6FF9 (R280K); AKT1: 1UNQ, 1H10, 1UNR, 1UNP (wild type), 2UZS, 8UVY (E17K); PIK3CA: 4OVU, 2RD0, 4L23, 5DXT (wild type), 8GUA (E542K), 8GUD (E545K); NRAS: 5UHV, 8TBI (wild type), 8VM2 (Q61K). **(E)** AKT1 E17K, engaged selectively in cells by others, and undetected here. Site geometry from PDB 8UVY, 54 atoms in the local shell. Searched as the three peptides the databases carried, two spanning the new lysine and one immediately downstream. The one model carrying the mutation that is in the proteomic set is KU-19-19 (RRID:CVCL_1344). The point is the contrast between what is chemically reachable and what this workflow can see. The engaged and unengaged comparators come from the three chemoproteomic screens (the lysine-targeting compound screen in Ramos and MDA-MB-231^2^; the meroterpenoid screen in MDA-MB-231 and MDA-MB-436^3^; the CyMPSA screen in HeLa^4^). Nothing in this figure is a statement about a tumor or a patient. See also Figures S9 and S10 and Table S1.

On a common anchor the three populations are difficult to tell apart. Four of the nine features have identical medians across acquired, engaged and unengaged lysines, and a fifth is identical between acquired and engaged sites (Figure 7B). Where the populations do differ they differ by less than one standard deviation of the acquired distribution. Reported without a threshold, the probability that a randomly chosen acquired lysine lies closer to the center of the engaged population than a randomly chosen engaged lysine does is 0.443, against 0.5 for populations that cannot be distinguished at all (Figure 7C). The reading does not depend on where a boundary is drawn: at the 50, 75, 90 and 95% engaged regions acquired lysines sit inside the engaged region less often than the labeled-but-unengaged control does, at every cut and under either anchoring and by 3.9 to 11.5 percentage points, so acquired lysines fall below both comparators rather than between them (Figure S9F).

Six of these sites have crystal structures, and they behave the same way. Local Cα displacement in a co-local shell, normalized to each protein’s own wild-type floor, puts five of the six at that floor; SHP2 E76K is the exception, and it is the one site carrying a substitution series, where other substitutions at the same position perturb the environment as much (Figure 7D).

Acquired lysines are not structurally unusual, which removes the cheapest reason to expect the ligandability rules measured in Figures 4 to 6 not to reach them. It does not establish that they can be engaged. Eight of the nine features compared here are among the fourteen from which the composite model of Figure 5 was built, every one except the count of neighboring lysines, and that model reaches an AUC of 0.617 against a measured reproducibility ceiling of at least 0.72. A feature set that predicts ligandability this weakly cannot support the reverse inference, from similarity to ligandability. The engaged and unengaged comparators come from the three chemoproteomic screens and describe those cells, not a tumor and not a patient. What this comparison supports is narrower and more useful: nothing about the chemistry of these sites explains why they are absent from the record. The barrier is detection.

One acquired lysine makes that concrete. AKT1 E17K, the transforming pleckstrin-homology-domain mutation identified in breast, colorectal and ovarian cancer^23^, has since been engaged selectively in cells through lysine targeting and neo-zinc chelation^24^, and it returns zero spectra here (Figure 7E). That negative rests on a single experiment: of the five DepMap models carrying the mutation, one is in the 375-line proteomic set, so its plex is the only one of the 42 in which the mutation was present to be found. It is one informative observation, not a survey. The one acquired lysine that has been drugged is the one this workflow did not see.

## Discussion

Mutation-created lysines are numerous, largely absent from common germline variation, enriched on driver hotspots, written by identifiable mutational processes, and almost entirely missing from the proteomic record. Every one of those is measured above, and the error rate that would license even the few identifications recovered does not hold at the reported filter.

The closest prior work approached this from two directions, and neither reaches the question asked here. The three chemoproteomic screens reanalyzed in this paper mapped ligandable lysines across several proteomes^2–4^, building on the reactivity and ligandability survey established earlier^1^, but they searched reference databases, so a mutation-created lysine was outside their search space by construction. From the other direction, AKT1 E17K has been engaged selectively in cells through lysine targeting and neo-zinc chelation^24^: an acquired lysine drugged by target-based design, one site at a time, without any claim about the class it belongs to. The measurability question has been asked and addressed for cysteine, where a chemoproteogenomic pipeline with variant-aware databases and staged error control cataloged acquired cysteines directly^7^. Coverage itself has been audited there too, by pairing predicted side-chain accessibility with a meta-analysis of eighteen public cysteine profiling datasets, which found the sampled residues biased by warhead chemistry and concluded that current approaches reach only a small fraction of the ligandable population^25^. Lysine has had no equivalent, and it is the harder case for a specific reason: an acquired cysteine leaves the tryptic peptide it sits in intact, while an acquired lysine creates a cleavage site and produces a different peptide altogether.

One of those obstacles is partly dissolved by the experiment that would be used to look for it. An acquired lysine creates a tryptic cleavage site in a shotgun experiment but not in a lysine-chemoproteomic one, because the probe acylates the ε-amine and trypsin does not cleave after an acylated lysine, so a labeled acquired lysine leaves the peptide boundaries near wild-type while the peptide is streptavidin-enriched.

The passenger reading of this class deserves a direct answer. Glutamate to lysine is the commonest substitution in cancer for reasons that carry no biological weight: glutamate is abundant, the dominant mutational process is C>T, and against a trinucleotide-context expectation the substitution is enriched only 1.12-fold, against more than threefold for each of the CpG hotspots. It would be reasonable to conclude from that alone that these are incidental residues. One observation argues otherwise: acquired lysines carry the driver-hotspot flag at an odds ratio of 1.68 against other substitutions, and that flag is conservative by construction, since the method behind it exists to strip recurrent passenger sites from hotspot lists^16^. The signature composition of those hotspots is not independent support. It compares lysine-creating hotspots against other hotspots, and since lysine is made 85.5% by C>T and never by T>C, any lysine-creating set would show the same enrichment whether or not it were under selection. What the odds ratio establishes is that these sites are selected more often than other substitutions; it does not establish that the affected proteins are expressed, detectable, or reachable by a molecule.

Which cells carry the class is not arbitrary, and it can be chosen. The four cell lines the three screens ran in carry 56 unique acquired lysines between them, against a maximum of 623 across 1,954 DepMap models, and ranked on the sites a screen could actually reach the leading lines carry 59 to 83 times what HeLa does. Which cells those are, which sites recur across models, how far burden survives gating to expression, protein detection and dependency, and what is independently known about the recurrent driver-flagged sites are set out in Extended Results with the per-line and per-site tables.

A probe panel is a sampling variable rather than a fixed choice, and that is as far as these data reach. Establishing which panel would maximize coverage requires prospective experiments a reanalysis cannot contain, and a two-probe comparison supports the principle without licensing a design.

What would actually recover this class is not what intuition suggests. Deeper or enrichment-based proteomics addresses abundance, and abundance does not look like the limiting gate: it separates sites whose mutant peptide was found from those whose region was never covered by Cliff’s δ = 0.31, on the 1,769 of 8,470 observations that carry an abundance value and only 40 found sites. A subset of the failures is instead associated with long tryptic peptides, the two distributions diverging in the tail beyond twenty residues rather than shifting as a whole, and no effect size is available for that comparison, so neither test is decisive. What can be said is that changing which proteins are sampled leaves the peptide boundaries untouched, and that three measured results point at the same lever. Half this class has no fully cleaved mutant peptide above the seven-residue floor the searches enforce, so its only searchable species carries a missed cleavage. An internal missed cleavage is the strongest negative predictor of identification measured here, at a standardized effect of −0.621. And the cleavage that fails is specifically at arginine, 29.4% of identified peptides whose first basic residue is arginine carrying a missed cleavage against 15.0% for lysine. What those three identify is completeness at the lysine and arginine sites trypsin already targets, not a different specificity: a protease cutting at additional residues would shorten peptides further, and short is where this class already fails. Digesting the same proteome more completely is a prediction this paper does not test.

The limitations are specific, and several of them bound results this paper otherwise leans on. Everything measured in cells is measured in cell lines, 375 in the proteomic re-search and four across the three chemistry screens, and no result is a statement about a tumor or a patient. Which lines carry the class, and what a re-search of the three screens could reach, are in Figure S10. Figure 1 is a census of reported variants and a population frequency comparison rather than a measurement in any cell. The screens are not saturated, so every ligandability figure is a lower bound: a site recorded as unengaged was unengaged by the compounds tested. The structural separation of reactivity from ligandability emerges progressively as the confidence gate tightens and is strongest above pLDDT 90, which makes it a claim about ordered protein, and AlphaFold confidence is not random with respect to biology. The compound descriptors tested encode neither charge nor chirality, so the finding that none of them predicts which lysines a compound prefers is silent on exactly the two axes this paper shows to matter. The nine environment features of the structural comparison describe a site’s surroundings within 8 to 10 Å; the lysine’s own Nζ chemistry is not compared, and at most acquired positions that Nζ is a built all-trans rotamer carrying a median error of 4.26 Å against a reference that is itself predicted, which is why the six-structure check matters, and that check crosses a method boundary at two PI3Kα pairs and rests on a single wild-type pair at NRAS. The cross-dataset version of the breadth result is the weakest evidence in the paper, 25 sites in 21 dimensions with two of six comparisons short of their null, and is reported as a limit rather than as support. And the detections themselves are thin: of 53, none rests on more than one experiment, which is why the error-rate audit exists and why the coverage counts are a lower bound rather than a catalog of sites.

Taken together the results describe a class that is large, selected more often than other substitutions, chemically unremarkable and effectively unmeasured, and they locate the obstacle precisely. It is not that acquired lysines are structurally unusual: nine environment features place them within the ordinary range of the lysines chemoproteomics already engages. The obstacle is not the chemistry either, since one acquired lysine, AKT1 E17K, has already been engaged selectively in a campaign built around that single site^24^, although success at one site does not make the class tractable. It is that the same site returns zero spectra in the one plex of the 42 whose cell line carries it, and that a class which cannot be seen cannot be prioritized or argued about with data. Screening does not require proteomic visibility, as the three reanalyzed screens show; knowing which sites exist, in which cells, and with what confidence does. The measurement problem is the tractable one.

## Resource availability

### Lead contact

Requests for further information should be directed to the lead contact, Mikail E. Abbasov.

### Materials availability

This study generated no new unique reagents. The lysine-directed probes and compounds reanalyzed here were reported in the three source studies and are available from their corresponding authors.

### Data and code availability

This study generated no new primary mass-spectrometry data. The proteomic data reanalyzed here are the CCLE proteomes at MassIVE MSV000085836, and the three chemoproteomic screens are available from their original publications. Variant calls are from DepMap Public 26Q1, the TCGA MC3 public call set (v0.2.8-PUBLIC) and COSMIC Cancer Mutation Census v103; sequences are MANE v1.2 and GENCODE v38; population frequencies are gnomAD v4.1 and the gnomAD v2.1.1 non-cancer exome subset; structures are AlphaFold DB. The acquired-lysine site catalog, the derived site-level tables, the panel-level data behind every figure, the mutation-aware database construction, the entrapment analysis and every figure script are deposited at github.com/Mikail-Abbasov/acquired-lysines, and every figure panel can be redrawn from the deposited data. Any additional information required to reanalyze the data reported in this paper is available from the lead contact on request.

## Acknowledgments

This work was supported by the National Institute of General Medical Sciences of the National Institutes of Health under award R35GM143027, and by a University of Oklahoma Health Sciences Center College of Pharmacy Seed Grant, PHR20102.

## Author contributions

**CRediT authorship contribution statement.** R.M.: Data curation, Investigation, Writing – review & editing. S.T.: Data curation, Investigation, Writing – review & editing. A.M.: Visualization, Writing – review & editing. M.K.H.: Investigation, Writing – review & editing. M.E.A.: Conceptualization, Methodology, Software, Formal analysis, Validation, Visualization, Writing – original draft, Supervision, Project administration, Funding acquisition.

## STAR Methods

### Key resources table

| **REAGENT or RESOURCE** | **SOURCE** | **IDENTIFIER** |
|---|---|---|
| **Deposited data** |  |  |
| CCLE quantitative proteomes (42 TMT plexes, 375 cell lines) | Nusinow et al.<sup>11</sup> | MassIVE: MSV000085836 |
| DepMap Public 26Q1 (OmicsSomaticMutations, Model, CRISPRGeneEffect, CRISPRInferredCommonEssentials, OmicsExpressionTPMLogp1HumanProteinCodingGenes) | DepMap<sup>12</sup> | depmap.org/portal/download |
| TCGA MC3 somatic mutation calls, v0.2.8-PUBLIC | Ellrott et al.<sup>13</sup> | gdc.cancer.gov/about-data/publications/mc3-2017 |
| COSMIC Cancer Mutation Census v103 | Tate et al.<sup>14</sup> | cancer.sanger.ac.uk/cmc |
| gnomAD v4.1 joint allele frequencies | Chen et al.<sup>15</sup> | gnomad.broadinstitute.org |
| gnomAD v2.1.1 non-cancer exome subset | Karczewski et al.<sup>26</sup> | gnomad.broadinstitute.org |
| dbNSFP 5.4a | Liu et al.<sup>27</sup> | database.liulab.science/dbNSFP |
| MANE Select v1.2 | NCBI and EMBL-EBI | ncbi.nlm.nih.gov/refseq/MANE |
| GENCODE v38 | GENCODE | gencodegenes.org/human/release_38.html |
| UniProt | UniProt | uniprot.org |
| AlphaFold Monomer v2.0 models | AlphaFold Protein Structure Database<sup>21,22</sup> | alphafold.ebi.ac.uk |
| Lysine-targeting compound screen (Ramos, MDA-MB-231): site occupancies and compound structures | Abbasov et al.<sup>2</sup> | Supporting information of the source study |
| Meroterpenoid screen (MDA-MB-231, MDA-MB-436): site occupancies and compound structures | Bracken et al.<sup>3</sup> | Supporting information of the source study |
| CyMPSA screen (HeLa): site occupancies and compound structures | Ryan et al.<sup>4</sup> | Supporting information of the source study |
| PTPN11 structures | Protein Data Bank | PDB: 2SHP, 4DGP, 4DGX, 4H1O, 5EHR, 6CMS, 6CRF, 6CRG, 7EMN, 6CMR, 5IBS |
| TP53 structures | Protein Data Bank | PDB: 2OCJ, 1TSR, 3KMD, 2AC0, 6FF9 |
| AKT1 structures | Protein Data Bank | PDB: 1UNQ, 1H10, 1UNR, 1UNP, 2UZS, 8UVY |
| PIK3CA structures | Protein Data Bank | PDB: 4OVU, 2RD0, 4L23, 5DXT, 8GUA, 8GUD |
| NRAS structures | Protein Data Bank | PDB: 5UHV, 8TBI, 8VM2 |
| Analysis and figure scripts, panel-level data for every figure, the acquired-lysine site catalog and site-level tables | This paper | github.com/Mikail-Abbasov/acquired-lysines |
| **Software and algorithms** |  |  |
| MSFragger 4.4.1 | Kong et al.<sup>28</sup> | msfragger.nesvilab.org |
| FragPipe v24.0 | Nesvizhskii laboratory | fragpipe.nesvilab.org |
| Philosopher 5.1.3-RC9 | da Veiga Leprevost et al.<sup>29</sup> | github.com/Nesvilab/philosopher |
| Percolator 3.7.1 | Käll et al.<sup>30</sup> | github.com/percolator/percolator |
| FDRBench v1.1.1 | Wen et al.<sup>10</sup> | github.com/Noble-Lab/FDRBench |
| fpocket 4.2.3 | Le Guilloux et al.<sup>31</sup> | github.com/Discngine/fpocket |
| PROPKA 3.5.1 | Olsson et al.<sup>19</sup> | github.com/jensengroup/propka |
| Python 3.10 | Python Software Foundation | www.python.org |
| NumPy 2.2.6 | Harris et al.<sup>32</sup> | numpy.org |
| pandas 2.3.3 | McKinney<sup>33</sup> | pandas.pydata.org |
| SciPy 1.15.3 | Virtanen et al.<sup>34</sup> | scipy.org |
| scikit-learn 1.7.2 | Pedregosa et al.<sup>35</sup> | scikit-learn.org |
| statsmodels 0.15.0 | Seabold and Perktold<sup>36</sup> | www.statsmodels.org |
| Matplotlib 3.10.9 | Hunter<sup>37</sup> | matplotlib.org |
| Biopython 1.88 | Cock et al.<sup>38</sup> | biopython.org |
| FreeSASA 2.2.1 | Mitternacht<sup>39</sup> | freesasa.github.io |
| RDKit 2026.03.5 | Landrum et al.<sup>40</sup> | www.rdkit.org |
| python-docx 1.2.0 | python-docx | github.com/python-openxml/python-docx |

### Method details

#### Variant catalogs and the union

Missense sites were pooled from three sources: DepMap release 26Q1^12^, the TCGA MC3 public call set (v0.2.8-PUBLIC)^13^, and COSMIC Cancer Mutation Census v103^14^. The union holds 4,395,403 distinct missense sites, keyed on gene, position, reference residue and alternate residue, of which 295,411 substitute a lysine. A site is counted once however many sources report it, and 93,769 are reported by more than one. Two TCGA cohorts, ovarian serous cystadenocarcinoma and acute myeloid leukemia, were generated from whole-genome-amplified material; a PASS-only filter removes them entirely, so filtering was applied without that restriction and the two cohorts are retained. Driver-hotspot and mutational-signature flags are those carried in the DepMap variant table, derived by the method of Hess et al.^16^; no gene list was chosen for this work.

#### Germline comparison

Germline rates were computed from dbNSFP 5.4a^27^, restricted to rows carrying a MANE annotation and an unambiguous single-letter amino-acid change. The reported figure is an expected diploid count per individual, the sum of twice the allele frequency over qualifying rows, and is not a count of distinct variants. Allele frequencies are gnomAD v4.1 joint frequencies, approximately 807,000 individuals^15^; 91,571,221 rows were scanned and 14,084,288 carried a frequency above zero. The control repeats the same calculation over the gnomAD v2.1.1 non-cancer exome subset, approximately 118,000 individuals^26^. The two cohorts overlap heavily and are not independent samples of human variation; the control exists to remove one specific objection, that cancer-affected individuals inflate the germline baseline.

#### Proteomic reanalysis

The 42 sample plexes of the CCLE proteomes^11^ (nine cell lines per plex and a pooled bridge in the tenth channel, 378 samples in all, three lines run twice, so 375 distinct cell lines) were re-searched with MSFragger 4.4.1 within FragPipe v24.0, with Percolator and Philosopher. Two architectures were run. The conventional search used a 20,431-protein MANE v1.2 database with contaminants and reversed decoys. The mutation-aware search used a peptide-level database of 10.17 million entries built from the variant union. Detection was scored at two gates that are never combined: gate 1 asks whether the wild-type region was observed at all, gate 2 whether the mutant peptide was recovered where it was. The unit throughout Figures 2 and 3 is a site-in-cell-line observation, not a distinct genomic site: 8,470 acquired-lysine observations cover 8,055 distinct sites across the 372 lines carrying one, in 10,372 rows, with 1,902 non-lysine variant observations carried through the identical pipeline as a control. Figure 1’s 295,411 is a distinct-site count and is a different denominator. Note N9 explains why PgxSAVy was not used here.

#### Entrapment and error control

Realized error rates for variant peptides were measured by burden-matched entrapment at r = 1 using FDRBench v1.1.1^10^. The prior work this design answers to is summarized in note N10. Each variant peptide was paired with an entrapment sequence built by permuting its own residues with the C-terminal lysine or arginine held in place, so the pair shares an exact precursor mass and an identical length and only fragment ions distinguish them; 589,588 variant-pair rows across eight plexes yielded entrapment sets of 147,397 each, all sharing residues with their target. Because every entrapment identification is false by construction, the proportion of entrapment hits estimates the realized false-discovery proportion directly, without the distributional assumptions target-decoy control requires^9^. The same proportion measured on acquired-lysine peptides alone is in note N11. The score margin is the difference between a spectrum’s best and second-best interpretation; margin ≥ 10 is used throughout. Two denominators are in use and are not interchangeable: Figure 3B counts distinct peptides, one best spectrum each, and gives 0.83% at that margin; Figure 3C and Figure S4E count identifications and give 0.58%. The proportion reported throughout is e/(t+e), entrapment hits over all hits. The r = 1 correction 2e/(t+e), which assumes a false match is equally likely to land on a target as on an entrapment sequence, is not used here: the entrapment partner shares its target’s exact mass, length and residues, so it competes harder than a random target would, which makes e/(t+e) an upper bound rather than the lower bound the survey warns against^10^. pooled_fdp.py prints the corrected form alongside; the two differ by exactly a factor of two.

#### Site assignment

Chemoproteomic site assignments in Figures 4 to 6 were reconciled between two independent routes, which concur at 98.21% (44,736 of 45,552 usable sites; note N1). Assignments are described as provisional throughout: the figures compare site *sets*, and an individual residue number within a set may be wrong even where the set is right. Figure 7 is the exception, since its sites come from variant calling rather than from chemoproteomic assignment.

#### Occupancy and engagement

Occupancy is the share of probe labeling a compound competes away, 100 × (1 − compound / DMSO) (note N2). A site is engaged at ≥ 60% occupancy; screening depth differs between the screens (note N3). Where source screens report competition ratios rather than occupancies, a ratio of 4 corresponds to 75% engagement; the 60% cut is a reporting convention shared with those screens rather than a boundary in the data, and Figure 4F and Extended Results, together with Figure S7C, show the results do not depend on it.

#### Structural features

Site features were measured on AlphaFold Monomer v2.0 models retrieved from the AlphaFold Protein Structure Database^21,22^, files tried at v6, v5 and v4. Positions are anchored at Cβ unless stated; where an Nζ is required at a position that carries no lysine in the model it is built as an all-trans rotamer, which reproduces measured amine positions to a median 4.26 Å and is an upper bound rather than a placement. Relative solvent accessibility follows Tien et al.^18^, predicted pK_a_ is from PROPKA3^19^ and pocket descriptors from fpocket^31^. Four distinct feature sets are used. Figure 4 pairs 15 features across both contrasts, 6 charge and 9 shape, and excludes the AlphaFold terms, because confidence is the gating variable there rather than a site property. Figure 5’s composite is 14 features: 13 environment features plus model confidence, and the composite is reported both with it (AUC 0.617) and without it (0.611). Figures 5B and 5E use a different set of 14 site features, extended to 15 with pocket occupancy in Extended Results and note N5, so those counts are not Figure 4’s. Figure 6 measures 22 site features and Figure 7 nine. The nine features compared at union positions describe a site’s surroundings within 8 to 10 Å: neighbor count, atom density, hydrophobic, aromatic and polar contacts, acidic and basic residues, net local charge, and other lysines. Three gate rungs are in use: Figures 4D and 4E run the ladder at none, ≥ 50, ≥ 70 and ≥ 90 and Extended Results bands confidence at the same rungs; Figures 4B and 4C display pLDDT ≥ 90; Figures 6 and 7 are gated at ≥ 70. Every structural gate is applied to both groups of a contrast at once, since gating one side would manufacture a difference rather than reveal one. The crystal structures compared in Figure 7D and their superposition are described in note N8.

#### Software

Analysis ran under Python 3.10 with numpy 2.2.6^32^, pandas 2.3.3^33^, scipy 1.15.3^34^, scikit-learn 1.7.2^35^, statsmodels 0.15.0^36^ and matplotlib 3.10.9^37^. AlphaFold models are parsed with Biopython 1.88^38^, which also computes solvent-accessible surface areas by the Shrake–Rupley method. FreeSASA 2.2.1^39^ provides an independent check of those areas (r = 0.9957), and compound descriptors are computed with RDKit 2026.03.5^40^. Peptide identification used MSFragger 4.4.1^28^ within FragPipe v24.0 with Percolator 3.7.1^30^ and Philosopher 5.1.3-RC9^29^, entrapment databases were built with FDRBench v1.1.1, pocket descriptors come from fpocket 4.2.3^31^ and predicted pK_a_ from PROPKA 3.5.1^19^. Manuscript and Supporting Information are assembled from the same scripts with python-docx 1.2.0, so every number in the text is printed by the analysis rather than transcribed into it.

#### Cell lines

Figures 2 and 3 are the CCLE proteomics reanalysis across 375 cell lines. Figures 4 to 7 are the three published chemoproteomic screens, in four cell lines: CyMPSA in HeLa (RRID:CVCL_0030)^4^; the meroterpenoid screen in MDA-MB-231 (RRID:CVCL_0062) and MDA-MB-436 (RRID:CVCL_0623)^3^; and the lysine-targeting compound screen in Ramos (RRID:CVCL_0597) and MDA-MB-231^2^. Compounds were applied to intact cells in the CyMPSA and meroterpenoid screens and to cell lysate in the lysine-targeting compound screen; in all three, probe labeling was performed on lysate after compound treatment, so the readout is identical across screens and they differ only in where the compound encountered the proteome (note N4). No cells were cultured for this work.

#### Code and reproducibility

No new data were generated. Every analysis and figure script is deposited with its version and random seed, and every number in this paper is printed by the script that produces the corresponding panel rather than transcribed from the artwork.

#### Data sources and versions

##### No new data were generated

Every source is public and is listed with its version below; each is also recorded in the deposit’s external_inputs.json, which the release build refuses to assemble if any input is undeclared.

##### Variant catalogs

DepMap Public 26Q1 (depmap.org/portal/download), comprising OmicsSomaticMutations, Model, CRISPRGeneEffect, CRISPRInferredCommonEssentials and OmicsExpressionTPMLogp1HumanProteinCodingGenes. TCGA MC3 v0.2.8-PUBLIC (gdc.cancer.gov/about-data/publications/mc3-2017). COSMIC Cancer Mutation Census v103 (cancer.sanger.ac.uk/cmc); COSMIC requires a license for commercial use, which this deposit does not confer.

##### Population frequency and reference sequence

dbNSFP 5.4a; gnomAD v4.1 joint and v2.1.1 non-cancer exomes (gnomad.broadinstitute.org); MANE Select v1.2 (ncbi.nlm.nih.gov/refseq/MANE); UniProt and AlphaFold DB (uniprot.org, alphafold.ebi.ac.uk).

##### Proteomes

The 42 TMT plexes reanalyzed here are the published CCLE proteomes of Nusinow et al.^11^; raw files were obtained from that study’s repository and re-searched with MSFragger 4.4.1, FragPipe v24.0, Percolator and Philosopher. Entrapment sets were built with FDRBench v1.1.1.

##### Chemoproteomic screens

Occupancy and compound structures were taken from the supporting information of the three published screens^2–4^. No new chemoproteomic experiments were performed.

##### Code and deposited data

All analysis and figure scripts, the panel-level data behind every figure, the acquired-lysine site catalog with fourteen site-level tables, and the checks described below are available at github.com/Mikail-Abbasov/acquired-lysines. The code carries the MIT license and the deposited data CC BY 4.0. The deposit has two tiers. Every published panel can be redrawn from the deposited panel data alone, with no download. Those files can in turn be regenerated from the public sources above, with two exceptions that are stated rather than implied: six large intermediates that are this study’s own outputs rather than public downloads, and the variant and decoy search databases. Neither is redistributed, both are named in external_inputs.json, and the second tier is therefore incomplete for the steps that read them while the first tier covers every figure affected. Each figure script also writes a machine-readable ledger of the numbers its panels contribute to the text; two scripts check those ledgers against the manuscript in both directions, numbers computed but no longer quoted, and numbers quoted that no script produces.

## Quantification and statistical analysis

### Statistics

Models were fitted with scikit-learn^35^. The composite classifier, fitted on 30,805 lysines across 6,201 proteins, is logistic regression cross-validated by protein rather than by site, since lysines on one protein are correlated; intervals on model AUCs are the 2.5th to 97.5th percentile across 20 protein-held-out splits, and intervals on reproducibility ceilings the same percentiles across 50 random half-splits. Permutation tests in Figure 4E are exact, enumerating all C(15,6) = 5,005 ways of splitting the features. Null distributions for the breadth analyses use 2,000 label shuffles, which sets a resolution limit of p < 0.0005; the control that excludes promiscuous compounds is in note N6. Compound descriptors were computed with RDKit^40^; NPR1 and NPR2 are principal-moment ratios as defined by Sauer and Schwarz^20^. Random seeds are fixed in the deposited scripts. Statistical parameters for individual panels, including sample sizes and interval definitions where they apply, are given in the figure legends and in Table S2.

## References

[1] Hacker SM, Backus KM, Lazear MR, Forli S, Correia BE, Cravatt BF. Global profiling of lysine reactivity and ligandability in the human proteome. Nat Chem. 2017;9(12):1181–1190. doi:10.1038/nchem.2826

[2] Abbasov ME, Kavanagh ME, Ichu T-A, Lazear MR, Tao Y, Crowley VM, am Ende CW, Hacker SM, Ho J, Dix MM, Suciu R, Hayward MM, Kiessling LL, Cravatt BF. A proteome-wide atlas of lysine-reactive chemistry. Nat Chem. 2021;13(11):1081–1092. doi:10.1038/s41557-021-00765-4

[3] Bracken AK, Gekko CE, Suss NO, Lueders EE, Cui Q, Fu Q, Lui ACW, Anderson ET, Zhang S, Abbasov ME. Biomimetic synthesis and chemical proteomics reveal the mechanism of action and functional targets of phloroglucinol meroterpenoids. J Am Chem Soc. 2024;146(4):2524– 2548. doi:10.1021/jacs.3c10741

[4] Ryan EM, Norinskiy MA, Bracken AK, Lueders EE, Chen X, Fu Q, Anderson ET, Zhang S, Abbasov ME. Activity-based acylome profiling with N-(cyanomethyl)-N-(phenylsulfonyl)amides for targeted lysine acylation and post-translational control of protein function in cells. J Am Chem Soc. 2024;146(40):27622–27643. doi:10.1021/jacs.4c09073

[5] Zhou M, Li S, Tan Y, Huang W, Li Y, Yuan X, Li Z. Global profiling lysine reactivity and ligandability with oxidant-triggered bioconjugation chemistry. Angew Chem Int Ed Engl. 2025;64(6):e202418473. doi:10.1002/anie.202418473

[6] Ostrem JM, Peters U, Sos ML, Wells JA, Shokat KM. K-Ras(G12C) inhibitors allosterically control GTP affinity and effector interactions. Nature. 2013;503(7477):548–551. doi:10.1038/nature12796

[7] Desai H, Andrews KH, Bergersen KV, Ofori S, Yu F, Shikwana F, et al. Chemoproteogenomic stratification of the missense variant cysteinome. Nat Commun. 2024;15(1):9284. doi:10.1038/s41467-024-53520-x

[8] Boatner LM, Palafox MF, Schweppe DK, Backus KM. CysDB: a human cysteine database based on experimental quantitative chemoproteomics. Cell Chem Biol. 2023;30(6):683–698.e3. doi:10.1016/j.chembiol.2023.04.004

[9] Elias JE, Gygi SP. Target-decoy search strategy for increased confidence in large-scale protein identifications by mass spectrometry. Nat Methods. 2007;4(3):207–214. doi:10.1038/nmeth1019

[10] Wen B, Freestone J, Riffle M, MacCoss MJ, Noble WS, Keich U. Assessment of false discovery rate control in tandem mass spectrometry analysis using entrapment. Nat Methods. 2025;22(7):1454–1463. doi:10.1038/s41592-025-02719-x

[11] Nusinow DP, Szpyt J, Ghandi M, Rose CM, McDonald ER 3rd, Kalocsay M, et al. Quantitative proteomics of the Cancer Cell Line Encyclopedia. Cell. 2020;180(2):387–402.e16. doi:10.1016/j.cell.2019.12.023

[12] Tsherniak A, Vazquez F, Montgomery PG, Weir BA, Kryukov G, Cowley GS, et al. Defining a cancer dependency map. Cell. 2017;170(3):564–576.e16. doi:10.1016/j.cell.2017.06.010

[13] Ellrott K, Bailey MH, Saksena G, Covington KR, Kandoth C, Stewart C, et al. Scalable open science approach for mutation calling of tumor exomes using multiple genomic pipelines. Cell Syst. 2018;6(3):271–281.e7. doi:10.1016/j.cels.2018.03.002

[14] Tate JG, Bamford S, Jubb HC, Sondka Z, Beare DM, Bindal N, et al. COSMIC: the Catalogue Of Somatic Mutations In Cancer. Nucleic Acids Res. 2019;47(D1):D941–D947. doi:10.1093/nar/gky1015

[15] Chen S, Francioli LC, Goodrich JK, Collins RL, Kanai M, Wang Q, et al. A genomic mutational constraint map using variation in 76,156 human genomes. Nature. 2024;625(7993):92–100. doi:10.1038/s41586-023-06045-0

[16] Hess JM, Bernards A, Kim J, Miller M, Taylor-Weiner A, Haradhvala NJ, Lawrence MS, Getz G. Passenger hotspot mutations in cancer. Cancer Cell. 2019;36(3):288–301.e14. doi:10.1016/j.ccell.2019.08.002

[17] Takahashi M, Chong HB, Zhang S, Yang T-Y, Lazarov MJ, Harry S, et al. DrugMap: a quantitative pan-cancer analysis of cysteine ligandability. Cell. 2024;187(10):2536–2556.e30. doi:10.1016/j.cell.2024.03.027

[18] Tien MZ, Meyer AG, Sydykova DK, Spielman SJ, Wilke CO. Maximum allowed solvent accessibilities of residues in proteins. PLoS One. 2013;8(11):e80635. doi:10.1371/journal.pone.0080635

[19] Olsson MHM, Søndergaard CR, Rostkowski M, Jensen JH. PROPKA3: consistent treatment of internal and surface residues in empirical pKa predictions. J Chem Theory Comput. 2011;7(2):525–537. doi:10.1021/ct100578z

[20] Sauer WHB, Schwarz MK. Molecular shape diversity of combinatorial libraries: a prerequisite for broad bioactivity. J Chem Inf Comput Sci. 2003;43(3):987–1003. doi:10.1021/ci025599w

[21] Jumper J, Evans R, Pritzel A, Green T, Figurnov M, Ronneberger O, et al. Highly accurate protein structure prediction with AlphaFold. Nature. 2021;596(7873):583–589. doi:10.1038/s41586-021-03819-2

[22] Varadi M, Anyango S, Deshpande M, Nair S, Natassia C, Yordanova G, et al. AlphaFold Protein Structure Database: massively expanding the structural coverage of protein-sequence space with high-accuracy models. Nucleic Acids Res. 2022;50(D1):D439–D444. doi:10.1093/nar/gkab1061

[23] Carpten JD, Faber AL, Horn C, Donoho GP, Briggs SL, Robbins CM, et al. A transforming mutation in the pleckstrin homology domain of AKT1 in cancer. Nature. 2007;448(7152):439–444. doi:10.1038/nature05933

[24] Craven GB, Chu H, Sun JD, Carelli JD, Coyne B, Chen H, et al. Mutant-selective AKT inhibition through lysine targeting and neo-zinc chelation. Nature. 2025;637(8044):205–214. doi:10.1038/s41586-024-08176-4

[25] White MEH, Gil J, Tate EW. Proteome-wide structural analysis identifies warhead- and coverage-specific biases in cysteine-focused chemoproteomics. Cell Chem Biol. 2023;30(7):828–838.e4. doi:10.1016/j.chembiol.2023.06.021

[26] Karczewski KJ, Francioli LC, Tiao G, Cummings BB, Alföldi J, Wang Q, et al. The mutational constraint spectrum quantified from variation in 141,456 humans. Nature. 2020;581(7809):434–443. doi:10.1038/s41586-020-2308-7

[27] Liu X, Li C, Mou C, Dong Y, Tu Y. dbNSFP v4: a comprehensive database of transcript-specific functional predictions and annotations for human nonsynonymous and splice-site SNVs. Genome Med. 2020;12(1):103. doi:10.1186/s13073-020-00803-9

[28] Kong AT, Leprevost FV, Avtonomov DM, Mellacheruvu D, Nesvizhskii AI. MSFragger: ultrafast and comprehensive peptide identification in mass spectrometry–based proteomics. Nat Methods. 2017;14(5):513–520. doi:10.1038/nmeth.4256

[29] da Veiga Leprevost F, Haynes SE, Avtonomov DM, Chang H-Y, Shanmugam AK, Mellacheruvu D, et al. Philosopher: a versatile toolkit for shotgun proteomics data analysis. Nat Methods. 2020;17(9):869–870. doi:10.1038/s41592-020-0912-y

[30] Käll L, Canterbury JD, Weston J, Noble WS, MacCoss MJ. Semi-supervised learning for peptide identification from shotgun proteomics datasets. Nat Methods. 2007;4(11):923–925. doi:10.1038/nmeth1113

[31] Le Guilloux V, Schmidtke P, Tuffery P. Fpocket: an open source platform for ligand pocket detection. BMC Bioinformatics. 2009;10:168. doi:10.1186/1471-2105-10-168

[32] Harris CR, Millman KJ, van der Walt SJ, Gommers R, Virtanen P, Cournapeau D, et al. Array programming with NumPy. Nature. 2020;585(7825):357–362. doi:10.1038/s41586-020-2649-2

[33] McKinney W. Data structures for statistical computing in Python. Proc 9th Python Sci Conf. 2010:56–61. doi:10.25080/Majora-92bf1922-00a

[34] Virtanen P, Gommers R, Oliphant TE, Haberland M, Reddy T, Cournapeau D, et al. SciPy 1.0: fundamental algorithms for scientific computing in Python. Nat Methods. 2020;17(3):261–272. doi:10.1038/s41592-019-0686-2

[35] Pedregosa F, Varoquaux G, Gramfort A, Michel V, Thirion B, Grisel O, et al. Scikit-learn: machine learning in Python. J Mach Learn Res. 2011;12:2825–2830.

[36] Seabold S, Perktold J. Statsmodels: econometric and statistical modeling with Python. Proc 9th Python Sci Conf. 2010:92–96. doi:10.25080/Majora-92bf1922-011

[37] Hunter JD. Matplotlib: a 2D graphics environment. Comput Sci Eng. 2007;9(3):90–95. doi:10.1109/MCSE.2007.55

[38] Cock PJA, Antao T, Chang JT, Chapman BA, Cox CJ, Dalke A, et al. Biopython: freely available Python tools for computational molecular biology and bioinformatics. Bioinformatics. 2009;25(11):1422–1423. doi:10.1093/bioinformatics/btp163

[39] Mitternacht S. FreeSASA: an open source C library for solvent accessible surface area calculations. F1000Res. 2016;5:189. doi:10.12688/f1000research.7931.1

[40] Landrum G, et al. RDKit: open-source cheminformatics [computer software]. https://www.rdkit.org. Accessed 6 September 2026. doi:10.5281/zenodo.591637

