## Supporting Information for "Proteome-wide reanalysis identifies coverage and error-rate limits on acquired lysines in cancer"

### Contents

Figure S1 (Figure 1). The substitution matrix in full. .......... S2
Figure S2 (Figure 2). The non-lysine control and what limits detection. .......... S3
Figure S3 (Figure 2). Four explanations that are not the answer. .......... S4
Figure S4 (Figure 3). The entrapment design in full. .......... S5
Figure S5 (Figure 4). What the confidence gate does. .......... S6
Figure S6 (Figure 5). The ceiling, and the limit. .......... S7
Figure S7 (Figure 5). The confound, and what no descriptor can see. .......... S8
Figure S8 (Figure 6). Each probe alone, and the pooling reversal. .......... S9
Figure S9 (Figure 7). The anchor tested rather than defended. .......... S10
Figure S10 (Figure 7). The acquired-lysine class across cell lines .......... S11
Table S1. The recurrent acquired lysines .......... S13
Table S2. Panel-level values for the main figures .......... S18
Extended Results .......... S20
Supplementary notes .......... S24
N1. Site validation: two-route agreement .......... S24
N2. Occupancy scale construction across the three datasets .......... S25
N3. Screening depth by dataset .......... S26
N4. Cross-screen concordance of site ligandability .......... S27
N5. Pocket membership survives the confidence gate .......... S28
N6. Promiscuity control .......... S29
N7. Compound descriptors and 3D shape .......... S30
N8. Crystallographic validation candidates .......... S31
N9. PgxSAVy answers a different question from the one this work asks .......... S32
N10. Entrapment prior art .......... S33
N11. Class-restricted entrapment .......... S34
Supplemental References .......... S35

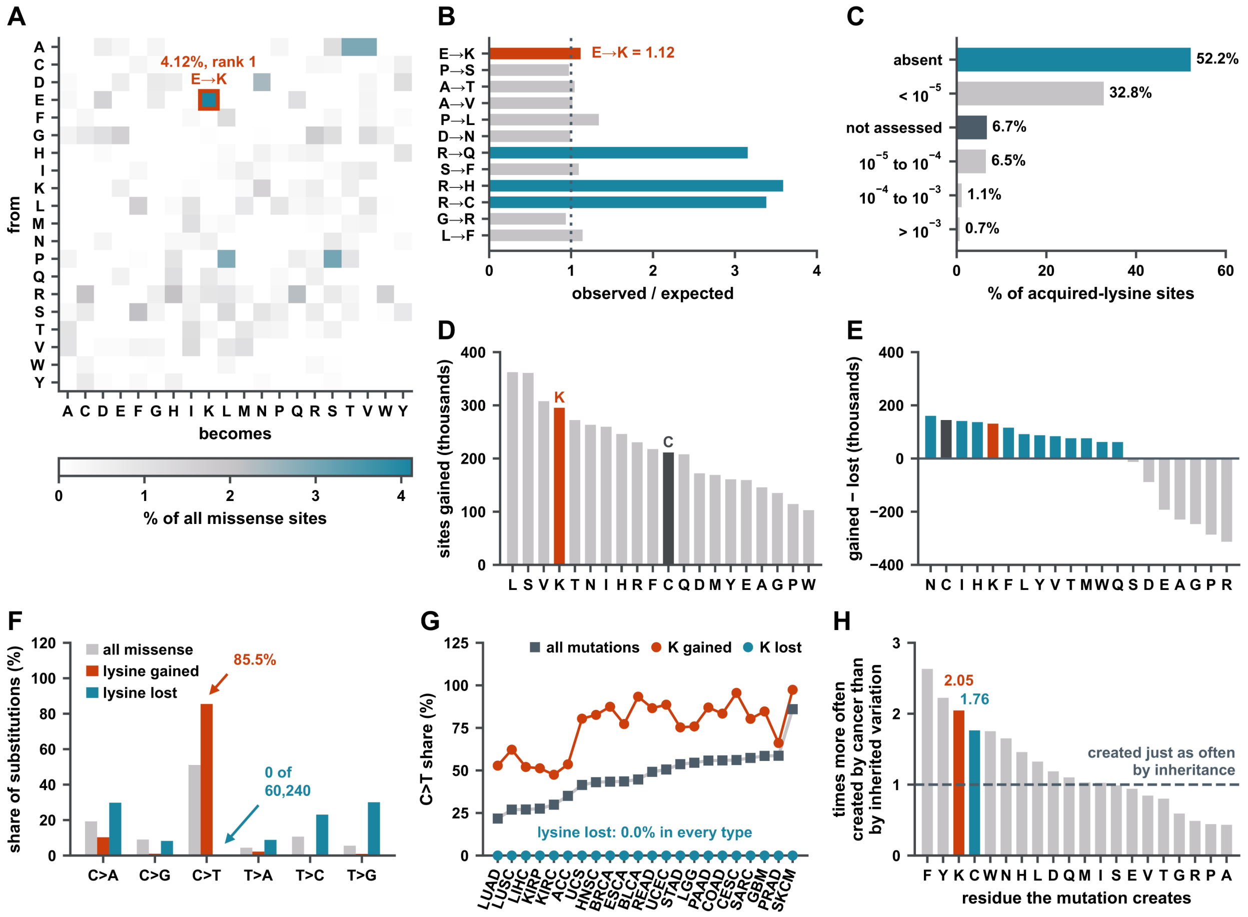


**Figure S1. The substitution matrix in full, Related to Figure 1. (A)** The complete 20 × 20 substitution matrix. Direct counts from the union, normalized over the whole matrix rather than per row, so cells are comparable across rows. The whole matrix sums to 100%, and a white cell was never observed. **(B)** Against a trinucleotide-context expectation, E→K is barely enriched. Observed over expected for the twelve commonest substitutions: E→K = 1.12, against three CpG hotspots that dominate the panel (R→H 3.59, R→C 3.38, R→Q 3.16). The expectation is built from trinucleotide context and codon usage, and R→Q, R→H and R→C are the CpG hotspots. **(C)** gnomAD frequency bands as shares. Absent 52.2%, below $10^{-5}$ 32.8%, $10^{-5}$ to $10^{-4}$ 6.5%, $10^{-4}$ to $10^{-3}$ 1.1%, above $10^{-3}$ 0.7%, not assessed 6.7%. 91.1% of assessed sites are absent or below $10^{-5}$. **(D)** Which residues are gained. Sites gained per residue across the union. Lysine is fourth at 295,411; cysteine eleventh at 211,386, so acquired lysines outnumber acquired cysteines 1.40-fold. **(E)** Net flux, gains minus losses. Lysine gains 295,411 and loses 164,237, netting 131,174 and ranking fifth. Cysteine nets 144,683 and ranks second, because cysteine is rarely lost (66,703) and so nets higher on far smaller numbers. Lysine turns over more. **(F)** Gains are one substitution class; losses are every other one. Lysine gain is 85.5% C>T and 0% T>C; lysine loss is 0% C>T and spread across the other five. All missense sits between, at 51.1% C>T. Missense only, so the three sets are like for like; nonsense adds 3,020 further losses, all T>A. **(G)** The asymmetry is not inherited from the background spectrum. The signature control, per lineage, ordered by each cancer's own C>T content, from tobacco-driven lung (LUAD, 22%) to ultraviolet melanoma (SKCM, 86%). Gains follow that ordering; losses do not, and lysine loss is 0.0% C>T in every lineage. **(H)** Every residue, not only lysine, and cysteine on the identical calculation. Somatic share of the missense burden divided by germline share, for all twenty residues, ordered by the ratio. Computed by step54_germline_by_residue.py on the same pass over dbNSFP 5.4a that produced Figure 1G: same file, same MANE restriction, same gnomAD v4.1 joint frequency field, same expected-diploid-count definition. It returns 471.1 acquired lysines per individual, which is Figure 1G's value, and that agreement is the check that generalizing the calculation did not change it. Rust, lysine at 2.05, third of twenty; teal, cysteine at 1.76, fourth. The distribution runs from 0.43 (alanine) to 2.63 (phenylalanine) with a median of 1.07, and eight residues fall below 1. The panel exists because Figure 1F shows all 32 lineages for the same reason: a single ratio cannot carry its own control. It is not a measure of selection in cancer. Germline shares are lowered by purifying selection, so residues whose introduction is most disruptive are depleted from inherited variation and score high here independently of any cancer-specific process; that is the likeliest explanation for phenylalanine and tyrosine, the only two residues above lysine. Selection is Figure 1D's odds ratio of 1.68; the context-corrected enrichment for E→K is panel B's 1.12-fold.

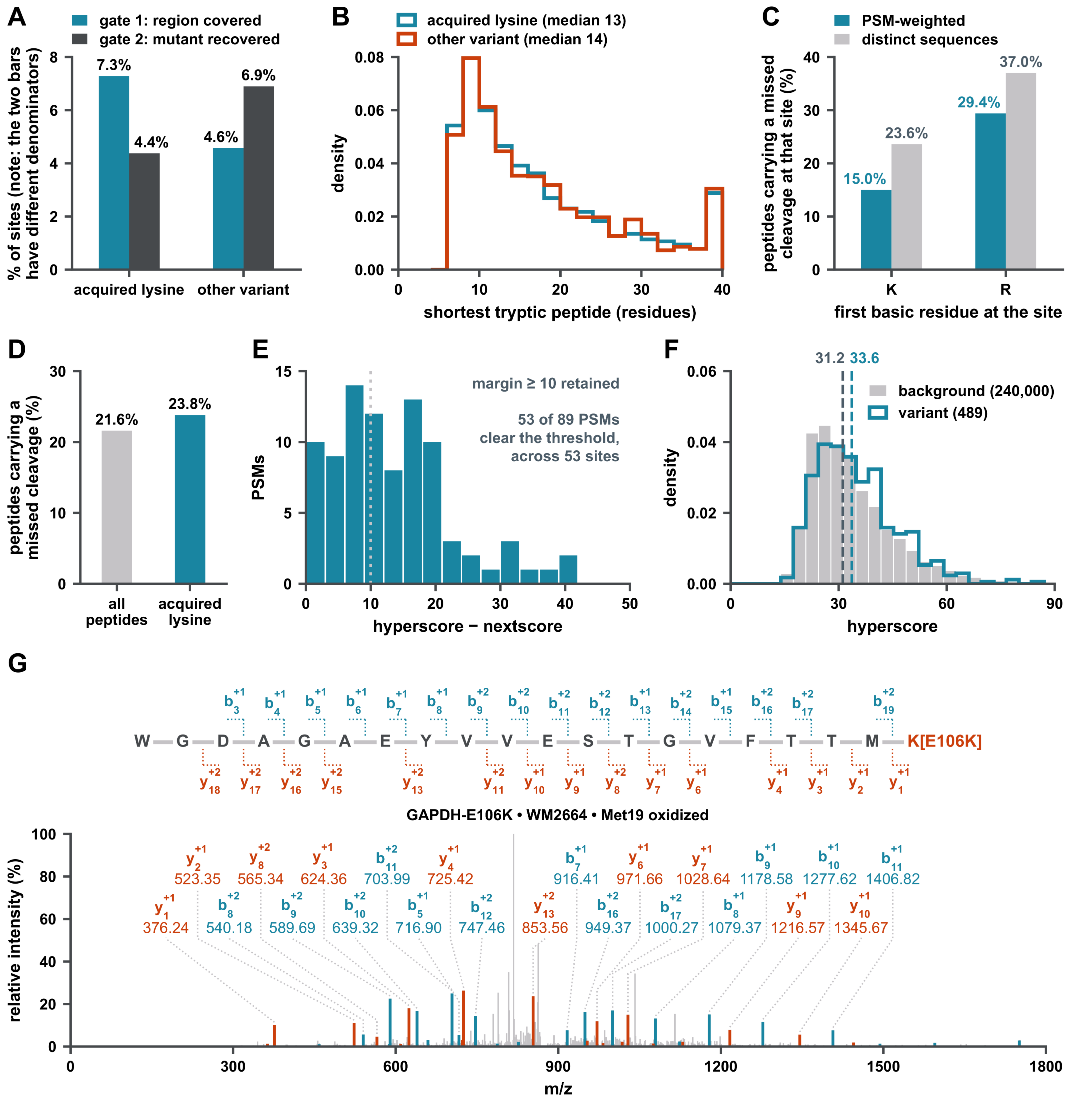


**Figure S2. The non-lysine control and what limits detection, Related to Figure 2. (A)** The two gates, both classes. 1,902 variant sites that do not create a lysine, through the identical pipeline. Acquired lysines (n = 8,470): gate 1 7.3% covered, gate 2 4.4% recovered. Other variants (n = 1,902): 4.6% and 6.9%. Acquired lysines are covered more and recovered less, because a non-lysine variant shares peptide boundaries with the wild-type peptide that established coverage while an acquired lysine produces a different peptide. **(B)** Peptide length, both classes. Medians 13 against 14 residues, so the difference in panel A is not a length artifact. **(C)** Cleavage after arginine fails about twice as often as cleavage after lysine. Peptides carrying a missed cleavage at the site, split by whether the first basic residue is K or R, PSM-weighted and by distinct sequence: K 15.0% / 23.6%; R 29.4% / 37.0%. **(D)** Acquired-lysine peptides are digested like any other. Missed cleavage across all identified peptides: 23.8% for peptides carrying an acquired lysine against 21.6% background. Panels C and D report missed / (missed + cleaved) among identified peptides, as the field reports it. A cleavage site that produced no observable peptide appears in neither term, so each is a lower bound on how often the protease fails rather than a digestion efficiency in the absolute sense. **(E)** Score margin. Hyperscore minus nextscore for the 89 acquired-lysine PSMs, which fall on 53 sites. 53 of the 89 PSMs clear margin ≥ 10, and those 53 PSMs fall on 30 of the 53 sites. **(F)** Spectrum quality. Hyperscore densities, variant against background: medians 33.6 against 31.2. **(G)** An annotated spectrum. GAPDH E106K in WM2664, scan 72116: 635 peaks, base peak 8378, 41 of 76 theoretical ions matched within 0.6 Da, and 18 of 19 backbone bonds observed, with y1–y3 directly evidencing the C-terminal lysine. The peptide is WGDAGAEYVVESTGVFTTMK, with the acquired lysine, written K[E106K], at its C terminus and the methionine at peptide position 19 oxidized (Met19 in the panel title). Teal marks b ions and rust y ions, in the sequence map and in the spectrum; each label carries the ion number as a subscript and the charge as a superscript, and gray peaks are unassigned TMT reporter and precursor-derived ions. Precursor 3+, 874.7772 observed against 874.7769 calculated; ion-trap CID; hyperscore 39.75, margin 16.0, expectation 1.6 × $10^{-3}$.

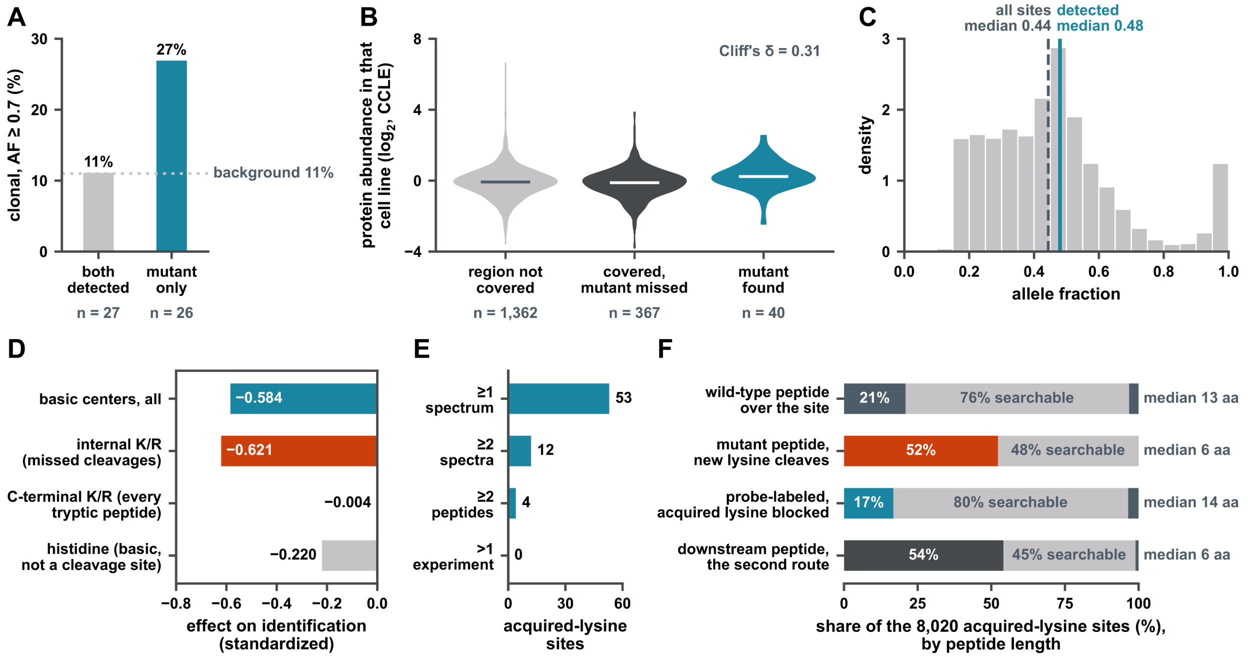


**Figure S3. Four explanations that are not the answer, Related to Figure 2. (A)** The detections are not carried by subclonal sites. Of the 53 sites where the mutant peptide was detected, the 27 whose wild-type region was also covered are 11% clonal (allele fraction ≥ 0.7) and the 26 seen only as the mutant are 27%, against an 11% background. **(B)** Protein abundance does not explain which regions are covered. Abundance from the CCLE quantification rather than a proxy; 1,769 of 8,470 observations have a value and the panel is drawn on those. Cliff's δ = 0.31 between found and never-covered, a real shift, far too small to account for roughly 93% of regions never being observed. **(C)** Allele fraction does not either. Detected sites have a median allele fraction of 0.48 against 0.44 across all acquired-lysine sites. **(D)** What predicts identification is incomplete digestion, not charge. Standardized effects: internal K/R (missed cleavages) −0.621; basic centers, all −0.584; histidine (basic, but never a cleavage site) −0.220; C-terminal K/R −0.004. Within peptides having no missed cleavage the apparent charge effect falls from −0.58 to −0.09. **(E)** The evidence behind the 53 sites, at full depth. 53 sites rest on at least one spectrum, 12 on two or more, 4 on two or more distinct peptides, and none on more than one experiment. Bars are drawn at the search's reported 1% false-discovery rate, which is the population this panel bounds; of these, 30 sites clear the score margin ≥ 10 of Figure 3B, with 24 on a single spectrum, six on two or more and two on two or more distinct peptides. **(F)** The mutant peptide is short, and the missed-cleavage allowance absorbs it. Peptide length for the four species that can carry an acquired-lysine site, over the 8,020 sites that map onto a reference sequence, at perfect digestion. Colored, shorter than seven residues; pale, seven to fifty and therefore searchable; dark, longer than fifty. The window is not a convention chosen here; both searches enforced it: digest_min_length = 7 and digest_max_length = 50 in the MSFragger parameters used for both architectures, confirmed in the output, where the shortest peptide identified is seven residues and the longest fifty; and the original CCLE search excluded "peptides below a length of 7" [S1]. Because a new lysine is a new cleavage site, the fully cleaved mutant peptide has a median length of six residues against thirteen for the wild-type peptide over the same site, and 52.3% of it falls below the floor. Acylating the acquired lysine removes that cleavage site and restores the geometry: median fourteen residues, 16.8% below the floor, the least truncated of the four. This panel is description, not explanation. Both searches allowed two missed cleavages, so an upstream missed cleavage lengthens a too-short mutant peptide back into the window, and 8,066 of 8,470 observations (95.2%) still carry a searchable species ending at the acquired lysine. The direct test does not resolve whether the geometry costs detections: only 27 of the 617 covered observations lack a fully cleaved species, and at that population's 4.4% recovery rate one event is expected, so the comparison, 1 of 27 against 26 of 590, has no power rather than a null result. What the panel does establish is that for half this class the only searchable species carries a missed cleavage, and panel D shows internal missed cleavages are the strongest negative predictor of identification measured here. Computed by step52_mutant_peptide_length.py; counts from the same per-observation table as panels A–E.

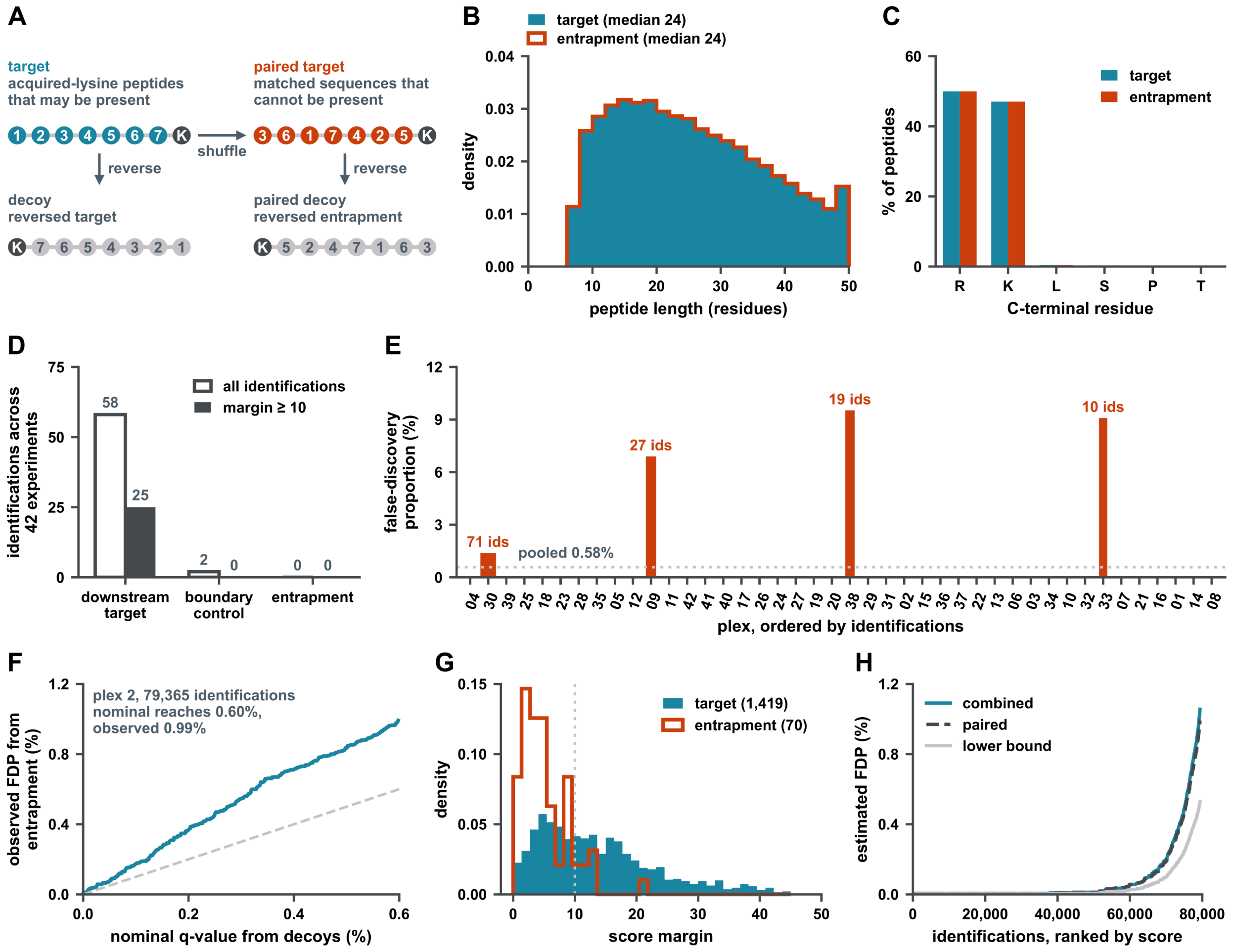


**Figure S4. The entrapment design in full, Related to Figure 3. (A)** The four-way design. 589,588 variant-pair rows read from 8 plexes; entrapment sets of 147,397 each. 100% of entrapment sequences share residues with their target. Each set holds 147,397 peptides. The entrapment peptide is its target's own residues permuted, with the C-terminal K or R kept, so it carries an identical mass and is still tryptic, and every entrapment identification is an observed false positive. In the schematic each circle is a residue: the target (teal); its paired target, which is the entrapment peptide of the other panels, made by shuffling the target's residues with the C-terminal residue kept (rust); and the decoy of each, the full reversal of its sequence (gray). **(B)** Length matching. Peptide-length densities for the two sets, identical by construction. Target is the filled block and entrapment the line over it. **(C)** What the shuffle preserves. C-terminal residue composition. Each entrapment peptide is its target's own residues permuted with the C-terminal K or R kept, so the pair shares an exact precursor mass and only fragment ions distinguish them. 100% of pairs share the same residues in a different order, so they have an identical precursor mass and only fragment ions discriminate. The R:K split survives because the C-terminal residue is held fixed. **(D)** Boundary control. Identifications across 42 experiments, all against those surviving margin ≥ 10: downstream target 58 → 25; boundary control (the same region shifted one residue, tryptic in neither protein) 2 → 0; entrapment for the downstream route 0 → 0. The boundary control is the same region shifted one residue, tryptic in neither the wild-type nor the mutant protein. **(E)** Per-plex FDP. 42 plexes ordered by identifications, 0.00% to 9.52%, pooled 0.58%. 38 of the 42 experiments contributed no entrapment identification at all. **(F)** Nominal against observed FDR. On plex 2 (79,365 identifications) the nominal q-value from decoys reaches 0.60% where the observed proportion from entrapment reaches 0.99%; decoys underestimate by 1.7-fold. The diagonal is where the two would agree. **(G)** All four sets. Score-margin densities. Entrapment peptides are identified, and score like a population that cannot be present should. **(H)** A published estimator, on the whole search. The Wen et al. estimator over the entire plex-2 search (79,365 identifications, not the variant subset), so it is not comparable with the 0.58% above. This runs over the whole plex-2 search rather than the variant subset, so it is not comparable with the 0.58% in panel E.

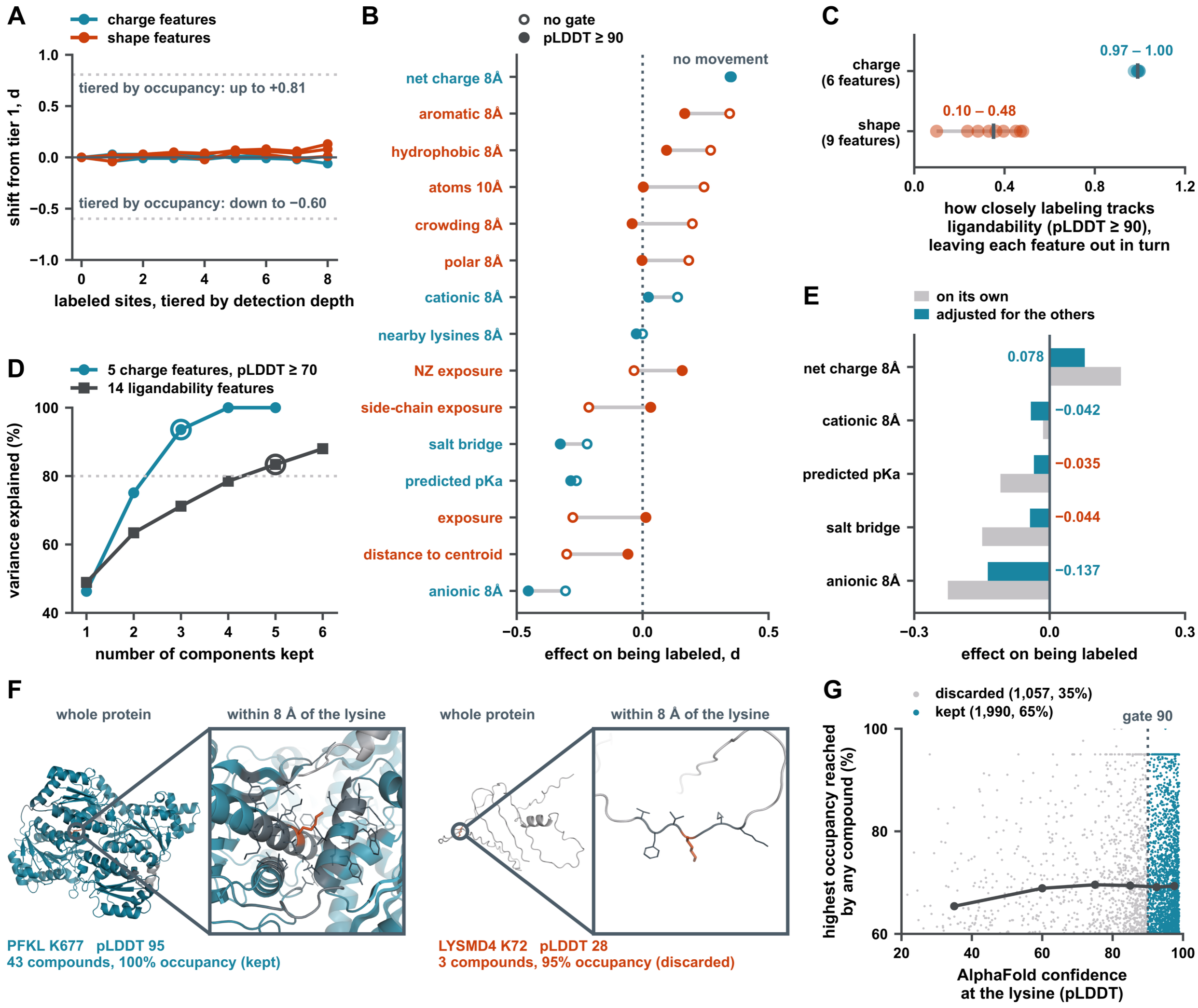


**Figure S5. What the confidence gate does, Related to Figure 4. (A)** The tiering null. Five series. The steepest slope per tier is 0.013, against 0.083 for ligandability, 6.3 times smaller, and the same feature appears on both sides. Labeled sites are tiered by detection depth rather than by occupancy; teal lines are charge features and rust lines shape features. The two dotted lines mark how far the same shifts reach when the tiering is by occupancy, up to +0.81 and down to −0.60. **(B)** Every feature, both gates. 15 features. Each feature's effect on being labeled with no gate (open circle) and at pLDDT ≥ 90 (filled circle), joined by a gray bar; feature names are teal for charge features and rust for shape features, as in panel A. "No movement" marks the one feature whose two values differ by less than 0.02. **(C)** Leave-one-out robustness. 15 features, the same set as Figure 4, each dropped in turn. Each point is how closely labeling tracks ligandability at pLDDT ≥ 90 with one feature of its class omitted, the printed range spans those values, and the vertical mark is the value with every feature of the class in. **(D)** Dimensionality. Principal components explain 46.3, 28.8, 18.5, 6.4 and 0.0% of variance for reactivity and 48.9, 14.5, 7.8, 7.3, 4.9 and 4.6% for ligandability; reactivity is close to three-dimensional, ligandability is not. 5 charge features reduce to 3 axes and 14 ligandability features to 5. Each set is redundant and neither collapses to one axis, but they are different sets on different populations, so this is not 3 against 5. The dotted line marks 80% of variance, and the ringed point on each curve is the first to exceed it. **(E)** Mutual adjustment. Each charge feature on its own against the same feature adjusted for the others. Salt bridge was anionic-neighbor density in another form, and does not survive the adjustment. **(F)** What the gate removes, shown rather than asserted. Two real engaged sites, each rendered whole-protein and as the lysine with everything within 8 Å, from the same AlphaFold models the features are measured on. PFKL K677, pLDDT 95, 43 compounds at 100% occupancy, kept. LYSMD4 K72 (UniProt Q6MZR2, which the deposited site table carries under its TrEMBL clone name DKFZp779J2370), pLDDT 28, 3 compounds at 95% occupancy, discarded. The same 8 Å sphere: in one of them there is no structure to measure. **(G)** The gate is not a strength filter, the control that excludes the objection that gating simply keeps the strongest sites. Median occupancy is flat across the whole confidence range, so the gate removes sites whose features cannot be measured rather than sites that are weakly engaged. The counts in the key are the sites it discards and the sites it keeps, and the dark line is the median occupancy in six confidence bands.

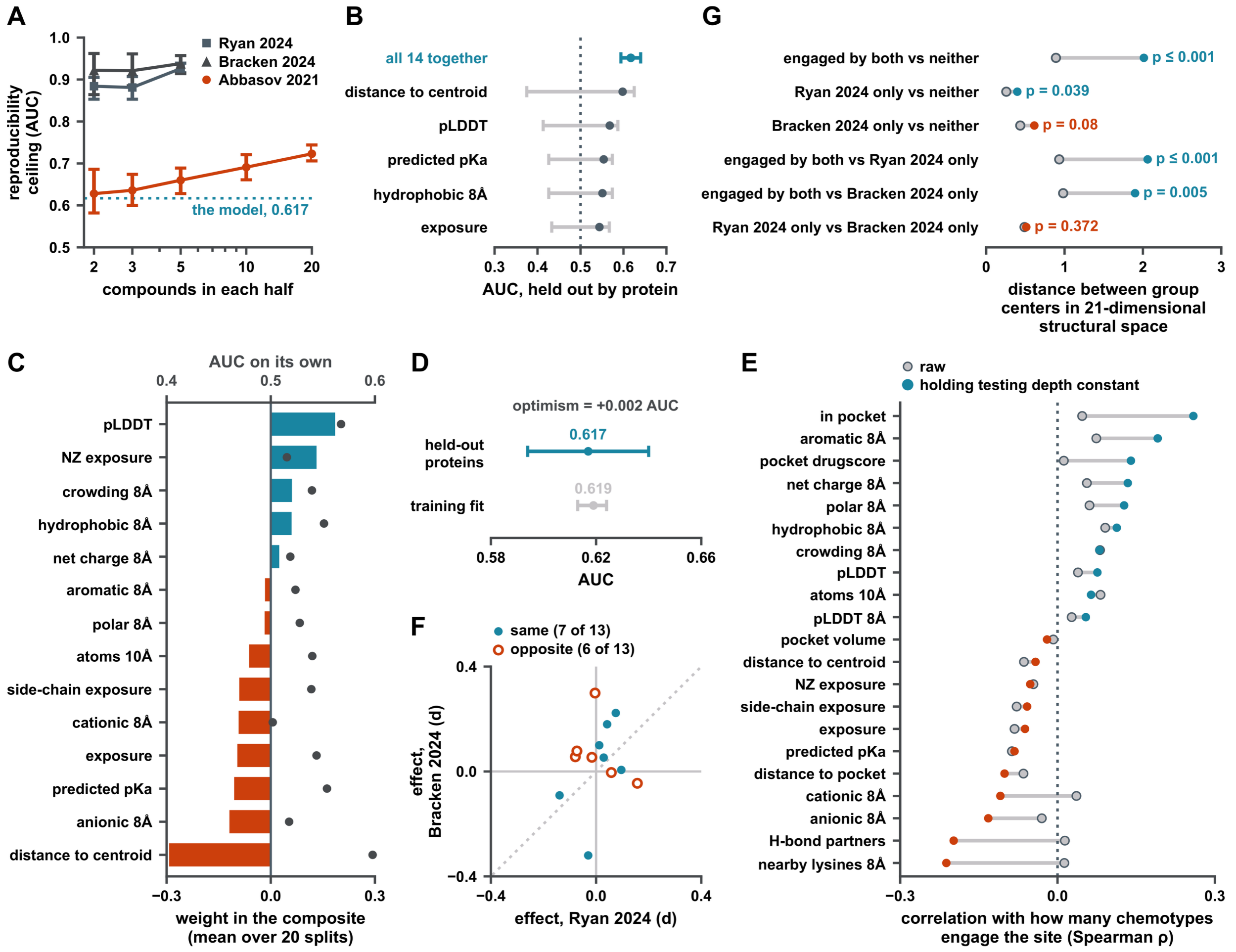


**Figure S6. The ceiling, and the limit, Related to Figure 5. (A)** The ceiling per dataset, against how deeply each screen is split. The key names each screen by the first author and year of its report: Ryan 2024 is the CyMPSA screen [S2], Bracken 2024 the meroterpenoid screen [S3] and Abbasov 2021 the lysine-targeting compound screen [S4]; each one's median compounds per site and the share of its sites engaged at the deepest split are in Table S2. Intervals are the 2.5th to 97.5th percentile across 50 random half-splits, the same construction as Figure 5C. The two shallow screens reach 0.881 to 0.938, but at engaged rates of 0.6 to 2.3%, where a few dozen positives make an AUC unstable. The lysine-targeting compound screen is the only screen deep enough to measure a ceiling at a realistic engaged rate, and its ceiling is still rising at 20 compounds per half, so 0.723 is a lower bound. The deeper the split, the less noisy the label. **(B)** Single features against the composite. Five of the 14 features alone, against the composite, held out by protein; every feature's own AUC is in panel C. Intervals are the 2.5th to 97.5th percentile across the 20 protein-held-out splits, as in panel D. The composite reaches 0.617 against 0.598 for the best single feature, distance to the protein centroid, a gain of +0.019. The 14 features are not independent. **(C)** Every coefficient. The 14 composite coefficients, averaged over the 20 protein-held-out splits, with each feature's own AUC on the twin axis. An AUC below 0.5 is an inverted feature rather than a weak one, so each is drawn at its flipped value: distance to the centroid reads 0.402 and is drawn at 0.598. **(D)** Train against held-out. Train 0.619, held-out 0.617, optimism +0.002. Intervals are the 2.5th to 97.5th percentile across the 20 protein-held-out splits, the same construction as Figure 5C and 5D. The model is not overfit. It is fitting all there is to fit, and that is the point. **(E)** The breadth gradient, adjusted for testing depth: 21 breadth features, the control for the objection that broadly engaged sites are simply the sites tested most often. Holding depth constant strengthens most of the gradient and flips 3 features outright. Gray markers are the raw correlations and colored markers the depth-adjusted ones, teal where positive and rust where negative. **(F)** Two chemotypes agree. 13 chemotype features, each a point, with its effect in the CyMPSA screen (Ryan 2024) on the x axis and in the meroterpenoid screen (Bracken 2024) on the y axis; filled teal points have the same sign in both screens and open rust points opposite signs. The points are unlabeled because thirteen names do not fit the panel; the three that disagree most between the chemotypes are net charge 8 Å, anionic 8 Å and hydrophobic 8 Å, and every per-feature value is in panel C and Table S2. None of the 13 differs significantly between the two chemotypes, and the same sites are engaged by both (26 of 8,211 shared, 5.5 expected, OR 6.05), so one general composite is defensible. **(G)** Across datasets, a limit, not a result. This is the only cross-dataset evidence in the paper and the weakest evidence in it, and Extended Results depends on it. The group carrying the cross-dataset claim, the sites engaged by both screens, is 25 sites in 21 dimensions, against 303 engaged only in the CyMPSA screen (Ryan 2024), 107 engaged only in the meroterpenoid screen (Bracken 2024) and 7,183 engaged in neither (these 7,618 are the sites among the 8,211 of panel F that carry all 21 features, and they keep 25 of its 26 shared sites), and 2 of 6 pairs fall short of their null over 2,000 label shuffles. Extended Results repeats this test along the breadth gradient within one dataset, and this panel is what happens when it is asked across two. Gray is the null, teal clears it, rust does not.

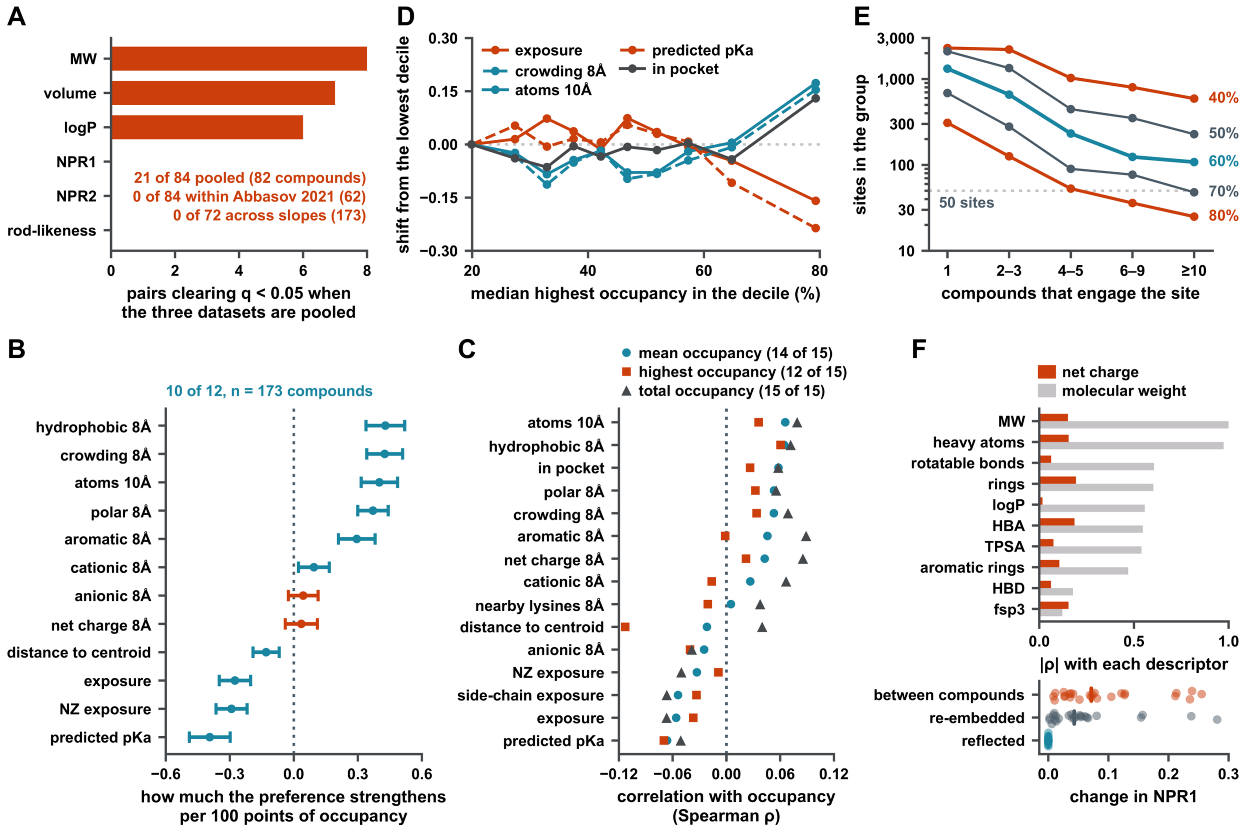


**Figure S7. The confound, and what no descriptor can see, Related to Figure 5. (A)** The provenance confound named. Pooled across the three screens, 21 of 84 descriptor × site-feature pairs clear q < 0.05 over 82 compounds, and every one of them falls on molecular weight (8), volume (7) or logP (6). The three pure-shape descriptors, NPR1, NPR2 and rod-likeness, produce none, so what survives pooling is the size and lipophilicity axis rather than shape. Within the lysine-targeting compound screen alone, labeled Abbasov 2021 in the panel, the count is 0 of 84 (62 compounds), and across the within-dataset slopes 0 of 72 (173 compounds). This is the same analysis as Figure 5E, at the same inclusion rule of at least fifteen engaged sites per compound; the panel names which descriptors carry that figure's pooled column, and Extended Results shows those are the descriptors that separate the three screens. **(B)** Per-compound slopes across occupancy. For each of 12 site features, the mean over 173 compounds of the per-compound slope of the feature's standardized shift against occupancy, per 100 points of occupancy; the 173 are the compounds with slopes across at least five occupancy tiers. Intervals are 95% confidence intervals (mean ± 1.96 standard errors). Teal, the mean slope differs from zero (one-sample t test, Benjamini–Hochberg q < 0.05), which is true of 10 of the 12 features; rust, it does not. **(C)** Three threshold-free continuous measures. Spearman ρ of 15 site features with mean occupancy (14 features significant at q < 0.05), highest occupancy (12) and total occupancy divided by the number of compounds tested (15), across 11,592 lysines with a pLDDT of at least 70 that were tested against 20 or more compounds, so the Figure 5 result does not depend on a 60% cut existing at all. **(D)** The decile profile. The same lysines in deciles of highest occupancy. Each line is the standardized shift of one feature from the lowest decile, plotted at the decile's median highest occupancy. Solid and dashed lines only distinguish two features drawn in the same color. **(E)** Where group sizes collapse. Sites are grouped by how many compounds engage them, at five occupancy cuts, and the panel shows the number of sites in each group. Teal is the 60% cut used throughout, rust the two cuts that are not strictly monotonic in Extended Results (40% and 80%), dark gray the other two. The dotted line marks 50 sites. At the 80% cut the groups are the smallest of any cut, down to 25 sites, and one step of the gradient reverses; at the 40% cut the groups are the largest of any cut and the first two are level to three decimals. Small groups therefore account for one of the two exceptions and not for both. **(F)** What no tested descriptor can see. Bars show each tested descriptor's |ρ| with net charge (rust) and with molecular weight (light gray). The strip plot shows, for 25 chiral compounds, the change in NPR1 on reflecting a conformer (teal), on re-embedding the enantiomer (dark gray) and each compound's distance from the mean NPR1 (rust); vertical bars are medians. Charge: maximum |ρ| across tested descriptors is 0.193, against a median |ρ| of 0.552 for molecular weight. Chirality: NPR1 and NPR2 are ratios of principal moments of inertia and achiral by construction, so reflecting a conformer changes NPR1 by 0.000 while re-embedding an enantiomer changes it by up to 0.281, against a between-compound standard deviation of 0.123, so that figure is conformer sampling rather than chirality. This panel bounds Figure 5's claim rather than supporting it.

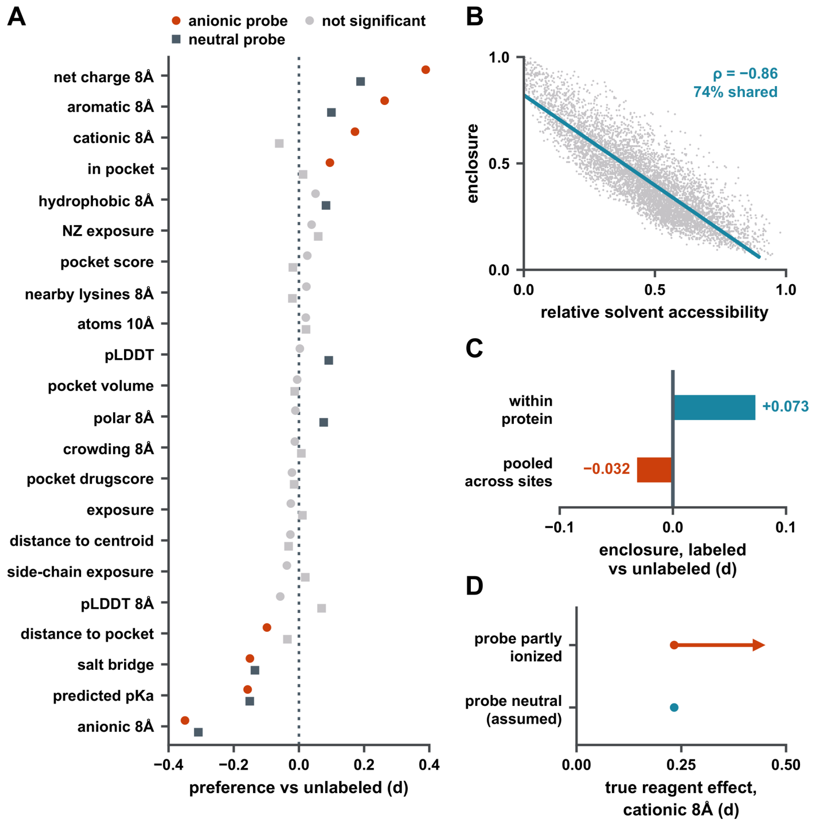


**Figure S8. Each probe alone, and the pooling reversal, Related to Figure 6. (A)** Each probe against unlabeled lysines. 22 features per probe, ordered by the anionic probe's effect; circles are the anionic probe and squares the neutral probe. The effect d is the within-protein standardized difference between labeled and unlabeled lysines. Colored marks are significant at $q < 0.05$ and light gray marks are not; 8 are significant for the anionic probe and 8 for the neutral one, so the two-of-twenty-two difference in Figure 6B is a statement about the contrast rather than about power. **(B)** Enclosure is not a restatement of solvent accessibility. Enclosure against relative solvent accessibility. The points are a random subsample of 6,000 of the 167,057 sites and the line is the linear fit; the fit and the correlation are computed on all 167,057 sites. Four of the 6,000 points have a relative solvent accessibility above 1 and lie beyond the axis. Spearman $\rho = -0.862$, so the two share about 74% of their variance and roughly a quarter is enclosure's own. **(C)** Pooling reverses the sign of the enclosure effect. Pooled across sites, labeled lysines look less enclosed ($d = -0.032$); within protein they are more ($d = +0.073$). The within-protein value is the one Figure 6E reports. **(D)** What the neutral-probe assumption bounds. If the neutral probe is in fact partly ionized, the true reagent effect on cationic environment is larger, not smaller: +0.233 under the neutral assumption, running to +0.46 if the probe is substantially ionized. The assumption makes the reported effect a lower bound. This is a sensitivity range across assumed ionization states, not a confidence interval: it carries no sampling statement, and the two endpoints are the neutral and substantially-ionized limits rather than percentiles.

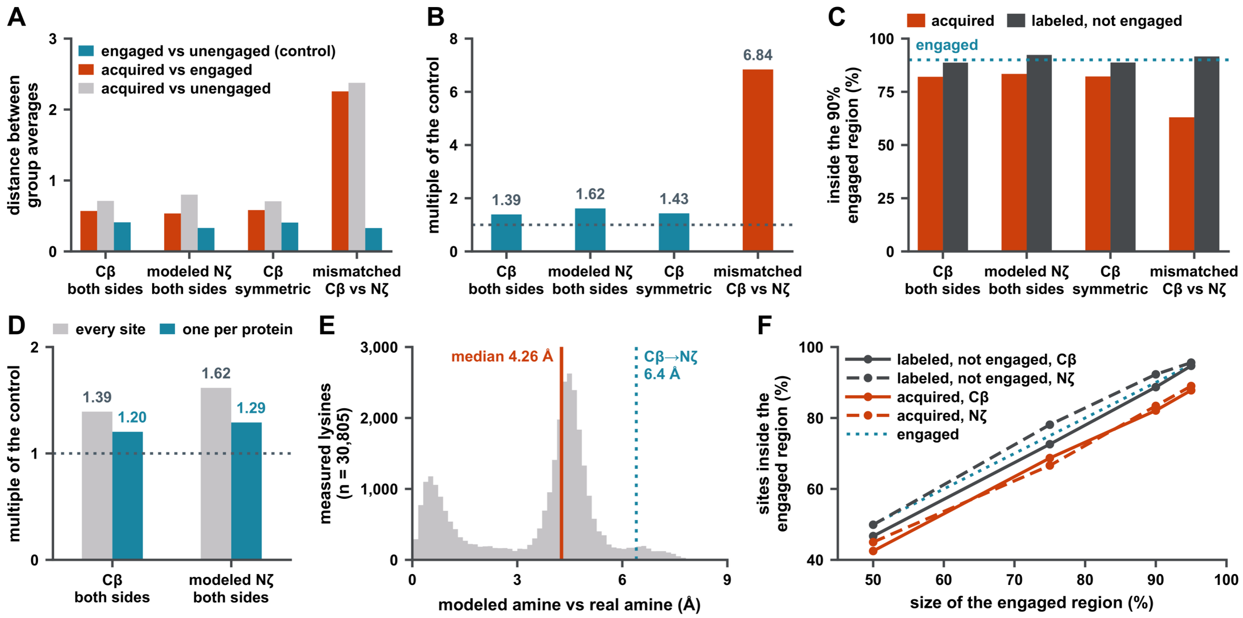


**Figure S9. The anchor tested rather than defended, Related to Figure 7. (A)** Distances, four ways, and (B) each arm against its own control. Three legitimate anchorings (both sides at Cβ, both sides modeled at Nζ, and both at Cβ under a symmetric convention) give acquired-versus-engaged distances of 0.571, 0.535 and 0.583 against controls of 0.410, 0.331 and 0.407. The mismatched anchoring, union positions at Cβ against measured lysines at Nζ, gives 2.256 against 0.330, a 6.84-fold ratio. The anchor does not change the answer; the mismatch does. The mismatched arm is retained as a negative control because it is the analysis this project originally ran. **(C)** Overlap, four ways. The dotted line marks 90%, the share of engaged lysines that lies inside the region by construction. **(D)** One site per protein. The correction for proteins contributing many lysines; the ratio shrinks but the direction holds. **(E)** The built side chain, checked against real lysines. All-trans rotamers were constructed at every position and compared with the amine positions in the measured set: median error 4.26 Å across 30,805 measured lysines, against a Cβ-to-Nζ separation of 6.4 Å. All-trans is the weakest possible construction, so this is an upper bound on the construction error rather than an estimate of it, and the reference amine positions are themselves predicted, since pLDDT is a backbone measure that does not certify side-chain placement. **(F)** Overlap at every cut. Four quantile cuts (50, 75, 90, 95), both legitimate arms, both comparison populations. Acquired lysines are at 42 / 69 / 82 / 88% inside the 50 / 75 / 90 / 95% engaged region with both sides at Cβ (solid lines) and 45 / 67 / 83 / 89% with both sides modeled at Nζ (dashed lines); labeled-but-unengaged lysines are at 47 / 73 / 89 / 95% and 50 / 78 / 92 / 96%, in the same order. The dotted line is the engaged lysines themselves, which by construction sit at each cut. The reading does not depend on where the cut is drawn.

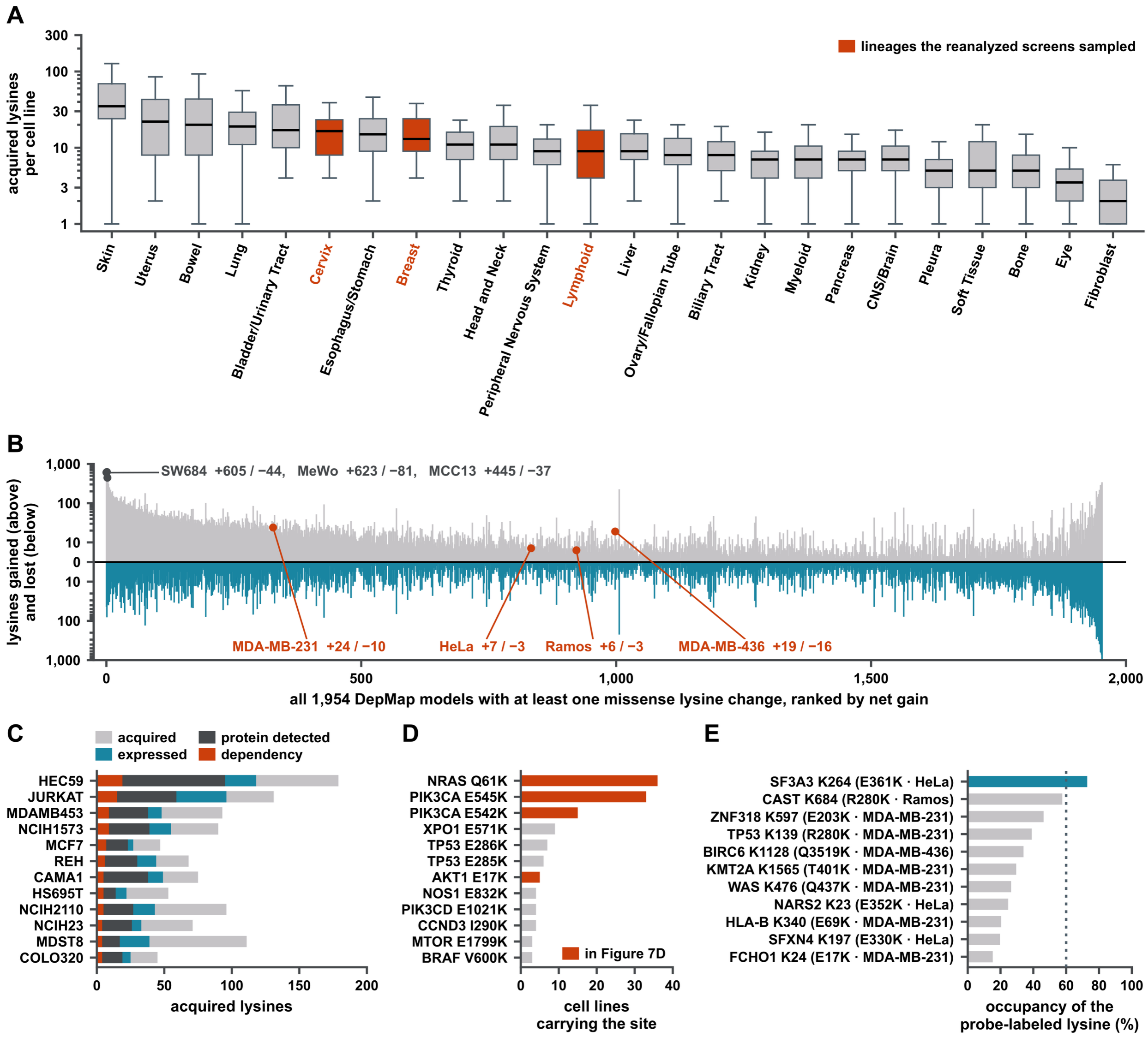


**Figure S10. The acquired-lysine class across cell lines, and what a re-search of the three screens could find, Related to Figure 7.** Counts are unique (model, gene, protein change) throughout. DepMap 26Q1 carries duplicate rows: 55,116 acquired-lysine rows collapse to 39,390 unique across 1,954 models, and 936 models are affected. **(A)** Burden by lineage recovers the mutational processes of Figure 1E in a second dataset. Acquired lysines per cell line, log scale, boxes at the quartiles with medians marked, whiskers to the most extreme value within 1.5 interquartile ranges of the box and more extreme values not drawn, lineages with at least 15 models, ordered by median. Skin leads at 35, then uterus 22, bowel 20 and lung 19; eye sits second from last at 4. At subtype resolution the same contrast is sharper still: cutaneous melanoma 37 (n = 130) against ocular melanoma 3 (n = 15), which is the cutaneous-versus-uveal comparison of Figure 1F in cell lines rather than tumors. Metastatic models carry more than primary ones (median 13 against 9, n = 576 and 1,089, $p = 7.7 \times 10^{-8}$, Cliff's δ = −0.160). Rust, the three lineages the reanalyzed screens sampled: cervix (HeLa), breast (MDA-MB-231, MDA-MB-436), lymphoid (Ramos). **(B)** Every cell line in DepMap. All 1,954 models carrying at least one missense lysine change, ranked by net gain; gray above the line, lysines gained, teal below, lysines lost. Symlog scale, linear within ± 10. Across all models 39,390 lysines are gained and 25,531 lost, a ratio of 1.54, and 494 models (25.3%) are net-negative. The ratio is strongly lineage-dependent: 6.4 gained per lost in skin, 2.8 in bladder and soft tissue, 1.3 in esophagus and stomach, the C>T asymmetry of Figure S1F seen one cell line at a time, since lysine is gained 85.5% by C>T and never lost by it. Labeled: the three highest net gainers and the four cell lines the reanalyzed screens were run in (rust). The extreme loss tail is a hypermutator losing 999 lysines. **(C)** Four nested gates, and the lines that carry the class are the least characterized. Of the 1,954 models, 1,630 have expression, 1,204 CRISPR and 372 CCLE proteomic data; only 308 have all three, and none of the ten highest net-gain lines does; MeWo has no CRISPR entry, SW684 none of the three. Bars are the twelve of those 308 with the most sites clearing every gate. Gray, unique acquired lysines; teal, gene at $\log_2(\text{TPM}+1) \geq 1$ in that line; charcoal, protein also quantified in that line's CCLE proteome; rust, gene also a CRISPR dependency at gene effect ≤ −0.5. Aggregate over the 28 models examined: 2,236 → 1,035 expressed (46%) → 686 also detected (31%) → 132 also a dependency (6%). **(D)** The recurrent driver-flagged acquired lysines are the sites Figure 7D already measured. 248 of 39,390 unique acquired lysines carry a hotspot or HessDriver flag (0.63%), in 210 of 1,954 models. Ranked by the number of cell lines carrying each: NRAS Q61K 36, PIK3CA E545K

33, PIK3CA E542K 15, XPO1 E571K 9, TP53 E286K 7, TP53 E285K 6, AKT1 E17K 5, the sites in five or more models. Rust marks the four with a structure in Figure 7D: three of them are the three most recurrent, and the fourth, AKT1 E17K, ranks seventh. BRAF V600K, three models, is an acquired lysine on the most-drugged oncogene in melanoma. Driver-flagged sites clearing all three gates of panel C in the models drawn: MTOR E1799K (HEC59), PIK3CA E545K (MCF7), BRAF V600K (MDST8 and IGR1), PIK3CD E1021K (REH). **(E)** Eleven acquired lysines in the four screen lines sit on a protein the matching screen probe-labels, and the measured lysine is never the acquired one. A site is reachable when the screen run in that line labels at least one lysine on that protein, the precondition for measuring occupancy. Each bar is the highest occupancy at the best-engaged probe-labeled lysine on that protein in that screen, named on the axis with the acquired mutation in parentheses; the acquired lysine was outside the search space of all three screens and has no occupancy value. Labeled and acquired positions are a median of 271 residues apart (7 for FCHO1 to 2,391 for BIRC6), so a bar reports that the protein is reachable by a probe, not that the acquired site is. Teal bar, the one protein whose labeled lysine is engaged at ≥ 60% (SF3A3 K264, 72.8%). Cell lines: SF3A3, NARS2, SFXN4 HeLa; CAST Ramos; BIRC6 MDA-MB-436; the rest MDA-MB-231. TP53 R280K is one of the six sites measured in Figure 7D. Dotted line, the 60% engagement convention. Computed by figS10_v1.py from DepMap 26Q1 somatic mutations, expression, CRISPR gene effect and model annotation, the CCLE proteomes, and the per-site occupancy table behind Figures 4 to 7; every number above is printed by that script. Sites are matched to screens by cell line: HeLa to CyMPSA, MDA-MB-231 to the meroterpenoid and lysine-targeting compound screens, MDA-MB-436 to the meroterpenoid screen, Ramos to the lysine-targeting compound screen.

**Table S1. The recurrent acquired lysines: how many patients carry them, and what they are reported to do, Related to Figure 7.**

| Site | TCGA patients | Where this paper measures it | Reported functional consequence | Evidence | Primary source |
|---|---|---|---|---|---|
| NRAS Q61K | 68 (0.67%) | Figure 7D structure; 36 models (Figure S10D) | Slows GTP hydrolysis rather than abolishing it, and the effect belongs to the loss of Gln61 rather than to the lysine that replaces it. Every codon-61 substitution tested in HRAS, whose catalytic domain is identical to that of NRAS through residue 86, hydrolyzed GTP eight- to tenfold more slowly than wild type, while transforming potency across the same set varied by more than a thousandfold, so the rate loss alone does not account for transformation [S5]; the tenfold figure and the decoupling were reproduced independently, with crystal structures attributing the rate loss to displacement of the water molecule bridging Tyr32 to the γ-phosphate [S6]. GAP-stimulated hydrolysis, by contrast, is not uniformly impaired at codon 61: NMR profiling found Q61L unexpectedly susceptible to it, in direct contrast with G12V [S7]. For NRAS itself, a crystal structure of the GTP-bound Q61K mutant resolves a ligand-induced pocket beside switch II [S8]. In vivo, NRasQ61K on an INK4a-deficient background produces metastasizing melanoma in mice [S9], and the allele drives aberrant melanocyte signaling, survival and invasiveness through Rac1 [S10]. | Established driver | [S5–S10] |
| PIK3CA | 263 | Figure 7D | Helical-domain hotspot. | Established | [S11,S1 |

| | | | | | |
|---|---|---|---|---|---|
| E545K | (2.57%) | structure; 33 models (Figure S10D) | Oncogenic in vivo in the chick chorioallantoic membrane, with increased angiogenesis and Akt activation [S11]; separately, releases p85β from p110α to translocate to the nucleus, where it stabilizes EZH1/EZH2 [S12]. | driver | 2] |
| PIK3CA E542K | 169 (1.65%) | Figure 7D structure; 15 models (Figure S10D) | As E545K: oncogenic in vivo [S11] and acting through the same p85β nuclear-translocation route rather than canonical PI3K–AKT alone [S12]. | Established driver | [S11,S12] |
| XPO1 E571K | 11 (0.11%) | 9 models (Figure S10D); not among the sites clearing all three gates in the models drawn (Figure S10C) | Alters cargo recognition at the NES-binding groove. Across 1,286 CLL patients, XPO1 E571 mutations associate with high-risk features; in mice, E571K with TCL1 accelerates leukemogenesis. Selinexor and eltanexor binding is unaffected [S13]. | Established driver | [S13] |
| AKT1 E17K | 54 (0.53%) | Figure 7D structure; 5 models (Figure S10D); the site of Figure 7E | Pleckstrin-homology-domain mutation driving constitutive membrane recruitment and PI3K–AKT–mTOR activation [S14]; the one acquired lysine engaged selectively by a covalent campaign [S15]. | Established driver | [S14,S15] |
| TP53 R280K | 16 (0.16%) | Figure 7D structure | Reported as a gain-of-function allele promoting NOX4-dependent reactive-oxygen production and focal-adhesion-kinase activation, and so migration and invasion [S16], rather than acting only through the loss of DNA binding that its crystal structure explains [S17]. | Functionally validated | [S16,S17] |
| PTPN11 E76K | 2 (0.02%) | Figure 7D structure | Charge reversal at the N-SH2–PTP autoinhibitory interface destabilizes the closed conformation of SHP2. Gain-of-function Ptpn11 alleles including E76K cause cytokine hypersensitivity in | Established driver | [S18] |

| | | | | | |
|---|---|---|---|---|---|
| | | | myeloid progenitors and myeloproliferative disease in mice [S18]. | | |
| BRAF V600K | 0, see note | 3 models (Figure S10D); clears all three gates in two lines (Figure S10C) | Constitutive BRAF–MEK–ERK activation. Present in 10–30% of BRAF-mutant melanoma [S19]; clinically distinct from V600E, with shorter progression-free survival on BRAF-directed therapy [S20] and lower ERK but higher PI3K–AKT pathway expression [S21]. | Established driver | [S19–S21] |
| TP53 E285K | 24 (0.23%) | 6 models (Figure S10D) | Disrupts a salt-bridge network stabilizing the DNA-binding domain, abolishing transcriptional activity and mislocalizing the protein to the cytosol, with cisplatin resistance reversible by ferroptosis induction [S22]; reported as a late secondary event in lobular breast-cancer evolution, where the mutant is temperature-sensitive and its reconstitution restores CDKN1A, MDM2 and PHLDA3 [S23]. | Functionally validated | [S22,S23] |
| TP53 E286K | 16 (0.16%) | 7 models (Figure S10D) | Transactivation-defective at every response element tested. In the arrayed yeast assay that scored the 2,314 p53 missense substitutions reachable by a single point mutation against eight response elements, E286K retained 18.3% of wild-type transactivation at the p21WAF1 promoter and 21.8% at p53R2, and between 6.0% and 25.9% across the eight, so the substitution is strongly hypomorphic rather than silent [S24]. | Loss of function | [S24] |
| MTOR E1799K | 11 (0.11%) | Clears all three gates (Figure S10C) | Hyperactivating. E1799K is one of 33 pathway-activating MTOR mutations curated from tumor sequencing, and one of ten expressed and assayed directly; all ten raised | Gain of function | [S25] |

| | | | | | |
|---|---|---|---|---|---|
| | | | phosphorylation of the mTORC1 and mTORC2 substrates S6K1, 4EBP1 or AKT1 [S25]. Two cancer lines carry it natively, HEC59 (endometrium) and SNU349 (kidney), with rapamycin $IC_{50}$ values of 0.49 and 0.19 nmol/L against 500 nmol/L for a line carrying a non-activating MTOR mutation, and rapamycin halted HEC59 xenograft growth outright [S25]. HEC59 is the model in which this site clears all three gates here. | | |
| XRCC5 E621K | 7 (0.07%) | Clears all three gates (Figure S10C) | No mutation-specific functional study identified. XRCC5 (Ku80) partners Ku70, whose K351 is one of the liganded lysines in the lysine-targeting compound screen [S4]. | Not assessed | None |
| PIK3CD E1021K | 1 (0.01%) | Clears all three gates (Figure S10C) | The one acquired lysine here with a proven gain of function, established in the germline rather than in cancer: E1021K is the canonical allele of activated PI3Kδ syndrome, and the substitution increases p110δ membrane association and kinase activity [S26], with germline gain of function at this subunit driving T-cell senescence and immunodeficiency [S27]. Reported somatically in Burkitt lymphoma [S28], but not characterized as a somatic driver. PIK3CD already carries approved drugs directed elsewhere on the protein. | Gain of function shown in the germline | [S26–S28] |

Patient counts are distinct TCGA participants carrying the substitution in the MC3 call set, against the 10,224 participants with at least one MC3 call; percentages are of that cohort. MC3 is already one of the three variant sources behind Figure 1, so this adds a patient denominator rather than a new dataset. No PASS-only filter is applied, for the reason given in the Methods: it would delete the ovarian and myeloid cohorts entirely; the PASS-only counts are in tcga_site_frequency.json and are within a few patients of these throughout. Column 3 is what this paper measures; columns 4 to 6 are what others report, and nothing in them is a result of this work. The table exists so that the sites named in the Discussion carry their independent evidence with them, and so that the sites with none are visible rather than absent.

**12 of the 13 rows carry a verified primary citation; the thirteenth, XRCC5 E621K, has none.** Every reference that does appear was verified against PubMed, with its PMID recorded in the bibliography file.

**BRAF V600K returns zero patients, and the zero is a property of the call set rather than of the disease.** MC3 contains no V600K call at all; at codon 600 it carries 588 V600E, 40 V600M, five V600V, four V600G and two indels. V600K is the one site in this table that cannot be reached by a single nucleotide substitution: valine’s codons are two changes from lysine’s, while the other twelve sites are one. MC3 calls single-nucleotide variants and does not merge adjacent ones into a dinucleotide event, so a substitution requiring two adjacent changes is invisible to it by construction, which is why DepMap reports V600K in three cell lines and this cohort reports it in none. The same reasoning applies to every acquired lysine created from A, C, D, F, G, H, L, P, S, V, W or Y, all of which need at least two nucleotide changes; only E, I, M, N, Q, R and T reach lysine in one. Patient counts in this column are therefore a floor for the class as a whole, not only for this row.

**Table S2. Panel-level values for the main figures, Related to Figures 1–7.**

*Every value plotted in a main figure panel, moved here so that each main legend states what is shown, the test, the unit, the replicate structure and the definition of every measure and error bar without also restating the panel. Nothing that bounds a claim was moved.*

| Figure | Panel | Values |
|---|---|---|
| Figure 1 | B | E→K ranks first of 363 observed substitution pairs at 181,252 sites, 4.12% of all missense; context-corrected enrichment 1.12-fold, against 3.59, 3.38 and 3.16 for R→H, R→C and R→Q. |
| Figure 1 | G | The gnomAD v2.1.1 non-cancer exome subset gives 508.0 per individual, 3.41%. Across all twenty residues lysine ranks third at 2.05 and cysteine fourth at 1.76, in a distribution from 0.43 to 2.63, median 1.07. |
| Figure 2 | A | Residues 280–299 are a single tryptic peptide of 20 residues in the wild-type protein. For an E→K site the mutant is heavier by 303.26 Da once the new lysine carries TMTpro, with different hydrophobicity and charge state. **This site is an instance of the asymmetry the figure is about, not an illustration of it:** across the 42 plexes the 20-residue wild-type peptide was never observed and the 8-residue mutant peptide was, which places LDHB E287K among the 26 of 7,853 uncovered sites in panel C. |
| Figure 2 | B | A conventional 20,431-protein search covers 640 (7.56%); the mutation-aware peptide-level search over 10.17 million entries covers 617 (7.28%). 7,853 sites, 92.7%, are never covered in any of the 42 plexes. |
| Figure 2 | C | Thirty of those 53 also clear the score margin of Figure 3B: 16 of 617 covered and 14 of 7,853 uncovered, 0.35% of the class on 29 distinct genomic sites. |
| Figure 2 | F | Of the 53 mutant-peptide sites, 41 rest on a single spectrum and 12 on two or more; of the 30 past the margin, 24 and six (Figure S3E). |
| Figure 3 | B | The false-discovery proportion is 4.70% at margin 0, 1,419 variant peptides against 70 entrapment, and falls to 3.91% at a margin of 2, 2.36% at 5 and 0.83% at 10, where 832 variant peptides survive against seven entrapment. |
| Figure 3 | C | Teal is the variant identifications across all 42 plexes, 5 to 79 per plex. Rust is the entrapment, six identifications falling in four plexes, 30, 09, 38 and 33. |
| Figure 3 | D | Peak count 0.91-fold, signal-to-noise 1.19-fold, from medians of 631 against 695 peaks and 0.221 against 0.185. |
| Figure 4 | B, C | Average distance between the pair: ungated charge 0.074, shape 0.075; gated charge 0.068, shape 0.126. |
| Figure 4 | D | The strip beneath each rung is what that rung keeps, drawn to scale against the ungated 245,684 lysines, with the labeled share as the dark segment: none 245,684 of which 30,805 labeled (246k on the figure, 100% kept, 13% labeled), ≥ 50 189,913 of which 28,316 (190k, 77%, 15%), ≥ 70 167,057 of |

| | | |
|---|---|---|
| | | which 25,979 (167k, 68%, 16%), ≥ 90 97,722 of which 17,409 (98k, 40%, 18%). |
| Figure 4 | E | The upper axis is that agreement as the gate tightens: charge stays at 0.98 to 0.99 throughout while shape falls 0.97, 0.90, 0.77, 0.35 at gates of none, 50, 70 and 90. The lower is the null itself, all 5,005 ways of splitting these fifteen features six against nine, with both observed values marked: +0.009 ungated and +0.639 gated. |
| Figure 5 | A | Of the 1,642,330 site × compound measurements in the three screens, 8,019 (0.49%) reach 60%. Counts per five-point occupancy bin fall steadily through the cut, 5,411 (50–55%), 3,582 (55–60%), 2,649 (60–65%) and 1,726 (65–70%). |
| Figure 5 | C | AUC 0.617 [0.594, 0.640] against a ceiling of 0.723 [0.706, 0.744] derived from agreement between independent halves of the same screen, a gap of 0.106; train AUC 0.619. 30,805 lysines on 6,201 proteins, 3,047 of them (9.9%) engaged by at least one compound. |
| Figure 5 | D | At 5% false positives it recovers 11% of engaged sites, and at 10% it recovers 19%. |
| Figure 5 | E | Pooled across the three screens (82 compounds), 21 of 84 pairs clear it against 4.2 expected; within the lysine-targeting compound screen alone (62 compounds), 0 of 84; in the within-dataset slope analysis (173 compounds), 0 of 72 against 3.6 expected. |
| Figure 6 | C, D | Cationic residues within 8 Å, d = +0.233, and net charge within 8 Å, d = +0.180, are the two features that separate the probes; the next largest, aromatic contacts at d = 0.118, does not. |
| Figure 6 | E | Enclosure d = +0.073 ($q = 1.1 \times 10^{-5}$) for labeled against unlabeled lysines, and d = 0.019 (q = 0.71) between the two probes. |
| Figure 7 | A | The first version of this comparison measured union positions at Cβ and measured lysines at Nζ, and returned a separation of 2.256 against a control of 0.330, a 6.84-fold ratio that is an artifact of the mismatch. The three legitimate anchorings, both sides at Cβ, both modeled at Nζ, and both at Cβ under a symmetric convention, give 0.571, 0.535 and 0.583 against controls of 0.410, 0.331 and 0.407. |

### Extended Results

*Analyses that support the main-text figures and are reported here in full.*

#### Supporting main-text Figure 2. The class is mostly missed by a search built to find it.

The ten driver hotspots among the 617 covered sites, all of them on the missed side: BRAF V600K (IGR1, MDST8), NRAS Q61K (HT1080, SKNAS), PIK3CA E542K (CAL51), PTEN I67K (LS180), and TP53 M133K (HEL), R280K (MDAMB231), E258K (MEWO) and E285K (RPMI8226).

#### Supporting main-text Figure 3. Reported error rates do not hold for variant peptides.

That rate is measurable on the acquired-lysine class alone, and measured there it is not lower. A rate estimated on every variant peptide and then applied to one kind of substitution is an inference across populations, and it does not have to hold, since a substitution to lysine creates a tryptic cleavage site and the peptide the search must find is not the peptide it would have looked for. The restriction is legitimate here because the entrapment design is pairwise, so keeping only lysine-creating substitutions preserves the one-to-one match instead of slicing an unmatched comparison (note N11). Restricted that way the class runs higher than the rest of the variant class, and although the counts cannot resolve the difference, the direction is the one that matters, because it means borrowing the whole-class figure is not the conservative choice. Past the margin the class contributes no entrapment peptides at all, which bounds its false-discovery proportion by the rule of three rather than measuring it at zero. Seven entrapment peptides across 42 plexes is the entire supply of counted errors, so no single class of substitution can be resolved to better than a few percent on its own.

Two denominators are in use and they are not interchangeable. Figure 3B counts distinct peptides, one best spectrum each, which is the unit the entrapment pairing is defined over; Extended Results counts identifications. Both are correct. Decoys do not see any of this, which is why the margin is applied rather than the reported q-value trusted.

The surviving identifications are ordinary spectra, not marginal ones. Judged on properties of the spectrum alone, with no reference to the search engine or the database, the acquired-lysine hit spectra are at the background peak count and a little above its signal-to-noise (Figure 3D). The one feature that invites suspicion is a tail at plus one dalton, the signature of the first isotope being selected instead of the monoisotopic peak, and it occurs at the same rate in variant and in background spectra, so it is instrument behavior and not a property of variant peptides.

The entrapment audit restricted to the acquired-lysine class: 66 peptides compete with five entrapment at margin 0, 7.04% against 4.58% for the rest of the variant class, a difference the counts cannot resolve (Fisher exact, odds ratio 1.58, $p = 0.38$). Past the margin the class contributes zero entrapment against the 38 variant peptides quoted in the main text, which is what bounds its proportion at 7.9% by the rule of three.

#### Supporting main-text Figure 4. Reactivity and ligandability separate only where the structure is trustworthy.

Packing features, feature by feature, reactivity against ligandability under the gate: crowding runs +0.197 to −0.042 in reactivity while holding at +0.143 to +0.100 in ligandability; atom density +0.244 to +0.003 against +0.157 to +0.109; polar contacts +0.183 to −0.002 against +0.185 to +0.197. The feature's own standard deviation holds or widens in each case (crowding

4.56 to 4.70, polar 1.66 to 1.81, aromatic 1.08 to 1.25). Exposure and distance to the centroid are the two exceptions, losing 12% and 29% of their spread.

Charge effects under the gate, for comparison with the shape features: anionic contacts strengthen from −0.307 to −0.454 and salt bridges from −0.221 to −0.327, while the ligandability arms run −0.210 to −0.336 and −0.183 to −0.286, so both sides move in step.

**Supporting main-text Figure 5. Ligandability reproduces at the site level and resists prediction from site structure or compound properties.**

**The ceiling is not an artifact of disordered sites.** Computed within five bands of AlphaFold confidence at the lysine [S29,S30], the lysine-targeting compound screen ceiling runs from 0.672 to 0.734 with no trend across them, and reads 0.734 on the 8,436 sites at pLDDT ≥ 90, where the structure is most trustworthy and the shortfall is therefore largest. The other two screens are higher in every populated band. The bands are disordered (< 50), low (50 to 70), confident (70 to 90), very high (≥ 90) and no structure; the lysine-targeting compound screen and CyMPSA populate all five, the meroterpenoids two.

**Nor is the shortfall a feature-count problem.** The fourteen features do not carry fourteen independent properties: the first principal component alone accounts for 48.9% of their variance, and five components clear 80%. Burial, crowding, hydrophobic packing and predicted $pK_a$ [S31,S32] are facets of one situation, a lysine in a pocket, measured several ways, which is also why the composite improves on its best single feature, distance to the protein centroid, by only +0.019 (Figure S6B). More features of this kind would not close the gap; a different kind of information would be needed.

**Two enantiomer pairs have enough shared coverage to compare.** Guadial B engages nineteen sites with both enantiomers against 0.1 expected by chance and eight with a single one; euglobal G2 engages six with both against 0.0 expected and one with a single one. Guadial B was measured at 7,043 sites and euglobal G2 at 7,560. The dominant signal is agreement: two molecules of identical constitution and opposite three-dimensional shape overwhelmingly engage the same lysines. The minority engaged by a single enantiomer cannot be intrinsic reactivity, since the two enantiomers present the same chemistry, but with n = 2 pairs that is a bounded observation, not a selectivity claim.

**The preference is a gradient, not a threshold effect.** Across 958,108 site × compound observations binned by occupancy, every one of six site features, taken as standardized shifts from the 0 to 10% tier, moves monotonically and accelerates through the 60% line. The tiers are very unequal (734,394 observations in the 0–10% bin against 648 at ≥ 90%), so the direction is secure while the acceleration at the right-hand end rests on the two smallest bins. Three threshold-free continuous measures agree (Figure S7C), so nothing here depends on a 60% cut existing at all.

**The pooled positives are provenance, not chemistry.** Molecular weight all but names the screen a compound came from (medians 306 Da for the lysine-targeting compound screen, 386 for the meroterpenoids and 480 for CyMPSA, Kruskal–Wallis $p = 1.3 \times 10^{-9}$), and each screen is a different cell line in a different sample state. More directly, the descriptors that separate the three screens are the descriptors that produced the pooled false positives: across ten descriptors against fifteen site features the two rank orders correlate at Spearman $\rho = 0.69$, $p = 0.027$. TPSA is the exception, fourth by effect size and the source of no pooled false positive at all. Pooling therefore lets provenance masquerade as chemistry, which is why the comparison is run within each screen rather than on the pool.

**The breadth gradient of Figure 5F does not depend on where the occupancy cut is drawn.** Repeating it here on the site features, the most broadly engaged sites differ from a never-engaged site by 0.48 SD on the average feature, averaged over fifteen features, the fourteen site features together with pocket occupancy, against 0.09 for shuffled labels, and the gradient holds at five cuts from 40% to 80%. Two cuts are not strictly monotonic, 40% and 80%. At 80% the groups are the smallest of any cut and one step reverses, between the groups of 53 and 36 sites; at 40% the groups are the largest of any cut and the first two are level to three decimals, so small groups account for the first exception and not for the second (Figure S7E).

**Supporting main-text Figure 6. The probe is not a neutral reporter: reagent charge steers labeling.**

Site environment of labeled lysines, the effect sizes behind the nucleophilicity reading, measured at pLDDT ≥ 70: fewer acidic neighbors (d = −0.46), more cationic ones (+0.40), fewer salt bridges (−0.32) and lower predicted $pK_a$ (−0.30).

**Supporting main-text Figure 7. Acquired lysines are structurally ordinary.**

Boundary independence, the numbers behind the threshold-free reading: acquired lysines fall inside the 50, 75, 90 and 95% engaged regions at 42, 69, 82 and 88% under one anchoring and 45, 67, 83 and 89% under another, against a labeled-but-unengaged control running 47, 73, 89 and 95% and 50, 78, 92 and 96%.

**From the Discussion: the prospective screen this paper does not run.**

Which cells carry the class, and what gating does to it. The four cell lines of the three chemoproteomic screens carry 56 unique acquired lysines between them, against a median of 10 and a maximum of 623 across 1,954 DepMap models: MDA-MB-231 24, MDA-MB-436 19, HeLa seven and Ramos six (Figure S10B). Eleven of the 56 fall on a protein the matching screen probe-labels at one or more lysines, and one of them, SF3A3, already yields a lysine engaged above 60% (Figure S10E). The lysine measured is not the acquired one and usually sits far from it, a median of 271 residues away, so what the eleven establish is that the protein is reachable by a probe rather than that the acquired site is. A re-search of those three screens is worth running, and it would test eleven sites rather than a class.

Burden by lineage runs skin, uterus, bowel and lung at the top and eye second from last; at subtype resolution cutaneous melanoma carries a median of 37 against ocular melanoma's three, which is the cutaneous-versus-uveal contrast recovered in cell lines rather than tumors (Figure S10A). The asymmetry that makes the class is visible one cell line at a time: across 1,954 models 39,390 lysines are gained against 25,531 lost, and the ratio runs from 6.4 in skin to 1.3 in esophagus and stomach (Figure S10B). A quarter of models are net-negative. Burden is not the same as opportunity either. Passing an acquired lysine through three gates, the gene expressed in that line, the protein quantified in that line's CCLE proteome, and the gene a CRISPR dependency, leaves 46%, then 31%, then 6% of 2,236 sites (Figure S10C). Six percent is not a small number in absolute terms: 132 sites in 27 cell lines are on a protein that is present, detectable and required. The gates also expose a gap in the reference data: only 308 of 1,954 models carry expression, proteomic and CRISPR data together, and none of the ten highest net-gain lines does. The cells that carry most of the class are the least characterized.

The sites that recur are the sites this paper already measured. Of the 39,390 unique acquired lysines, 248 carry a hotspot or driver flag, and ranked by how many cell lines carry them the leaders are NRAS Q61K in 36, PIK3CA E545K in 33, PIK3CA E542K in 15, XPO1 E571K in nine, TP53 E286K in seven, TP53 E285K in six and AKT1 E17K in five (Figure S10D). Four of the six sites whose structures are measured in the structural comparison are on that list: three of them are its three most recurrent, and AKT1 E17K is seventh, which was not how they were chosen. BRAF V600K, in three models, is an acquired lysine on the most-drugged oncogene in melanoma. Four driver-flagged sites clear all three gates in the models examined, in five site-in-line instances: MTOR E1799K in HEC59, PIK3CA E545K in MCF7, BRAF V600K in MDST8 and in IGR1, and PIK3CD E1021K in REH. PIK3CD already carries an approved drug against another site. What is independently known about these sites is collected in Table S1, which is provenance rather than a survey. Twelve of the thirteen are carried by TCGA patients as well as by cell lines (PIK3CA E545K by 263 of the 10,224 participants in the MC3 call set, E542K by 169, NRAS Q61K by 68, AKT1 E17K by 54), and nine carry a functional consequence measured by someone else, among them impaired GTP hydrolysis at NRAS Q61K and accelerated leukemogenesis at XPO1 E571K. The thirteenth is instructive rather than absent: BRAF V600K returns no MC3 patients because valine reaches lysine only through two adjacent nucleotide changes and that call set does not merge adjacent variants, so the site is invisible to it by construction while DepMap reports it in three lines. These are the prospective screen's first targets, and naming them, with the gaps marked, is what a catalog is for.

A choice of cells follows. Lysine chemoproteomics has been run in lines that barely carry the class, while the chemoproteogenomic work on acquired cysteine ran in high-burden lines [S33]. Ranked on the sites a screen could actually reach, acquired lysines on a gene the line expresses, the leaders are MAMEL46 with 248, MCC13 with 178 and HEC251 with 176, against three in HeLa. A prospective screen in any of them would put the same chemistry against 59 to 83 times the reachable sites.

### Supplementary notes

#### N1. Site validation: two-route agreement, the peptide correction and the marker rescue.

Every chemoproteomic site used in Figures 4 to 6 was assigned by two independent routes and the two assignments reconciled before any analysis ran.

**Of the 45,552 usable sites, 44,736 agree between the two routes, 98.21%.** The remaining 816 were recovered rather than discarded: 470 were corrected from the peptide and 346 resolved by the marked-lysine method below. The verdict is retained as a column, so any analysis here can be restricted to the two-route agreement set alone.

**The marker was validated before it was used.** On 4,000 sites whose position was already confirmed two other ways and which also carried a marked peptide, the marker pointed to the same residue in 3,999 of them, 99.97%, and disagreed once.

**What the rescue recovered.** 529 rows were unusable before it: 442 ambiguous, 49 matching several positions and 38 with no positional evidence. The marker resolved 346 of them, 281 in CyMPSA and 65 in the lysine-targeting compound screen; 121 had neither a marker nor a structure and 62 sat on a peptide that was not unique. All 346 came from the ambiguous class, in which the peptide matched the sequence exactly once, carried more than one lysine, and the number the source reported named none of them.

**How far the resolved positions moved.** No resolved position coincides with the number originally reported, and none can, for the reason just given. The signed displacement has a median of +1 residue, but 166 displacements are negative and 180 positive, so that figure is cancellation rather than agreement and is not the one to quote. The magnitude is: the resolved lysine sits a median of 34 residues from the reported number, interquartile range 12 to 111, maximum 3,234. As raw geography, the reported number falls inside the matched peptide but on a residue that is not a lysine in 36 rows, elsewhere in the same protein in 276, and outside the protein's modeled residue range altogether in 34; beta-actin, 375 residues long, carries a reported position of 913.

**Most of that displacement is a numbering frame, not a wrong site.** Sorting the 346 by the strongest explanation available for each: in 64 the reported number lands exactly on the marked lysine in an annotated alternative isoform of the same protein, so the source coordinate was never wrong, only written in another frame; in 31 it falls inside the matched peptide on a non-lysine, a median of one residue away, and these are almost the whole of the 21 rows that sit within a single residue; in 27 it falls outside the protein entirely; the remaining 224 fall elsewhere in the protein with no isoform that accounts for them. Of the 52 proteins contributing more than one rescued row, 38, covering 127 rows, are displaced by one identical signed offset throughout, which is what a frame difference looks like and not what a randomly misassigned site looks like. Capicua is displaced by +909 residues in all three of its rows, and its shorter annotated isoform is exactly 909 residues shorter than the modeled one.

**Why a wrong reported number cannot reach the analysis.** The marker never reads the reported number as a coordinate. It locates the peptide in the AlphaFold sequence, requires that the peptide occur exactly once, and takes the lysine at the marked offset; the reported number serves only to join back to the source row. Geometry is therefore computed at a residue fixed by the peptide, not by the number. These 346 rows are 0.76% of the 45,552 usable sites, they retain the verdict MARKER, and the engagement-rate model carries the verdict as a covariate, so every result can be reproduced with them excluded. step6c_audit_shift.py regenerates every figure in these two paragraphs from features_final.csv and UniProt.

**A denominator warning, because both numbers are real.** 44,736 of the 45,735 raw feature rows is 97.82%, and 44,736 of the 45,552 usable sites is 98.21%. The usable-site denominator is the one this work quotes, since rows that never became usable sites are not sites the two routes could have agreed about.

**N2. Occupancy scale construction across the three datasets.**

Occupancy is the share of probe labeling that a compound competes away. The three screens measure it on two different instruments, and the scale was built so that one number means the same thing in all of them.

For the TMT competition datasets, CyMPSA and the meroterpenoids, occupancy is 100 × (1 − mean(compound channels) / mean(DMSO channels)). For the isoTOP-ABPP ratio dataset, the lysine-targeting compound screen, where R is the DMSO-to-compound ratio, occupancy is 100 × (1 − 1/R). The published criterion of the lysine-targeting compound screen of liganded at a ratio of 4 or more is therefore a 75% occupancy cut, not the 60% cut the other two screens use; a ratio of 2.5 is 60%. This work uses 60% throughout, and Figure S7C shows the Figure 5 result does not depend on a 60% cut existing at all.

**Three scale caveats travel with every row as explicit columns rather than as prose.** The number of conditions per site differs enormously between screens, a median of 41 in the lysine-targeting compound screen against 5 to 15 for the others, so a per-site maximum taken over more conditions is larger for purely statistical reasons; n_conditions is a covariate in the models rather than an afterthought. isoTOP quantifies isotopologue pairs at MS1 while the TMTpro screens use isobaric reporters, which compress ratios toward one and therefore understate occupancy. A residual platform offset survives matching on n_conditions, so datasets are held out as cross-validation folds and raw occupancies are never pooled as though they were one measurement.

Every row also carries a single-target-residue flag, set where the peptide contains exactly one residue of the probed type, so any analysis here can be repeated on the unambiguous subset as a sensitivity check.

**N3. Screening depth by dataset, and why every ligandability estimate here is a lower bound.**

A site that no compound engaged may not be ligandable, or may simply not have been tried hard enough. The two are separated by depth, and depth is wildly uneven across the three screens.

**Depth per site**, the denominator behind every negative: CyMPSA and the meroterpenoids test every site at exactly 5 compounds. The lysine-targeting compound screen runs from 1 to 328, median 41, 90th percentile 214. Its depth is counted in compound-by-concentration conditions, because 181 of the 186 compounds in its library were profiled at one to three concentrations each, giving 328 conditions. A negative from 5 compounds and a negative from 41 are not the same statement, which is why n_conditions is a covariate throughout rather than a footnote.

**Discovery curves**, compounds added in random order over 30 permutations, median reported. CyMPSA reaches 77 engaged sites at 1 compound, 307 at 5 and 721 at 15, with the last compound still adding 19 sites, 2.6% of the total. The meroterpenoids reach 27, 95 and 143 at 1, 5 and 10 compounds, the last still adding 5, 3.5%. The lysine-targeting compound screen reaches 5, 51, 166, 1,611 and 2,465 at 1, 5, 10, 162 and 324 conditions, and its last condition adds 1 site, 0.0%.

**Two of the three screens are plainly unsaturated and the third is not evidence of saturation.** CyMPSA and the meroterpenoids are still gaining sites at their last compound, so for them not engaged means not yet engaged by what was tried. The lysine-targeting compound screen's curve has flattened, but it flattened over 328 conditions applied to a population whose per-site depth runs from 1 upward, so the flattening describes the compounds that were tried rather than the sites that remain. Every ligandability figure in this work is therefore a lower bound.

**Per-compound reach**, for scale: CyMPSA has 15 compounds engaging a median of 64 sites (16 to 304), the meroterpenoids 10 compounds engaging a median of 26 (10 to 47), and the lysine-targeting compound screen 328 conditions engaging a median of 6 (0 to 467).

**N4. Cross-screen concordance of site ligandability, and the compound-treatment condition.**

Compounds were applied to intact cells in the CyMPSA and meroterpenoid screens and to cell lysate in the lysine-targeting compound screen. In all three, the probe was applied to lysate after compound treatment, so the readout is identical and the screens differ only in where the compound encountered the proteome.

No compound in these data was assayed in both conditions, so no controlled comparison of the condition is available. What can be measured is whether ligandability agrees between screens. Among sites assayed in both members of a pair, ligandability in one screen predicts ligandability in the other in every pair: the lysine-targeting compound and CyMPSA screens, 6,811 shared sites, odds ratio 1.60, Fisher exact $p = 5.5 \times 10^{-5}$; the lysine-targeting compound and meroterpenoid screens, 3,712 shared sites, 3.12, $p = 2.4 \times 10^{-6}$; CyMPSA and the meroterpenoids, 8,211 shared sites, 6.05, $p = 2.0 \times 10^{-11}$.

The largest odds ratio belongs to the pair that shares a condition. That ordering is consistent with a condition effect and equally consistent with library similarity (CyMPSA and the meroterpenoids are 15 and 10 elaborated natural products, the third is 186 lysine-targeting compounds), and across three screens the two explanations are collinear. No condition effect is claimed from the ordering, and none is excluded. The ligandable base rates carry the same warning: the lysine-targeting compound screen calls 17.5% of its sites ligandable against 2.9% and 1.5%, which is compound promiscuity and compound count rather than condition.

These odds ratios are therefore a lower bound on site-intrinsic transfer, measured across condition, cell line and compound library at once, and not an estimate of any one of them.

Lysate-phase measurement is the standard on which ligandability atlases are built. The chemoproteogenomic platform closest to this work profiles cell lysate and makes its druggability claim from it [S33]. DrugMap compiles cysteine ligandability across 416 cancer cell lines from lysate, and reports ligandable newly encoded cysteines among them [S34]. SLC-ABPP reports that the same workflow runs on intact cells, lysates and tissue with minimal change and without loss of quantitative cysteine depth [S35].

Computed by step50_condition_concordance.py; a site is ligandable in a screen at ≥ 60% occupancy by at least one of that screen's compounds.

**N5. Pocket membership survives the confidence gate, and the pocket axis is not the charge axis.**

Figure 4 shows every shape feature collapsing once the structure is required to be well-modeled while the charge features strengthen. Pocket membership is a shape feature by any ordinary reading, so it should collapse too. It does not.

**Overall.** Of 30,805 probe-labeled lysines 44.7% sit in a pocket, median distance 5.53 Å, against 39.8% and 6.47 Å for the 214,879 unlabeled ones.

**Across the gate**, within protein, reported as effect over the feature's own standard deviation so that a shrinking effect can be checked against a shrinking spread. Being in a pocket runs 0.187, 0.111, 0.107 and 0.111 at gates of none, 50, 70 and 90, on a standard deviation flat at 0.49 to 0.50. Pocket volume strengthens, 0.047 to 0.064. Distance to the nearest pocket collapses, −0.316 to −0.066, but its standard deviation collapses with it, 8.65 to 4.14 Å, so that decline is spread rather than signal. Pocket score and druggability score are flat and never significant.

**Read against Figure 4**, where crowding reverses from +0.197 to −0.042, atom count collapses from +0.244 to +0.003 and polar contacts reverse from +0.183 to −0.002 on flat spreads while anionic contacts strengthen from −0.307 to −0.454, pocket membership behaves like the charge features rather than like the shape features. Reactive lysines are at cavities, and probe capture is already enriching for sites a molecule could occupy.

**The pocket axis is not the charge axis.** In a principal-component analysis over 167,035 lysines at pLDDT ≥ 70 and 17 features, PC1 carries 40.1% of the variance and is a burial axis: side-chain solvent accessibility −0.362, relative accessibility −0.350, Nζ accessibility −0.338, atom count +0.369 and crowding +0.368. The pocket terms ride on it, distance −0.135 and in-pocket +0.118, while the charge terms are at zero, net charge −0.032 and predicted $pK_a$ −0.020. PC2 carries 13.9% and is the charge axis: predicted $pK_a$ −0.471, anionic contacts −0.406, salt bridges −0.384, with the pocket terms back at zero, distance −0.066 and volume +0.024. The two properties travel on different components, so the pocket result is not the charge result under another name.

**N6. Promiscuity control: the breadth gradient sharpens when promiscuous compounds are excluded.**

The objection is that sites engaged by many compounds are simply the sites a few very promiscuous compounds hit, so the breadth gradient would measure the compounds rather than the lysines.

**The library is genuinely uneven.** Across 353 compounds, counting each concentration of a compound in the lysine-targeting compound screen as a separate entry (15 + 10 + 328), the median engages 7 sites, the 90th percentile 58 and the most promiscuous 467. The ten most promiscuous compounds account for 34.0% of all site-compound engagements. The test is therefore to delete them and rerun.

**The gradient survives every deletion and steepens.** On the 3,513 lysines measured in all three datasets, mean distance from a never-engaged site rises monotonically with breadth in every run. With all compounds the tiers run 0.264, 0.627, 0.825, 1.129 and 2.616 against nulls of 0.177 to 0.546. Excluding the five most promiscuous compounds they run 0.247, 0.783, 0.835, 1.378 and 2.853; excluding the ten, 0.247, 0.921, 1.030, 1.199 and 3.201. The most broadly engaged tier moves from 2.616 to 3.201 as the promiscuous compounds come out, which is the opposite of what the objection predicts.

**What weakens instead is the singly-engaged tier**, from $p = 0.058$ with all compounds to $p = 0.185$ with ten removed, on a falling n. Sites engaged by exactly one compound are the tier promiscuity was most likely to manufacture, and they are the tier that does not survive. The gradient is carried by the sites that several distinct compounds reach.

**N7. Compound descriptors and 3D shape, the nulls: the full descriptor list behind Figure S7, which is what makes Figure S7F's bound checkable.**

Figure 5 is a null result over the descriptors that were tested, so the descriptors have to be listed and their blind spots measured rather than asserted.

**The ten tested descriptors** are molecular weight, logP, topological polar surface area, hydrogen-bond donors, hydrogen-bond acceptors, ring count, aromatic ring count, rotatable bonds, heavy-atom count and the fraction of sp3 carbons. The shape terms are NPR1 and NPR2, ratios of principal moments of inertia.

**Charge: the claim is a contrast, not an absence.** Over the 212 compounds with a parsable structure, net charge at pH 7.4 is 0 for 171, +1 for 35, +2 for 2, +3 for 2 and −1 for 2. Two descriptors are weakly but significantly correlated with that charge, ring count at $|\rho| = 0.193$ ($p = 0.0047$) and hydrogen-bond acceptors at 0.186 ($p = 0.0066$). The largest charge correlation in the set is therefore 0.193 against a median size correlation of 0.552, a ratio of 0.35. The defensible statement is that the tested descriptors carry about a third as much charge information as size information, not that none of them encodes charge.

**Handedness: the shape terms are blind to it by construction.** Reflecting a conformer changes NPR1 and NPR2 by exactly zero. Re-embedding an enantiomer changes them by up to 0.281 and 0.251 across 25 chiral compounds, against a between-compound standard deviation of 0.123 and 0.092. That difference is conformer sampling rather than chirality, and an earlier check measured it and would have been read as a chirality effect. The descriptors respond to handedness not at all, and their own sampling noise is already a third of the spread they resolve between compounds.

This is what bounds Figure 5. Charge and chirality are the two axes the descriptor set sees least well, and Figure 6 and Extended Results are where each is shown to matter.

### N8. Crystallographic validation candidates.

Figure 7D asks whether an acquired lysine moves the structure around it more than two crystals of the same wild-type protein differ from each other. Superposition is on residues beyond 12 Å of the site and the reported displacement is over Cα atoms within 8 Å of it, so wild type against wild type is the noise floor rather than zero.

**Six sites were defined in advance and all six were measured; none was excluded.** They are PTPN11 76, TP53 280, AKT1 17, PIK3CA 542, PIK3CA 545 and NRAS 61. Where a substitution series exists it was carried through: PTPN11 76 has E76K, E76A, E76D and E76Q against five wild-type structures, which is what makes the 3.53 of Figure 7D interpretable against 3.21 for other substitutions at the same position.

**The NRAS floor rests on a single pair, and deliberately so.** The wild-type set was restricted to one nucleotide state so that every pair is state-matched. Residue 61 sits in switch II, and an earlier run that mixed GDP and GTP structures inflated that floor to 4.25 Å and flattered the mutant. The panel is annotated with that n of 1 rather than leaving it to be inferred. Two of the six comparisons also cross a method boundary the others do not: the PIK3CA mutant structures 8GUA and 8GUD are cryo-EM while their wild-type partners are crystal structures.

**Global rearrangement is reported separately from the local question.** Eleven pairs align locally but not globally, ten of them involving 6CRF, which is deposited under its own title as SHP2 E76K in the open conformation. E76K is an activating mutation that releases the autoinhibited state, so a large whole-protein difference alongside an ordinary local one is that effect. It argues for the figure rather than against it: the site itself is unremarkable even when the protein as a whole has moved.

**The limit, stated because it is real.** Crystal structures differ for crystallographic reasons, among them space group, resolution, bound ligands and temperature, and this measure cannot separate those from the effect of the mutation. That is exactly why wild type against wild type is the comparator.

### N9. PgxSAVy answers a different question from the one this work asks.

PgxSAVy is the obvious tool to reach for here, and it does not do the job this work needs, which is worth saying plainly rather than leaving as an omission.

**What it does.** PgxSAVy rescores individual variant peptide-spectrum matches and combines match quality, variant-event and search-result features into a variant ambiguity score, then labels each match confident, semi-confident or doubtful. On a reanalysis of roughly 2.8 million spectra it labeled 3,028 of 12,705 variant matches confident, 1,409 semi-confident and 8,268 doubtful.

**Why that is not what is needed.** A per-match confidence label is not a false-discovery proportion for a class. The question here is what fraction of the variant identifications in a defined population are wrong, and a classifier answers it only if its calibration on this data is known, which nothing establishes. Burden-matched entrapment measures the proportion directly, on this search and this database, which is why Figure 3 uses it.

**Two further mismatches.** PgxSAVy evaluates matches one at a time and does not conflate them into peptides, whereas the unit that matters here is the distinct peptide and the identification, kept apart throughout Figure 3. And its scoring was developed and calibrated on a different search pipeline, so its thresholds are not transferable to this one without a recalibration nobody has done.

The two approaches are complementary rather than competing. PgxSAVy would improve confidence in individual variant matches; it would not tell this work what its realized error rate is.

**N10. Entrapment prior art: the survey this design answers to. Cited, not claimed.**

Entrapment is not new and this work claims no novelty for it. What the recent literature establishes is that entrapment is easy to run and easy to run wrongly, which is why the design is spelled out in Figure 3 rather than named.

**The survey finding.** Wen et al. identify three validation methods in common use: one invalid, one that provides only a lower bound, and one valid but under-powered. Reviewing published entrapment experiments, they report that only three of the studies they tabulate use entrapment estimation correctly, and all three were data-dependent acquisition. The two common errors are treating the lower-bound estimator as though it established error control, and using the sample-entrapment method.

**Where this work sits.** The design here is burden-matched at a ratio of one, with each entrapment sequence built from its own target's residues so that the pair shares an exact precursor mass and an identical length. That is an upper-bound construction rather than a lower bound, it is applied to data-dependent acquisition, and the proportion is measured within the variant class rather than assumed from the whole search.

**What is cited and what is not.** The survey and its taxonomy are cited. No claim is made here that burden-matched entrapment is a new method; the contribution is its correct application to a variant population, and note N11 states the bound that follows for the acquired-lysine class specifically.

**N11. Class-restricted entrapment: the false-discovery proportion measured on the acquired-lysine class itself rather than borrowed from the whole variant class.**

Figure 3B measures the realized false-discovery proportion over every variant peptide in the search: 4.70% at margin 0, falling to 0.83% at margin ≥ 10. The main text then applies that number to acquired lysines. That is an inference from one population to a subset of it, and it is not automatic. A substitution to lysine introduces a tryptic cleavage site, so the peptide the search must match differs systematically from the peptide it would have looked for at the same position without the mutation; there is a mechanism by which the class could carry its own error rate.

**Why the subset can be taken.** The entrapment design is pairwise. Each target variant peptide is paired at build time with one entrapment peptide of matched burden (peptide_pair_index), and the pair inherits the same class tag from the same variant record: KGAIN for a substitution that creates a lysine, KNOVEL for a frameshift or novel stretch that contains one, NOK otherwise. Restricting to KGAIN therefore preserves the r = 1 burden match inside the subset; it is not a post-hoc slice of an unmatched comparison. step51_kgain_entrapment.py verifies this rather than assuming it, and reports zero class-mismatched pairs across all 42 plexes, over 817,808 target/entrapment pairs.

**Validation of the restriction.** Run without the class filter the script reproduces Figure 3B exactly: 1,419 variant peptides against 70 entrapment at margin 0 (4.70%) and 832 against 7 at margin ≥ 10 (0.83%), which is the check that it is slicing the published population and not a different one. The unit throughout is one best spectrum per distinct peptide sequence per plex, and the estimator is $e / (t + e)$, both as in Figure 3B.

**Result.** At margin 0, 66 acquired-lysine peptides compete with five entrapment: 7.04% (Wilson 95% CI 3.05–15.45%), against 4.58% for the rest of the variant class (1,353 against 65). Fisher's exact test does not resolve the difference (odds ratio 1.58, $p = 0.38$). At margin ≥ 10 the class contributes zero entrapment peptides against 38 variant peptides, so its false-discovery proportion is bounded at 7.89% by the rule of three, or 9.18% by the Wilson interval, rather than measured at zero. KNOVEL, which is a different class and is not the subject of this paper, is worse at every margin, on counts too small to interpret (eight peptides at margin 0).

**What this settles and what it does not.** Seven entrapment peptides across 42 plexes at margin ≥ 10 is the entire supply of counted errors, so no single class of substitution can be resolved on its own to better than a few percent; the class-specific run cannot replace the whole-class estimate, and 0.83% remains the number quoted in the main text. What it establishes is the direction of the borrowing. Before the margin filter the acquired-lysine point estimate is higher than the rest of the class, not lower, so quoting the whole-class rate is not the conservative choice; after the margin filter the class contributes no counted errors at all, so the filtered set that the paper reports is not carrying a concealed excess. Both statements are reported here because only the second one flatters the result.

Computed by step51_kgain_entrapment.py from the per-plex varpair definitions and the mutation-aware search PSM tables; outputs in kgain_margins.csv and kgain_entrapment_sweep.csv.

**Supplemental References**


[S1] Nusinow DP, Szpyt J, Ghandi M, Rose CM, McDonald ER 3rd, Kalocsay M, et al. Quantitative proteomics of the Cancer Cell Line Encyclopedia. Cell. 2020;180(2):387–402.e16. doi:10.1016/j.cell.2019.12.023

[S2] Ryan EM, Norinskiy MA, Bracken AK, Lueders EE, Chen X, Fu Q, Anderson ET, Zhang S, Abbasov ME. Activity-based acylome profiling with N-(cyanomethyl)-N-(phenylsulfonyl)amides for targeted lysine acylation and post-translational control of protein function in cells. J Am Chem Soc. 2024;146(40):27622–27643. doi:10.1021/jacs.4c09073

[S3] Bracken AK, Gekko CE, Suss NO, Lueders EE, Cui Q, Fu Q, Lui ACW, Anderson ET, Zhang S, Abbasov ME. Biomimetic synthesis and chemical proteomics reveal the mechanism of action and functional targets of phloroglucinol meroterpenoids. J Am Chem Soc. 2024;146(4):2524–2548. doi:10.1021/jacs.3c10741

[S4] Abbasov ME, Kavanagh ME, Ichu T-A, Lazear MR, Tao Y, Crowley VM, am Ende CW, Hacker SM, Ho J, Dix MM, Suciu R, Hayward MM, Kiessling LL, Cravatt BF. A proteome-wide atlas of lysine-reactive chemistry. Nat Chem. 2021;13(11):1081–1092. doi:10.1038/s41557-021-00765-4

[S5] Der CJ, Finkel T, Cooper GM. Biological and biochemical properties of human rasH genes mutated at codon 61. Cell. 1986;44(1):167–176. doi:10.1016/0092-8674(86)90495-2

[S6] Buhrman G, Wink G, Mattos C. Transformation efficiency of RasQ61 mutants linked to structural features of the switch regions in the presence of Raf. Structure. 2007;15(12):1618–1629. doi:10.1016/j.str.2007.10.011

[S7] Smith MJ, Neel BG, Ikura M. NMR-based functional profiling of RASopathies and oncogenic RAS mutations. Proc Natl Acad Sci U S A. 2013;110(12):4574–4579. doi:10.1073/pnas.1218173110

[S8] Gebregiworgis T, Chan JY, Kuntz DA, Privé GG, Marshall CB, Ikura M. Crystal structure of NRAS Q61K with a ligand-induced pocket near switch II. Eur J Cell Biol. 2024;103(2):151414. doi:10.1016/j.ejcb.2024.151414

[S9] Ackermann J, Frutschi M, Kaloulis K, McKee T, Trumpp A, Beermann F. Metastasizing melanoma formation caused by expression of activated N-RasQ61K on an INK4a-deficient background. Cancer Res. 2005;65(10):4005–4011. doi:10.1158/0008-5472.CAN-04-2970

[S10] Li A, Ma Y, Jin M, Mason S, Mort RL, Blyth K, et al. Activated mutant NRasQ61K drives aberrant melanocyte signaling, survival, and invasiveness via a Rac1-dependent mechanism. J Invest Dermatol. 2012;132(11):2610–2621. doi:10.1038/jid.2012.186

[S11] Bader AG, Kang S, Vogt PK. Cancer-specific mutations in PIK3CA are oncogenic in vivo. Proc Natl Acad Sci U S A. 2006;103(5):1475–1479. doi:10.1073/pnas.0510857103

[S12] Hao Y, He B, Wu L, Li Y, Wang C, Wang T, et al. Nuclear translocation of p85β promotes tumorigenesis of PIK3CA helical domain mutant cancer. Nat Commun. 2022;13(1):1974. doi:10.1038/s41467-022-29585-x

[S13] Walker JS, Hing ZA, Harrington B, Baumhardt J, Ozer HG, Lehman A, et al. Recurrent XPO1 mutations alter pathogenesis of chronic lymphocytic leukemia. J Hematol Oncol. 2021;14(1):17. doi:10.1186/s13045-021-01032-2

[S14] Carpten JD, Faber AL, Horn C, Donoho GP, Briggs SL, Robbins CM, et al. A transforming mutation in the pleckstrin homology domain of AKT1 in cancer. Nature. 2007;448(7152):439–444. doi:10.1038/nature05933

[S15] Craven GB, Chu H, Sun JD, Carelli JD, Coyne B, Chen H, et al. Mutant-selective AKT inhibition through lysine targeting and neo-zinc chelation. Nature. 2025;637(8044):205–214. doi:10.1038/s41586-024-08176-4

[S16] Boudreau HE, Casterline BW, Burke DJ, Leto TL. Wild-type and mutant p53 differentially regulate NADPH oxidase 4 in TGF-β-mediated migration of human lung and breast epithelial cells. Br J Cancer. 2014;110(10):2569–2582. doi:10.1038/bjc.2014.165

[S17] Gomes AS, Trovão F, Andrade Pinheiro B, Freire F, Gomes S, Oliveira C, et al. The crystal structure of the R280K mutant of human p53 explains the loss of DNA binding. Int J Mol Sci. 2018;19(4):1184. doi:10.3390/ijms19041184

[S18] Xu D, Zheng H, Yu WM, Qu CK. Activating mutations in protein tyrosine phosphatase Ptpn11 (Shp2) enhance reactive oxygen species production that contributes to myeloproliferative disorder. PLoS One. 2013;8(5):e63152. doi:10.1371/journal.pone.0063152

[S19] Rubinstein JC, Sznol M, Pavlick AC, Ariyan S, Cheng E, Bacchiocchi A, et al. Incidence of the V600K mutation among melanoma patients with BRAF mutations, and potential therapeutic response to the specific BRAF inhibitor PLX4032. J Transl Med. 2010;8:67. doi:10.1186/1479-5876-8-67

[S20] Menzies AM, Haydu LE, Visintin L, Carlino MS, Howle JR, Thompson JF, et al. Distinguishing clinicopathologic features of patients with V600E and V600K BRAF-mutant metastatic melanoma. Clin Cancer Res. 2012;18(12):3242–3249. doi:10.1158/1078-0432.CCR-12-0052

[S21] Pires da Silva I, Wang KYX, Wilmott JS, Holst J, Carlino MS, Park JJ, et al. Distinct molecular profiles and immunotherapy treatment outcomes of V600E and V600K BRAF-mutant melanoma. Clin Cancer Res. 2019;25(4):1272–1279. doi:10.1158/1078-0432.CCR-18-1680

[S22] Dunsche L, Ivanisenko N, Riemann S, Schindler S, Beissert S, Angeli C, et al. A cytosolic mutp53(E285K) variant confers chemoresistance of malignant melanoma. Cell Death Dis. 2023;14(12):831. doi:10.1038/s41419-023-06360-4

[S23] Christgen M, Noskowicz M, Heil C, Schipper E, Christgen H, Geffers R, et al. IPH-926 lobular breast cancer cells harbor a p53 mutant with temperature-sensitive functional activity and allow for profiling of p53-responsive genes. Lab Invest. 2012;92(11):1635–1647. doi:10.1038/labinvest.2012.126

[S24] Kato S, Han S-Y, Liu W, Otsuka K, Shibata H, Kanamaru R, Ishioka C. Understanding the function–structure and function–mutation relationships of p53 tumor suppressor protein by high-resolution missense mutation analysis. Proc Natl Acad Sci U S A. 2003;100(14):8424–8429. doi:10.1073/pnas.1431692100

[S25] Grabiner BC, Nardi V, Birsoy K, Possemato R, Shen K, Sinha S, et al. A diverse array of cancer-associated MTOR mutations are hyperactivating and can predict rapamycin sensitivity. Cancer Discov. 2014;4(5):554–563. doi:10.1158/2159-8290.CD-13-0929

[S26] Angulo I, Vadas O, Garçon F, Banham-Hall E, Plagnol V, Leahy TR, et al. Phosphoinositide 3-kinase δ gene mutation predisposes to respiratory infection and airway damage. Science. 2013;342(6160):866–871. doi:10.1126/science.1243292

[S27] Lucas CL, Kuehn HS, Zhao F, Niemela JE, Deenick EK, Palendira U, et al. Dominant-activating germline mutations in the gene encoding the PI(3)K catalytic subunit p110δ result in T cell senescence and human immunodeficiency. Nat Immunol. 2014;15(1):88–97. doi:10.1038/ni.2771

[S28] Penther D, Viailly PJ, Latour S, Ruminy P, Bobee V, Kaplan B, et al. A recurrent clonally distinct Burkitt lymphoma case highlights genetic key events contributing to oncogenesis. Genes Chromosomes Cancer. 2019;58(9):595–601. doi:10.1002/gcc.22743

[S29] Jumper J, Evans R, Pritzel A, Green T, Figurnov M, Ronneberger O, et al. Highly accurate protein structure prediction with AlphaFold. Nature. 2021;596(7873):583–589. doi:10.1038/s41586-021-03819-2

[S30] Varadi M, Anyango S, Deshpande M, Nair S, Natassia C, Yordanova G, et al. AlphaFold Protein Structure Database: massively expanding the structural coverage of protein-sequence space with high-accuracy models. Nucleic Acids Res. 2022;50(D1):D439–D444. doi:10.1093/nar/gkab1061

[S31] Tien MZ, Meyer AG, Sydykova DK, Spielman SJ, Wilke CO. Maximum allowed solvent accessibilities of residues in proteins. PLoS One. 2013;8(11):e80635. doi:10.1371/journal.pone.0080635

[S32] Olsson MHM, Søndergaard CR, Rostkowski M, Jensen JH. PROPKA3: consistent treatment of internal and surface residues in empirical $pK_a$ predictions. J Chem Theory Comput. 2011;7(2):525–537. doi:10.1021/ct100578z

[S33] Desai H, Andrews KH, Bergersen KV, Ofori S, Yu F, Shikwana F, et al. Chemoproteogenomic stratification of the missense variant cysteinome. Nat Commun. 2024;15(1):9284. doi:10.1038/s41467-024-53520-x

[S34] Takahashi M, Chong HB, Zhang S, Yang T-Y, Lazarov MJ, Harry S, et al. DrugMap: a quantitative pan-cancer analysis of cysteine ligandability. Cell. 2024;187(10):2536–2556.e30. doi:10.1016/j.cell.2024.03.027

[S35] Kuljanin M, Mitchell DC, Schweppe DK, Gikandi AS, Nusinow DP, Bulloch NJ, Vinogradova EV, Wilson DL, Kool ET, Mancias JD, Cravatt BF, Gygi SP. Reimagining high-throughput profiling of reactive cysteines for cell-based screening of large electrophile libraries. Nat Biotechnol. 2021;39(5):630–641. doi:10.1038/s41587-020-00778-3
